# A structured coalescent model reveals deep ancestral structure shared by all modern humans

**DOI:** 10.1101/2024.03.24.586479

**Authors:** Trevor Cousins, Aylwyn Scally, Richard Durbin

## Abstract

Understanding the series of admixture events and population size history leading to modern humans is central to human evolutionary genetics. Using a coalescence-based hidden Markov model, we present evidence for an extended period of structure in the history of all modern humans, in which two ancestral populations that diverged ∼1.5 million years ago came together in an admixture event ∼300 thousand years ago, in a ratio of ∼80:20 percent. Immediately after their divergence, we detect a strong bottleneck in the major ancestral population. We inferred regions of the present-day genome derived from each ancestral population, finding that material from the minority correlates strongly with distance to coding sequence, suggesting it was deleterious against the majority background. Moreover, we found a strong correlation between regions of majority ancestry and human-Neanderthal or human-Denisovan divergence, suggesting the majority population was also ancestral to those archaic humans.

## 2 Introduction

Improvements in the technology to extract ancient DNA has enabled an increasingly detailed picture of human evolutionary genetics in the late Pleistocene and Holocene [1], which overwhelmingly suggests that in the last tens of thousands of years there has been repeated separation of populations which subsequently remix back, or admix. Further back in time, high-coverage genomes from Neanderthals and Denisovans strongly indicate gene flow from these archaic humans into non-Africans [2, 3, 4, 5], and more ancient gene flow from the ancestors of modern humans into the ancestors of Neanderthals [6, 7, 8]. Moreover, researchers have demonstrated that models which incorporate a contribution of ancestry within the last ∼100ky from an unknown archaic population better explain patterns of polymorphism in African populations than a model without such a contribution [9, 10, 11, 12, 13, 14, 15, 16]. However, the presence of more ancient admixture events is less clear [17].

The history of effective population size changes is another important quantity in understanding human evolutionary genetics [18]. The pairwise sequentially Markovian coalescent (PSMC) [19] was introduced to infer the coalescence rate over time, the inverse of which can be interpreted as the history of effective population sizes. PSMC assumes that a population evolved under panmixia, with random mating in the ancestral population at all times. Recent work, however, has shown that the ancestry of most populations is structured, in that individuals in most populations derive their ancestry from two or more populations that were separated for some time and then came back together in the process of admixture. In light of this work, PSMC’s assumption of an unstructured evolutionary history is potentially problematic. Moreover, theoretical analysis shows that for any panmictic model with changes in the effective population size, there necessarily exists a structured model that can generate exactly the same coalescence rate profile [20, 21].

Here, we specifically address this question of identifiability of structured ancestry from a pair of sequences. We demonstrate that the transition matrix of the PSMC hidden Markov model (HMM) has information that can distinguish a structured model from a panmictic model, even if they have matching coalescence rate profiles. We parameterise a model of ancestral population structure that leverages this information and introduce this in a new HMM called *cobraa*. Applying *cobraa* to real human data from the 1000 Genomes Project (1000GP) [22, 23, 24, 25, 26] and the Human Genome Diversity Project (HGDP) [27, 28], we show that a model of deep population structure, where modern humans are a result of two populations that diverged ∼1.5Mya admixing together ∼300kya in a ratio of ∼ 80:20%, better explains the data than does a continuously panmictic model. We use posterior decoding to infer regions of the modern human genome that are derived from each population, and find evidence for selection against the material from the population contributing the minority of ancestry. Moreover, we find a strong association between regions derived from the major ancestral population and and human-Neanderthal or human-Denisovan divergence, suggesting that the majority population was the ancestral population to Neanderthals and Denisovans.

## Results

### Identifiability of structured ancestry through the SMC transition matrix

We consider a pulse model of population structure, where there are two populations *A* and *B* which descend from a common, ancestral population (Figure 1a). Looking backwards in time, population *A* is panmictic until time *T*_1_ when a fraction *γ* of the lineages instantaneously move to a new population *B*; *A* and *B* remain in isolation until time *T*_2_ when all lineages merge into a panmictic, ancestral population. Population *A* may vary in time, but we enforce that *B* must be of constant size, and hence the parameters of this model are the population sizes in *A, N*_*A*_(*t*), the population size in *B* between *T*_1_ and *T*_2_, *N*_*B*_, the admixture fraction, *γ*, and the split and rejoin times, *T*_2_ and *T*_1_, respectively.

**Figure 1:**
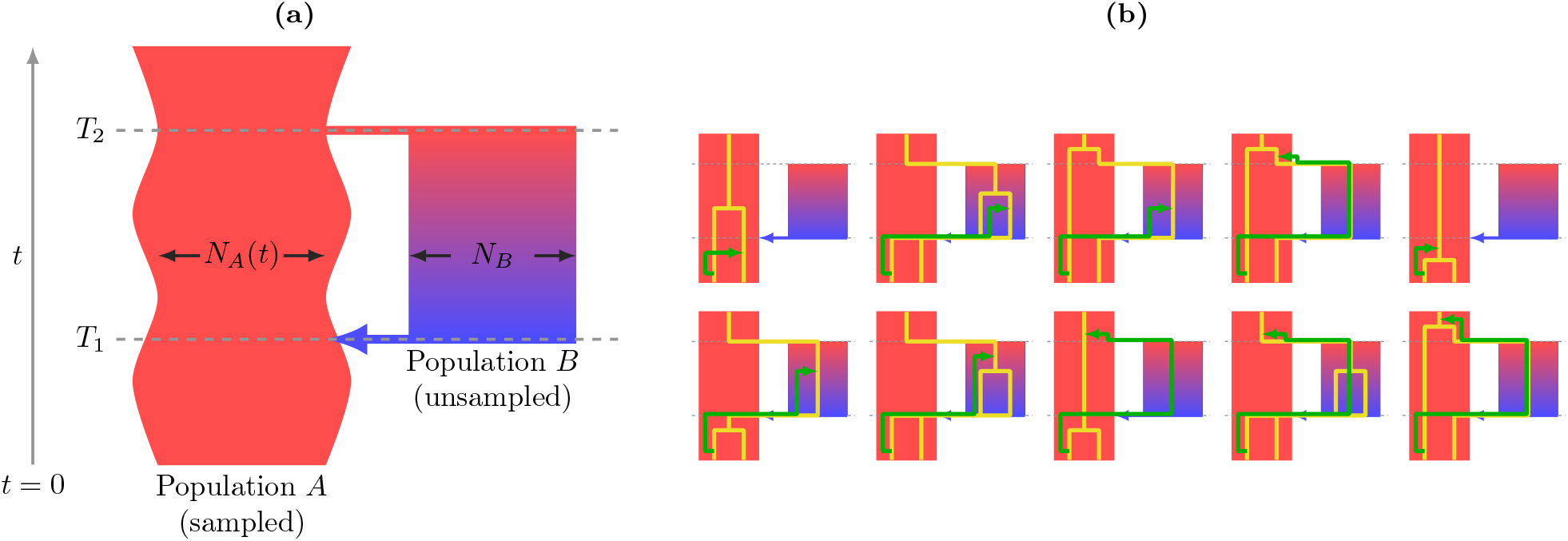
(**a)** Diagram of the structured model. Going forwards in time, an ancestral population splits into two populations, *A* and *B*, at time *T*_2_. These remain in isolation until time *T*_1_ when there is an admixture event. **(b)** We extend the SMC model to include the structure in (a). We consider two sampled lineages from population *A* at *t* = 0. The consequences of ancestral recombination are partitioned into 10 mutually exclusive cases, with an example from each illustrated. The gold lines indicate the two chromosomes sampled from the present at a particular locus and the green line indicates the “floating” lineage under the SMC model, which coalesces somewhere higher up on the tree in either population *A* or *B*.

The sequentially Markovian coalescent (SMC) was introduced to tractably approximate the coalescent with recombination [29], by modelling the coalescence time of a locus as conditionally dependent only on preceding locus [30, 31]. By assuming panmixia and discretising the SMC framework in the transition matrix of its hidden Markov model (HMM), the pairwise sequentially Markovian coalescent (PSMC) [19] infers the historical changes in the effective population size and the coalescence times across the genome. We extended the SMC framework to handle migration, so that changes in local ancestry can now be described according to the structured model, which involved partitioning the ancestral recombination process into 10 mutually exclusive cases according to possible migration or coalescence events. We give illustrative examples in Figure 1b and provide the mathematical details in Supplementary Text.

We now demonstrate that even if a structured and unstructured (i.e. panmictic) model have the same coalescence rate profile, the conditional distribution of neighbouring coalescence times distinguishes them.

Consider a structured model where *A* and *B* have a constant size and calculate its coalescence rate profile (blue line, Figure 2a); we can then construct an unstructured model with changes in its effective population size such that it has the same rate profile [20, 21] (orange line, Figure 2a). Now define the conditional distributions for the structured and unstructured model, and discretise these into transition matrices *Q*^*S*^ and *Q*^*U*^, respectively. The relative differences between these matrices *ξ* = (*Q*^*U*^ − *Q*^*S*^)*/Q*^*S*^ are clearly non-zero 2b. This is not a consequence of time discretisation, as the relative difference does not decrease as the number of time intervals increases (Figure S1). Moreover, the relative difference increases as does the admixture fraction or duration of population separation between *A* and *B* (Figure S1). Thus, even if a structured and unstructured model have indistinguishable coalescence rate profiles, the conditional distribution of neighbouring coalescence times can provide information to discriminate between them. Recently Shchur et al. have shown a similar result for a model with continuous migration [32].

**Figure 2:**
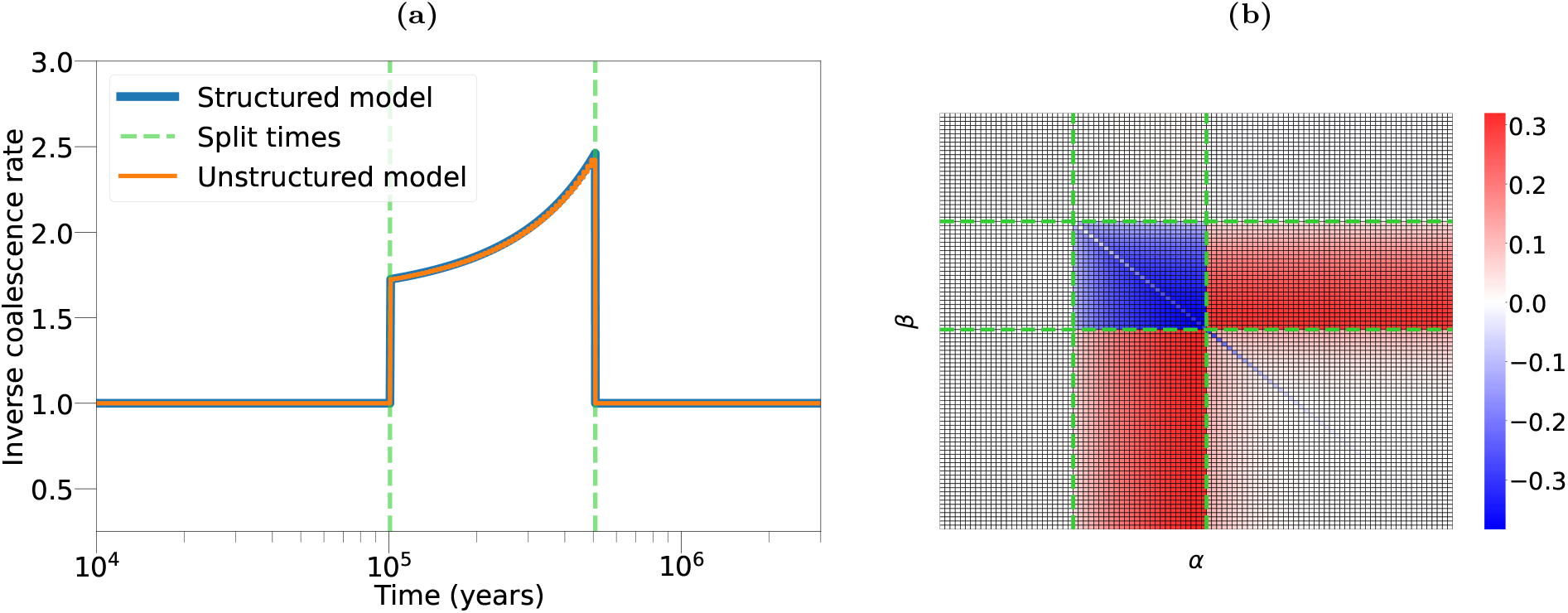
(**a**) Matching coalescence rate profiles for a structured and unstructured model. The blue line indicates the theoretical inverse coalescence rate for the structured model, where populations *A* and *B* are of constant size, the split/rejoin times are given by the vertical, dashed green lines, and the admixture fraction is 30%. The orange line indicates an unstructured model with size changes that generate a coalescence rate equal exactly to the structured model. (**b**) A visualisation of *ξ* = (*Q*^*U*^ − *Q*^*S*^)*/Q*^*S*^, the relative difference between the transition matrices for the structured and unstructured models in (a). *Q*^*U*^ and *Q*^*S*^ are row stochastic matrices, where *Q*(*α*|*β*) describes the probability of transitioning from discretised time window *β* to *α*, conditional on a recombination having occurred.

We introduce a new method for coalescence-based reconstruction of ancestral admixture, *cobraa*, which is an HMM [33, 34] to infer the parameters of our structured model. Like PSMC, *cobraa*’s hidden states are a set of discrete coalescence time windows, and the observations describe whether positions across the genome are homozygous or heterozygous. The emission matrix describes the probability that a mutation has arisen given a particular coalescence time, and the transition model describes the probability of an ancestral recombination event changing the local coalescence time as a function of the effective population sizes (*N*_*A*_(*t*)) in an unstructured model. The emission model and hidden states of *cobraa* are the same, but instead the transition matrix is a function of the parameters of the structured model (*N*_*A*_(*t*), *N*_*B*_, *γ, T*_2_ and *T*_1_)). A full description of *cobraa* is given in Supplementary Text.

### 3.2 *cobraa* ‘s power to infer parameters of a structured model

We show in Figure S2 that structure can be discriminated from lack of structure using a likelihood framework if the parameters are known. However, there are limits to the ability of any stochastic model, including coalescent HMMs, to identify parameters [35]. Using simulated data we explored how well the parameters of a structured model can be inferred with *cobraa* and whether it has power to distinguish ancestral structure from panmixia.

We first tested how recoverable the admixture fraction *γ* is, provided that the population sizes and split times are known. We simulated 10 replicates of a structured evolutionary history with constant size in population *A* and *B, µ/r* = 10, *T*_1_=250kya, *T*_2_=1Mya or 2Mya, and 3Gb of sequence for various values of *γ*. We ran *cobraa* until convergence (defined as the change in total log-likelihood being less than one, *ψ*_*L*_ *<* 1, between consecutive Expectation-Maximisation (EM) iterations) and plot the inferred admixture fraction in Figure 3a. The simulated value is generally well recovered, even if the admixture fraction is as small as 5%, though is increasingly underestimated as it gets larger. Similar to PSMC, we can take advantage of the rarity of recombination events to increase *cobraa*’s computational speed (and decrease memory usage) *b*-fold by binning the genome into windows of *b* base pairs. Parameter inference is not substantially affected with values of *b* up to 100, as seen in Figure S3, so from here we set *b* = 100.

**Figure 3:**
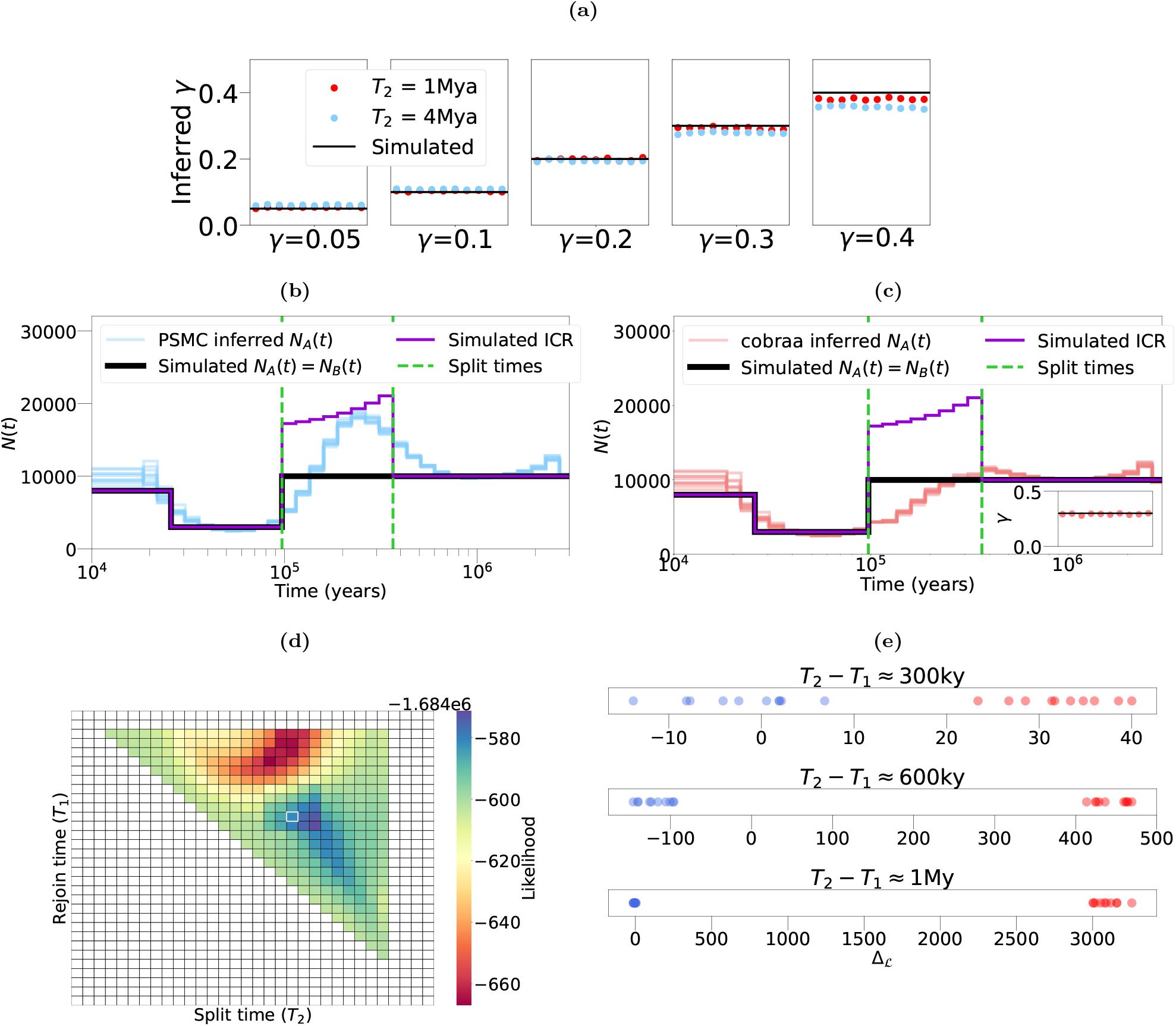
Ability of *cobraa* to infer parameters of a structured model. **(a)** Inferring the admixture fraction *γ*, when the population sizes and split/rejoin times are known. The simulated model has constant size in population *A* and *B, µ/r* = 10, *T*_1_=250kya, *T*_2_=1Mya or 4Mya, and 3Gb of sequence for various values of *γ*. Ten replicates are shown for each value of *γ*. **(b)** PSMC inference of *N*_*A*_(*t*), on a simulated structured model. The black line indicates simulated *N*_*A*_(*t*), which is the same as simulated *N*_*B*_(*t*). The green, dashed, vertical lines indicate the split and rejoin times (*T*_1_ =100ky and *T*_2_ =380ky, respectively) with *γ*=30%, *µ/r* = 1.25, and 3Gb of sequence. The purple line is the simulated inverse coalescence rate (ICR). **(c)** *cobraa* inference of *N*_*A*_(*t*) for the same structured model. **(d)** Using *cobraa* to search for *T*_1_ and *T*_2_ by iterating EM till convergence and recording the log-likelihood. The vertical axis represents the rejoin time *T*_1_, with values closer to the top indicating more recent times. The horizontal axis represents the split time *T*_2_, with values more right indicating more ancient times. Corresponding inference of *γ* for each pair is shown in Figure S4. **(e)** Difference in model fits between *cobraa* and PSMC, Δ_*ℒ*_ = ℒ_*S*_ − ℒ_*U*_, for a structured simulation (red points) and an unstructured simulation (blue points), both of which have the same coalescence rate profile. The top panel corresponds to the inference as shown in (b) and (c); inference of *N*_*A*_(*t*) and *γ* for the middle and bottom panel are not shown.

We next simulated a structured model with an admixture fraction *γ* of 30%, over a sequence length of 3Gb with *µ/r* = 1.25, as in humans, and a bottleneck between 25,000 and 100,000 years ago. Fifteen replicates were performed, on which we ran PSMC and *cobraa* until convergence. Inference from PSMC (light blue lines) on this model is shown in Figure 3b, where we see that the simulated population size changes are not well recovered in the structured period, with a false peak detected instead of a flat population size. The purple line indicates the simulated inverse coalescence rate (ICR), which PSMC interprets as the effective population size. Inference from *cobraa* on this model is shown in Figure 3c, where we see that the inferred changes in population *A* more closely reflect the simulated values. The inferred admixture fraction from each replicate is shown in the inset, and is relatively accurate. We also explored whether *cobraa* could recover changes in the size of the B population, *N*_*B*_(*t*), as well as *N*_*A*_(*t*), and found identifiability problems (see Supplementary Text), supporting our decision to maintain the size of population *B* constant over time.

The population size changes and admixture fraction can be inferred as part of the EM algorithm in *cobraa*, though the split and rejoin times are considered fixed in a single run. To search a range of split and rejoin times, we run *cobraa* over various (*T*_1_, *T*_2_) pairs, iterate each pair until convergence (*ψ*_*L*_ *<* 1) and record the log-likelihood (ℒ). These are shown in Figure 3d, with the simulated pair shown in the white cell. The maximum inferred log-likelihood is not exactly at the simulated pair, but it is close and in a relatively small neighbourhood of high scoring pairs, indicating that we have reasonable power to infer the split and rejoin times. In the region of high log-likelihood values, the inferred *γ* is also around the simulated value (Figure S4). By contrast, in the lower scoring pairs, the inferred *γ* is increasingly different from the simulated value, and in the minimal *T*_1_ / maximal *T*_2_ pairs the inferred *γ* is close to zero (Figure S4).

Next we explored the extent to which we are able to distinguish between a structured and a panmictic evolutionary history. We simulated from a series of paired panmictic and structured models with the same coalescence rate profiles, and calculated the difference between log-likelihoods obtained from fitting a stuctured and unstructured model for each data set Δ_ℒ_ = ℒ_*S*_ − ℒ_*U*_ . The results for various simulated split times are shown in Figure 3e. Consistently, we see that if the simulation was structured then inference from the structured model better explains the data than unstructured inference (as in PSMC), as seen by positive log-likelihood differences, Δ_ℒ_. The differences increase with the period of separation between the split and rejoin time, *T*_2_ − *T*_1_. Conversely, if the simulation was panmictic then structured inference is not able to explain the data any better than unstructured, as the log-likelihood differences are around 0 or negative.

### 3.3 Inference on human data

We use one high-coverage, whole-genome sequence from each of the 26 distinct human populations in the 1000 Genomes Project (1000GP) [24, 26]. To see how well an unstructured or structured model explains the data, we run PSMC (unstructured) and *cobraa* (structured) inference until convergence. To ensure that each sample uses the same discrete time interval boundaries, we fix the scaled mutation rate at *θ* = 0.0008 which is close to the mean across populations (Figure S5 and Table S1).

In Figure 4 we show *N*_*A*_(*t*) as inferred from the unstructured (PSMC) (4a) structured (*cobraa*) (4a) models on each population from 1000GP. For *cobraa* we take the composite maximum likelihood (CML) estimate of the split and rejoin times across populations (see Methods). The strongly positive Δ_ℒ_ values (Figure 4c) for each population indicate that a structured model explains the data much better than does a continuously panmictic model, as inferred by PSMC. The CML estimate of the split and admixture time (Figure 4b) suggest that two populations diverged ∼ 1.5Mya then subsequently admixed ∼290kya, around or shortly before the proposed origin of modern humans [36, 37]. The inferred admixture fraction indicates that present day humans derive 79% and 21% of their ancestry from ancestral populations *A* and *B*, respectively (Figure 4b).

**Figure 4:**
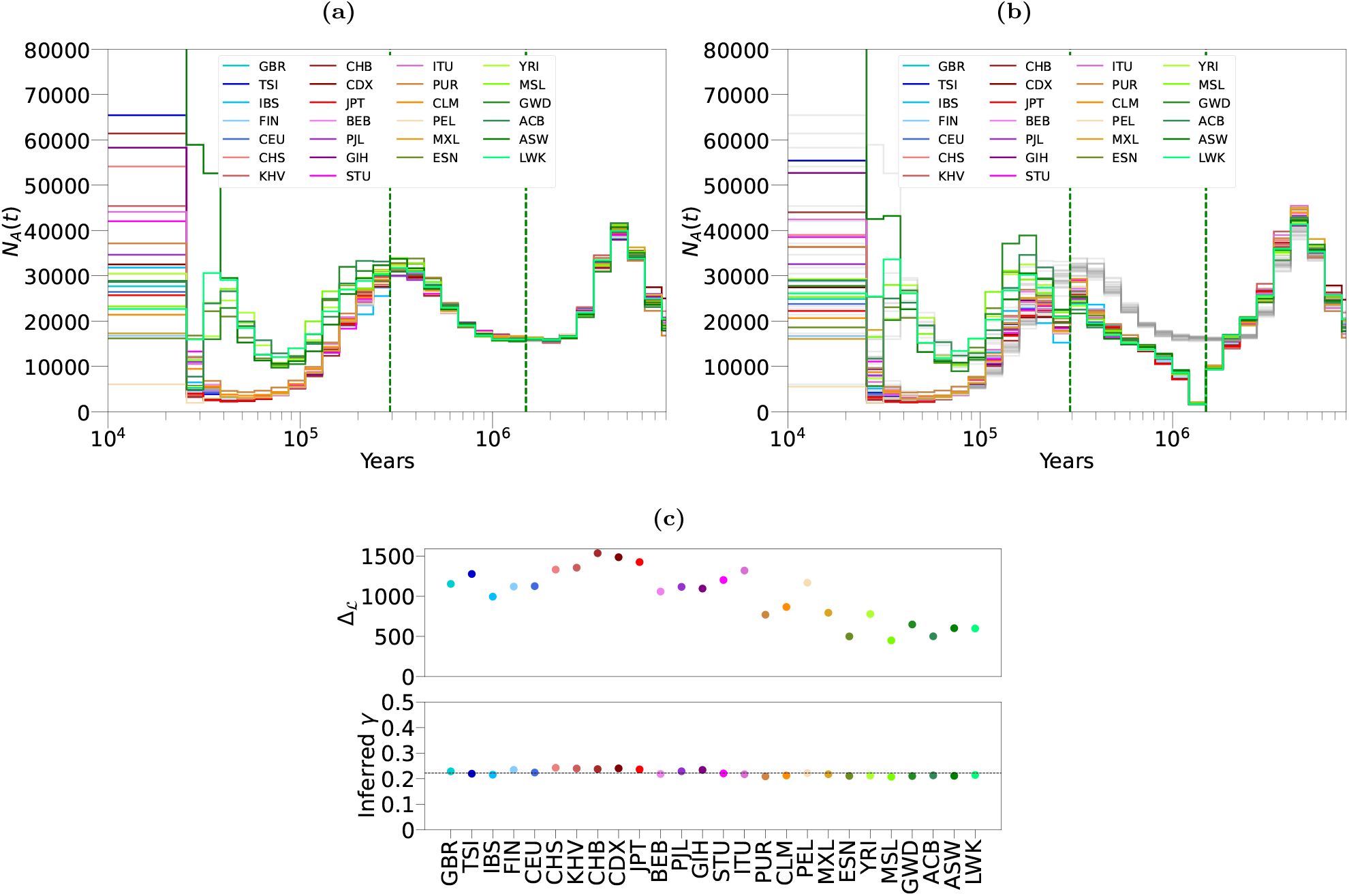
Inference from PSMC and *cobraa* on 26 populations from the 1000 Genomes project, using one individual per population. **(a)** PSMC’s estimate of *N*_*A*_(*t*). **(b)** *cobraa*’s estimate of *N*_*A*_(*t*), with the estimated split/admixture time shown in vertical, dashed, green lines. For direct comparison, the PSMC inference from (a) is also plotted in grey. **(c)** The upper panel shows the difference between the log-likelihood from *cobraa*’s inference and PSMC’s inference, Δ_ℒ_ for each population; the lower panel shows *cobraa*’s inferred admixture fraction *γ*. The inferred size of population *B* is shown in Figure S6.

Outside the structured period (more recently than the admixture time or more anciently than the split time), the inferred effective population changes in each population are very similar between the structured and unstructured models (Figure 4a and 4b). However, immediately more recently than the split time at ∼1.5My, in each 1000GP population the structured inference in *cobraa* infers a strong bottleneck in population *A*. The effective population sizes increases progressively until the admixture time. The population size estimates of population *B* are shown in Figure S6, and are generally larger than those inferred in *A*. The discrepancy between the population sizes inferred in *A* by *cobraa* and PSMC as well as other coalescence-based population size inference methods [38, 39, 40, 41, 42, 43]) are rather striking. However, our estimates are not necessarily inconsistent because most methods assume panmixia and as demonstrated theoretically by Mazet et al. ([20, 21]), for any panmictic model there necessarily exists a structured model than can generate the same coalescence rate history. To this end, we simulated 3Gb of sequence data from the *cobraa*-inferred structured model for various populations in Figure 4b and then ran unstructured (PSMC) inference on this. The inferred changes in population size are shown in Figure S7 and are extremely similar to those inferred by PSMC in real human data (Figure 4a), suggesting our structured model is compatible with previous coalescence-based estimations. We note our simulations suggest that the site-frequency spectrum could be used to distinguish between a structured or unstructured model (Figure S8; see Methods); thus a method combining coalescence and site frequency information to infer ancestral structure could be powerful [40, 32].

We also ran *cobraa* on the Human Genome Diversity Project (HGDP) [27, 28], and find very similar results (Figure S9) to those as inferred in 1000GP. Even in the San, who are estimated to harbor the most divergent ancestry across present day humans [44, 45, 37], the structured model is a substantially better fit to the data than a continuously panmictic model.

To explore the degree of support for the bottleneck in population *A* after divergence from *B*, we ran *cobraa* with various parameter constraints. We experimented with different levels of freedom in *N*_*B*_(*t*), and also tried optimisation after enforcing constant size in *A* (see Methods). We were not able to fit a model lacking a bottleneck in *A* that has comparable L, suggesting that the bottleneck is a necessary feature. We also investigated how the inference from *cobraa* changes in the presence of low quality regions of the genome, with widespread linked selection, or with more natural estimations of heterozygosity, and found that none of these have a significant effect on inference (see Methods).

### 3.4 Inferring admixed regions of the genome

We expanded the HMM of *cobraa* such that the hidden state represents not just the coalescence time, but also the path through the structured model taken by both lineages before they coalesce. We call this extended HMM *cobraa-path*. The hidden states are then a tuple (*t, c*) where *c* ∈ [*AA, BB, AB*] is the lineage path taken by the two lineages which coalesce at time *t*. If the coalescence time occurs more recently than the admixture event, then *c* = *AA*; if the coalescence occurred in the structured period, then *c* ∈ [*AA, BB*]; if the coalescence occurred more anciently than the population divergence, then *c* ∈ [*AA, BB, AB*]. An illustration of each possibility is given in Figure S10. The transition and emission probabilities of *cobraa-path* follow naturally from *cobraa* and are given in Supplementary Text. Running *cobraa-path* on simulations suggest we can infer *c* with reasonable accuracy using posterior decoding, though the Viterbi decoding appears to be limited S11a.

Using the inferred structured model from the previous section, we decoded each of the 26 samples from 1000GP and analyse the marginal posterior probability of each lineage path at each position, *P* (*c*_*i*_|*X*). We then condition these estimates on the coalescence time being larger than the admixture event, *P* (*c*_*i*_|*X, t > T*_1_), to reduce confounding. The correlation between *P* (*c*_*i*_ ∈ [*AA, AB, BB*]|*X, t > T*_1_) and the distance to closest CDS (dcCDS) on chromosome 1 for various populations is shown in Figure S12a. There is a weak but extremely significant Spearman correlation between dcCDS vs *P* (*c*_*i*_ = *AB*|*X*) and dcCDS vs *P* (*c*_*i*_ = *BB*|*X*) (∼0.081 and ∼0.078, respectively, with p-value*<*1e-100 for all populations; see Table S2 for full information) suggesting that genetic material from the minority population was selected against in modern humans post admixture. To confirm this, we examined the correlation between *P* (*c*_*i*_|*X, t > T*_1_) and a high resolution B-map [46], which integrates functional and genetic map information to estimate the strength of background selection across the genome. The Spearman correlation between *P* (*c*_*i*_ ∈ [*AB, BB*]|*X, t > T*_1_) and the B-map is larger than that with dcCDS and also extremely significant (∼0.21 and ∼0.31 for *AB* and *BB*, respectively, with p-value*<*1e-100 for all populations. See Table S2 for full information), as shown for various populations in Figure S12b, supporting the suggestion of negative selection on material from population *B*.

We next looked at regions of the genome that are enriched or depleted of inferred admixture, across all 1KGP populations. To do this, we define a test statistic *H*(*x*) which calculates for each position *x* the expected amount of admixture across all populations (see Methods). For 1kb regions that are in the top or bottom 1% of *H*(*x*) values, we looked to see if they overlap with any protein coding genes. We found 666 protein coding genes that overlap with regions in the top 1% of *H*(*x*) (∼3-fold enrichment above a neutral expectation; p-value *<* 1e-100), which we call admixture-abundant genes (AAGs), and 1270 protein coding genes that overlap with regions in the bottom 1% of *H*(*x*) (∼6-fold enrichment above a neutral expectation; p-value *<* 1e-100), which we call admixture-scarce genes (ASGs). We show an example of *H*(*x*) and inferred probability of admixture at KNG1 (an AAG) and FOXP2 (an ASG) in Figure S13.

To examine whether AAGs or ASGs are associated with any biological processes, we performed a gene ontology (GO) analysis [47, 48] using PANTHER [49] (see Methods). The 666 AAGs were reported to associate with multiple biological processes. Specifically, GO analysis reported an 11.6-fold enrichment for genes associated with neuron cell-cell adhesion, 8.5-fold enrichment for startle response, 6.2-fold enrichment for neuron recognition, 3.7-fold for neurotransmitter transport, 2.5-fold enrichment for chemical synaptic transmission, and 2.2 enrichment for circulatory system processing. Additionally, a twofold depletion in genes associated with gene expression was reported. Similarly, numerous associations between the 1270 ASGs and biological processes were reported. Notably, it reported an 8.3-fold enrichment for genes associated with pre-miRNA processing, 4-fold enrichment for cortical actin cytoskeleton organisation, and 3.6-fold enrichment for golgi to plasma membrane transport, among many others. Moreover, ASGs were found to have a 3.7-fold depletion in genes associated with adaptive immune response, 7.7-fold depletion in lymphocyte mediated immunity, 25-fold depletion in detection of chemical stimulus involved in sensory perception of smell, and *>*100-fold depletion in antimicrobial humoral response. More subprocess information is given in Tables S3 and S4, with full GO analysis (including parent categories) available online.

We next investigated whether population *A* or *B* is closer to Neanderthals and Denisovans. To test this, we examined the correlation between *P* (*c* = *BB*|*X, t > T*_1_) or *P* (*c* = *AA*|*X, t > T*_1_) and African to Neanderthal or Denisovan divergence (see Methods). In Figure 5 we show that African-archaic divergence decreases as *P* (*c* = *AA*|*t > T*_1_) increases, and that African-archaic divergence increases with *P* (*c* = *BB*|*t > T*_1_). The same is generally true for non-Africans (Figure S14), though the relationship is not monotonic, possibly due to recent gene-flow from Neanderthals and Denisovans [2, 4, 3]. The suggested association of recent archaic admixture with *B* ancestry is supported by a significant correlation between Sankararaman et al.’s [50] local probability of Neanderthal admixture and *P* (*c* = *AA*|*t > T*_1_) or *P* (*c* = *BB*|*t > T*_1_) (mean Spearman *ρ* -0.08 and 0.11, respectively, all p-values less than 1e-24. See Table S5 and Methods), indicating that regions from *B* are enriched in the same parts of the genome as regions of Neanderthal introgression in present-day non-Africans, possibly due to them both being selected against post admixture [51]. We note that if we condition on the coalescence time being older than *T*_1_ and younger than *T*_2_, the monotonicity in correlation with non-African to archaic divergence increases (Figure S14), which is consistent with the admixture event between population *A* and *B* being older than the time at which Neanderthals and Denisovans admixed with non-Africans. Finally, the same relationship holds when conditioning on the coalescence time being older than *T*_2_ (Figure S14); see Table S6 for correlations in each population. Overall, this is consistent with a scenario in which population *A* rather than *B* was the ancestral population to Neanderthals and Denisovans.

**Figure 5:**
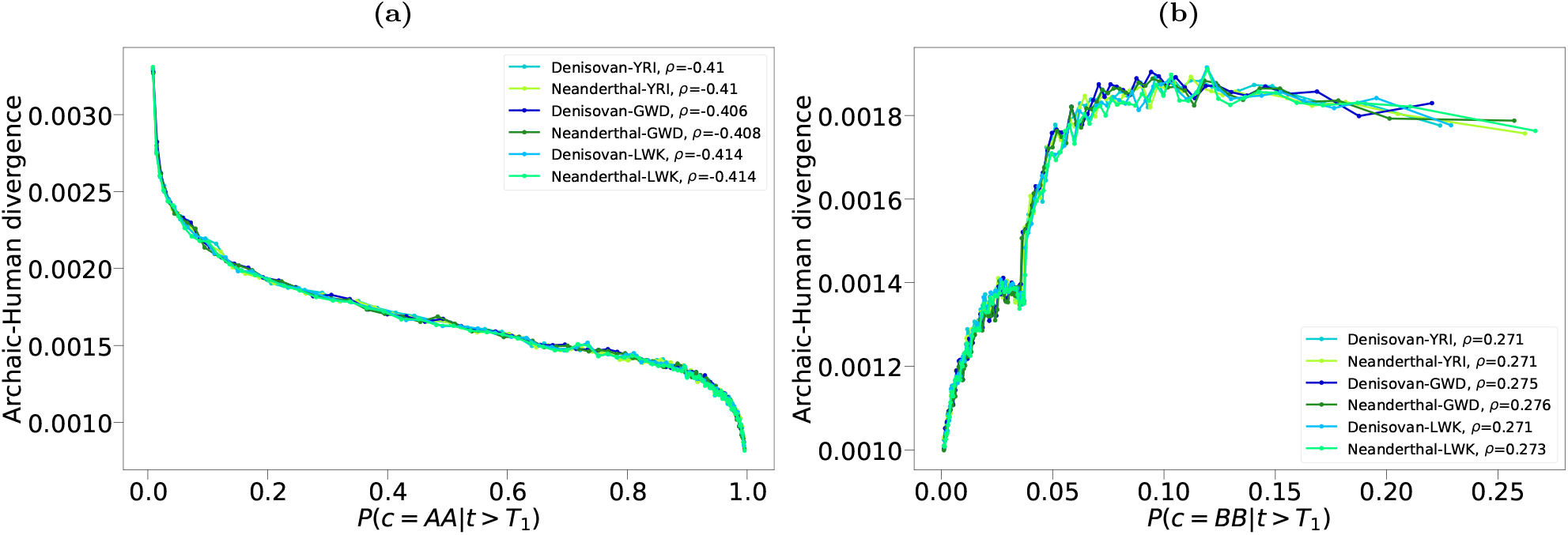
Relationship between the probability of a present day human region deriving from population *A* or *B*, and human-Neanderthal (greens) or human-Denisovan (blues) divergence, for three African samples. (a) As confidence in a region being assigned to *AA* increases, human to Neanderthal divergence and human to Denisovan divergence decreases . (b) As confidence in a region being assigned to *BB* increases, human to Neanderthal divergence and human to Denisovan divergence **i**ncreases. This suggests that Neanderthals and Denisovans derive from population *A*. We used the Altai Neanderthal genome and calculated the divergence in windows of 5kb. These relationships are shown for all 1000GP populations in Figure S14, along with different conditioning on the coalescence time. All Spearman correlations are reported in Table S6.

## 4 Discussion

We introduce here a new method to infer a structured ancestry from a diploid genome sequence, which we have used to infer a deep split in human ancestry 1.5 million years ago, rejoining 300 thousand years ago, around the time of the first evidence for anatomically modern humans. The evidence that Neanderthal and Denisovan genomes diverged more recently from ancestry on the major (*A*) lineage, rather than the minor (*B*) lineage, supports the conclusion that the structure we infer is neither artefactual nor arbitrary, since the inference procedure is independent of these archaic genomes. A simplified diagram of human evolutionary history, with *cobraa*’s contribution highlighted in red, is shown in Figure 6. We provide evidence of general selection against introgressed material from the minor (*B*) lineage, while also seeing enrichment of introgression specifically in a set of categories associated with neuronal development and processing. The admixture percentage of ∼20% is much higher than the fraction of Neanderthal or Denisovan admixture into present-day non-African populations, but was not discernible using standard f-statistics because it is shared by all present day humans. We note that similarly high admixture fractions with enrichment of admixture in gene categories important in speciation have been seen at the base of speciation/radiation events in other taxa, for example in Lake Malawi cichlids [52].

**Figure 6:**
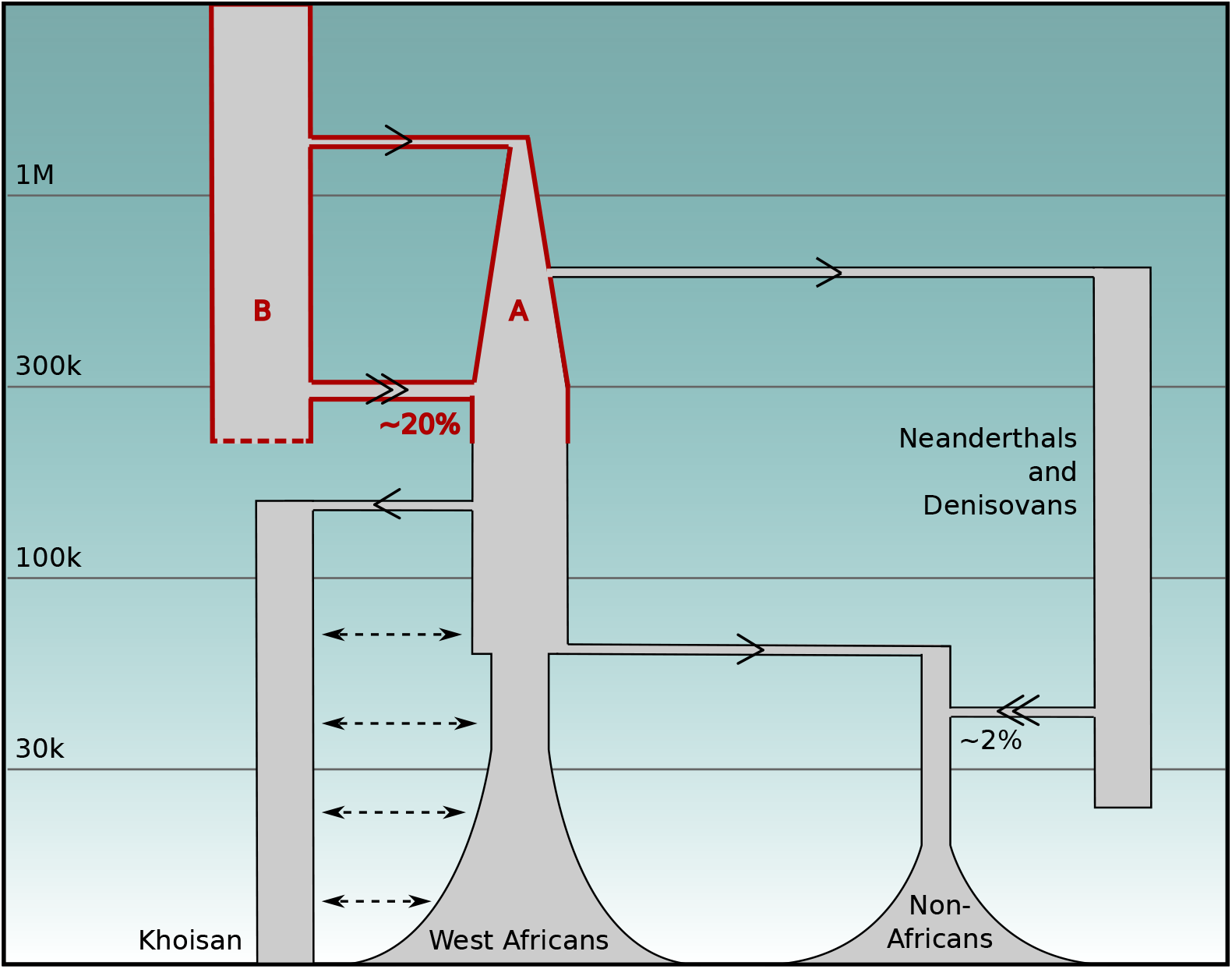
A simplified model of human demographic history showing deep population structure 1.5My 300ky ago shared by all present-day humans, as inferred by *cobraa* (red). Arrows indicate the direction of gene flow, with admixture events (double arrows) labeled by their percentage genetic contribution to the recipient population. Of the two ancestral branches *A* and *B, A* represents 80% of subsequent ancestry and features a sharp bottleneck immediately after its founding. Dashed arrows between Khoisan and other African populations reflect the fact that this divergence, the deepest among present-day human populations, has involved ongoing or intermittent gene flow [32, 37, 39]. The y-axis represents time in years before present.

Recently, Ragsdale et al. also proposed deep ancestral structure in the ancestors of modern humans [16]. Their best fitting model suggested an ancestral population diverged ∼1.7Mya into two populations, “stem 1” and “stem 2”, with continuous gene-flow between them until ∼500kya at which point stem 1 split into populations 1S and 1E. Stem 1S is estimated as being a primary ancestral population to the Khoisan, and stem 1E to all other modern human subpopulations. Stem 2 is estimated to have contributed 70% of ancestry into the ancestors of the Khoisan ∼120kya, and 50% of ancestry into stem 1E ∼ 100kya. The estimated split time of ∼1.7Mya between stem 1 and 2 is remarkably similar to the split time as estimated by *cobraa*, though our estimates of admixture time are not well aligned. Notably, we estimate a single admixture event occurred ∼300kya, which is substantially more ancient than the earliest admixture reported by Ragsdale et al. (stem 2 to the ancestors of Khoisan ∼120kya). This could due to the space of models considered by *cobraa*, which for example requires population *A* and *B* to be in isolation after they split from an ancestral population, contrasting with Ragsdale et al.’s inferred continuous gene-flow between stem 1 and 2. Conversely, the model in Ragsdale et al. assumes a constant population size in stem 1 and 2, whereas *cobraa* allows these to be variable over time and we found that a early bottleneck in population *A* was an important feature when fitting the model to human data.

Numerous authors have reported evidence for there being more recent contributions of unknown archaic ancestry to modern humans, especially in West Africans [9, 10, 11, 12, 13, 14, 15, 16, 53, 54]. Parametric estimates vary, though all models of structure in West Africans infer that admixture occurred more recently than ∼150 thousand years ago [13, 14], with many even inferring it more recently than 50 thousand years [9, 10, 11, 12, 15]. Moreover, the inferred population divergence time is always estimated as being more recent than 1 million years ago. Although this appears to be a different event to the one that we describe, not shared by all present day humans, these inferences suggest a plausible reason why the *cobraa*-inferred maximum likelihood estimates of the split and rejoin time in Africans are more recent than the composite maximum likelihood estimate (Figure S15).

Technically, we have demonstrated that the conditional distribution of adjacent coalescence times has information about ancestral structure, partly overcoming the pairwise coalescence rate identifiability problem [20, 21]. Recent theoretical work demonstrates that the coalescence rate from three lineages (i.e. joint density of the first and second coalescence events) can also distinguish population structure from population size changes [55]. Thus, a method specifically estimating the second as well as the first coalescence rate could help extend our approach.

There are several caveats to our approach. We have shown that a pulse model of structure better fits the data than does a continuously panmictic model, but even if we limit ourselves to older events shared by all modern humans, our evolutionary history is very likely more complex than this (Figure 1a, [16]). We can not rule out more complicated scenarios such as there being more than two divergent populations admixing together, numerous split and rejoin events, or other models incorporating more continuous gene flow.

Additionally, there are assumptions in the PSMC/*cobraa* framework which are known not to hold. Most notably, the assumptions of an absence of selection and constant mutation and recombination rates across the genome are false [56, 57, 58, 59, 60, 61, 62, 63, 64, 65, 66, 67, 68, 69]. However previous studies that incorporated realistic variation in mutation and recombination rates into simulations have shown that they only have negligible consequences for SMC-based inference of population size changes over time [19], and [70] shows that genomic variation in linked selection does not have a noticeable effect on the identification of structure.

The model of ancestral structure we propose raises intriguing questions about the relationship of lineages *A* and *B* to previously identified hominins. Archaeological evidence suggests numerous forms of archaic hominins, and it is not settled which if any contributed directly to the ancestry of modern humans, or split from the main lineage and subsequently died out without contributing any genetic material [17]. Various *Homo erectus* and *Homo heidelbergensis* populations that are potential candidates for lineages *A* and *B* existed both in Africa and elsewhere in the period prior to this. It is tempting to ascribe the sharp bottleneck that we infer in lineage *A* after separation from lineage *B* to a founder event potentially involved with migration and physical separation. Furthermore the ancestors of Neanderthals and Denisovans were in Eurasia before modern humans expanded there, and we can ask whether the gene flow from the ancestors of modern humans into Neanderthals [7, 71, 72, 6, 8] came from *A* or *B*, and also how the proposed archaic gene-flow event into Denisovans [2, 4] was related to these populations. Further clarifying the genetic contributions to modern humans and connecting them to the fossil record is an ongoing challenge.

## 5 Acknowledgements

We are grateful to members of the Durbin and Scally groups, especially Regev Schweiger and Casper Siu, for helpful discussion. We are also grateful to David Reich for his comments on an early version of the manuscript. T.C. was funded by a Wellcome Postgraduate Studentship 108864/B/15/Z, and R.D. by Wellcome Investigator Award 207492/Z/17/Z. For the purpose of open access, the author has applied a CC BY public copyright licence to any Author Accepted Manuscript version arising from this submission.

## 6 Data and code availability

*cobraa* is freely available to use and download at github.com/trevorcousins/cobraa. Processed 1000GP sequence data, summaries of posterior decoding, the list of AAGs and ASGs, and full GO analysis are available to download and analyse at doi:10.5281/zenodo.10829577. Key scripts for reproducibility are also given at github.com/trevorcousins/cobraa/reproducibility.

## 7 Methods

### 7.1 Model summaries

*cobraa* is a hidden Markov model (HMM) which builds on the PSMC framework. The hidden states are the discretised coalescence times across the genome, and the observations are the series of homozygotes or heterozygotes in a diploid genome sequence. The model parameters are *N*_*A*_(*t*), *N*_*B*_(*t*), *γ, T*_1_, and *T*_2_, which are, respectively, the population sizes in the sampled population, the population sizes in the ghost population, the admixture fraction, the rejoin time and the split time.

The emissions describe the probability of a mutation arising given a particular coalescence time. The transition matrix *Q*(*N*_*A*_(*t*), *N*_*B*_(*t*), *γ, T*_1_, *T*_2_) is governed by the SMC framework [30, 31], as a function of a structured model’s parameters. Calculating the transition probabilities required considering the 10 distinct possibilities for changes in coalescence time due to recombination or migration, as shown in Figure 1b. The population sizes and the admixture fraction can be optimised as part of the EM algorithm, though the split/rejoin times are fixed and are searched through independent model runs. We also note that when the admixture fraction *γ* is equal to zero then this corresponds to an unstructured model exactly as in PSMC, so PSMC is nested in *cobraa*.

When running *cobraa* on the 1000GP data, we enforce every sample has the same discrete time interval boundaries by fixing *θ* across all populations. This is so that we can use a composite maximum likelihood (CML) search over all possible pairings of the split and rejoin time. More explicitly, for a population *j* and split and rejoin time of *t*_1_ and *t*_2_, we get the log-likelihood ℒ (*j, t*_1_, *t*_2_) by running *cobraa* till convergence. We then select the CML time pairing with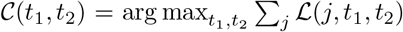 . We enforced that *N*_*B*_(*t*) = *k* for all *t*, where *k* was optimised as part of the EM algorithm, due to identifiability problems between *N*_*A*_(*t*) an *N*_*B*_(*t*) (Figures S26 and S27.)

We expanded *cobraa* into a new HMM, *cobraa-path*, whose hidden states are the ancestral lineage path, *c*, and the discretised coalescence times. If the sampled lineages coalescence more recently than the admixture event, then they can only coalesce in population *A* so *c* = *AA*. If they coalesce more anciently than the admxiture time but more recently than the split time, then either both lineages stayed in population *A* or they both migrated to *B* so *c* ∈ [*AA, BB*]. If they coalesce more anciently than the split time, then they either both stayed in *A*, both migrated to *B*, or one stayed and one migrated, so *c* ∈ [*AA, AB, BB*] (see Figure S10). The emission probabilities follow from *cobraa*, though are repeated across different values of *c*. For example, given a coalescence of *t* that is more ancient than population divergence, the probability of observing a mutation is the same for each *c* ∈ *AA, BB, AB*. The transition probabilities also follow naturally from *cobraa*, though do not require explicit calculations of the lineage path at the previous locus. We also note that if *γ* = 0 then *cobraa-path* reduces to standard PSMC.

The main advantage of using *cobraa-path* is that we can decode the HMM to infer the regions of admixture, i.e. where the ancestral lineage path went partially or wholly through the minority population. Using the forward/backward algorithm we can thus rapidly obtain the joint posterior probability of the ancestral lineage path and coalescence time at each position *P* (*c*_*i*_, *t*_*i*_|*X*, Θ), where *X* is the observed data and Θ is the set evolutionary parameters given to *cobraa-path*. The marginal posterior probabilities of each ancestral lineage path *c* are easily obtained by summing over the coalescence times *P* (*c*_*i*_|*X*, Θ) =∑_*τ*_ *P* (*c*_*i*_, *t*_*i*_ = *τ* |*X*, Θ).

To calculate the expected amount of admixture at each position *x*, we define:

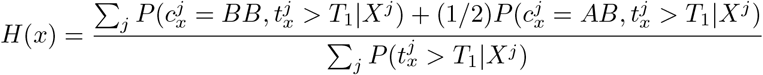

where *j* represents each population, *X*^*j*^ is all observed data for population *j*, and 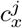 and 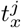 is the ancestral *x* lineage path and coalescence time, respectively, for population *j* at position *x*.

Full mathematical details of *cobraa* and *cobraa-path* are given in Supplementary Text.

### 7.2 Associations between *cobraa-path* and functional information

The positions of coding sequence (CDS) and their annotations were obtained from HAVANA, as downloaded from GENCODE ftp.ebi.ac.uk/pub/databases/gencode/Gencode_human/release_45/gencode.v45.chr_patch_hapl_scaff.basic.annotation.gff3.gz. To obtain the protein coding genes overlapping the top or bottom *H*(*x*) values, we looked to see if *x* occurred between the start and stop position of any CDS. To save disc space *H*(*x*) was inferred every 1kb, and positions not passing the GRCh38 mappability mask were excluded from analysis. This left 2,158,664 positions at which *H*(*x*) could be confidently calculated, and thus 21,587 positions in the top or bottom 1% of *H*(*x*) values. A binomial test with probability of success of 0.01 was used to calculate significance values for fold enrichment for AAGs and ASGs.

For the GO analysis we entered our AAGs or ASGs into `geneontology.org` and only considered associations where p-value (Fisher’s exact test) and FDR (Benjamini-Hochberg) *<* 0.05. In Tables S3 and S4, superclasses of each process are not shown, though are available for download at doi:10.5281/zenodo.10829577.

We used the B-map as inferred in [46], and lifted over from GRCh37 to GRCh38 [73]. Positions that did not pass the GRCh38 mappability mask were excluded from the analysis.

### 7.3 Processing 1000GP and HGDP Data

We took high-coverage whole-genome-sequence cram files for one individual in each of the 26 populations from the 1000 Genomes project. These are aligned to GRCh38. The cram files were converted to bam and indexed with samtools [74, 75]. The genotype likelihoods were calculated with bcftools mpileup [76] by skipping alignments with mapping quality less than 20, skipping bases with base alignment quality less than 20, and setting the coefficient for downgrading mapping quality to 50. SNPs were called using bcftools and all indels were excluded. Variants were then designated as uncallable if the minimum mapping quality was less than 20, the minimum consensus quality was less than 20, or the coverage was less than half or more than double the mean coverage. Finally, we designated all regions in the strict mappability mask for GRCh38 (ftp.1000genomes.ebi.ac.uk/vol1/ftp/data_collections/1000_genomes_project/working/20160622_genome_mask_GRCh38) as uncallable. Uncallable positions are labelled as missing data in the HMM. See Table S1 for information regarding number of heterozygous, homozygous, and uncalled positions for each individual. The triplet codes used by the 1000 Genomes Project for each population is given in Table S7.

HGDP data was downloaded from ftp.1000genomes.ebi.ac.uk/vol1/ftp/data_collections/HGDP, and was processed in exactly the same way.

### 7.4 Model convergence and fitting

For structured and unstructured model fitting, we iterated the EM algorithm until the change in log-likelihood was less than 1. The value of 1 was somewhat arbitrary, but is a convenient stopping criteria that allowed us to be consistent across different models. To check that we were not overfitting, we took the inferred parameters on seen data (training) and calculated the log-likelihood of unseen data (testing), by taking a new individual from each population. As shown in Table S8, the differences in log-likelihood for the structured and unstructured model in the test data are also strongly positive, suggesting that a better structured model is not due to overfitting.

### 7.5 Effect of different parameter constraints

To investigate whether the inferred bottleneck (Figure 4b and S9) is attributable to population *A*, we reran *cobraa* after relaxing the constraint that *N*_*B*_ must be constant (Figure S16). Figure S16a indicates that *cobraa* still infers a bottleneck in *N*_*A*_(*t*), with estimates of *N*_*B*_(*t*) generally being large immediately post divergence and decreasing until the time of admixture, though with greater variance across populations compared to *N*_*A*_(*t*). The inferred admixture fraction is extremely similar, and the fit of this model is slightly better than when we enforced constant *N*_*B*_ (Figure S16b), which is unsurprising due to there being more parameters. We also examined how well the model fits the data with *N*_*B*_(*t*) not being large post divergence, and find that this model is still well supported (Figures S16b and S16c).

In order to explore the degree of support for the bottleneck inferred by *cobraa* soon after the split at ∼1.5Mya, we constrained the parameters to search for a constant *N*_*A*_(*t*) during the structured period, but conclude that is not well supported by the data, as seen in Figure S17. Notably, the ℒ difference Δ_ℒ_ is often negative, and the split/admixture times and admixture fraction are not consistent, indicating that even a panmictic model fits the data better than this constrained structured model, and that the variation in *N*_*A*_(*t*) is necessary for the better *cobraa* fit.

To search for the optimal split and rejoin time, we ran *cobraa* over various values of *T*_1_ and *T*_2_, independently for each population representative. In each case we iterated till convergence (*ψ*_*L*_ *<* 1) and recorded the ℒ. The differences in ℒ between each pair and the panmictic inference from PSMC 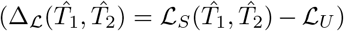 are shown in Figure S15, where red indicates positive 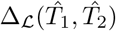 and white zero. The maxmimum varies a bit between samples, but is always within one cell of the CML except for West African populations, where a more recent structured event is preferred. We return to this observation in the discussion.

### 7.6 Effect of low quality data in the HMM

Low quality regions of the genome are labelled as missing data in the observations of the HMM. To check that this was not artificially inflating likelihood differences between the panmictic or structured model, or biasing inference, we reran inference after removing the centromeres and telomeres, as the GRCh38 mappability mask labels them as uncallable. This involved splitting each chromosome into two parts, the first of which begins after the end of the telomere and ends before the start of the centromere. The second part starts after the end of the centromere and ends before the start of the telomere. Doing this for each chromosome resulted on average in removing ∼100Mb of missing data in each population. These new chromosome parts were then given to PSMC or *cobraa* to run inference with composite-likelihood optimisation.

The resulting inference was practically indistinguishable from the full data (Figure S18). The differences in Δ_ℒ_ and the inferred admixture fraction between each data set were extremely similar, as seen in Figure S18, suggesting that the evidence for the structured model is not due to large regions of missing data in the HMM.

### 7.7 Effect of fixing *θ* across populations

To enforce parameter estimates across populations used the same discrete time interval boundaries, we fixed the scaled mutation rate *θ* at 0.0008, despite there being as much as a ∼45% difference between the highest and lowest population (PEL=0.00069, ESN=0.0010; Figure S5 and Table S1). We checked that our inference was not misled by this constraint by rerunning the analysis with *θ* inferred in the natural sense (Figure S19). The inferred split and rejoin times are noisier (Figure S19b) due to the time interval boundaries not aligning, but the inferred population sizes, admixture fraction, and ℒ differences are similar. Thus we conclude the evidence for the structured model is not due to fixing *θ*.

### 7.8 Effect of widespread linked selection

It has been demonstrated that widespread linked selection is pervasive in humans [77, 46], though PSMC and *cobraa* assume the genome evolves neutrally. Wrongly assuming neutrality has been shown to affect demographic inference [78, 79, 80], though solutions have been proposed [81, 82, 83, 70]. To check that widespread linked selection is not falsely interpreted as structure, we ran *cobraa* on an unstructured evolutionary history with linked selection. We used the SLiM [84] simulated data in [70], where the coalescent rate profile was constructed to mimic that as inferred in West Africans, and the distribution of fitness effects was chosen according to parameters that were inferred in humans [85]. *N* was scaled down to avoid excessive memory usage, and *µ* and *r* were chosen such that diversity as a function of distance to exon imitated that as observed in humans. For inference, we ran PSMC and *cobraa* until the change in log-likelihood in subsequent iterations of the EM algorithm was less than 0.1. In Figure S20 we show that *cobraa* is not able to explain the data any better than PSMC, indicating that widespread selection is not interpreted as structure.

### 7.9 Site-frequency-spectrum

In addition to coalescence-based approaches, it is possible to infer population history from the site frequency spectrum (SFS) [86, 87, 88, 89, 90, 91]. However, even for inference of a panmictic history (as assumed by most methods) this is an ill-posed problem [92, 93, 94, 95, 96], in that many different size histories can generate the same SFS. Despite this, we note that our simulations suggest the SFS for a structured model is distinct from the SFS for an unstructured model with the same coalescence rate profile, as shown in Figure S8. This contrasts with the identifiability problem from the pairwise coalescence rate profile [20, 21], and in principle suggests that a method that uses the SFS to jointly estimate population size changes and ghost admixture could have power to detect the structured event we propose [91].

### 7.10 Neanderthal and Denisovan Divergence

We downloaded the Altai Neanderthal genome sequence [3] and the Denisovan genome sequence [4]. We polarised the ancestral allele by ensuring that the human reference allele matched the chimpanzee and gorilla allele, and excluding all sites that did not satisfy this. Variant positions were aligned to GRCh37, so we used LiftOver to convert these to GRChg38 [73]. To adjust for phasing uncertainty in both humans and archaics, we randomly sampled the genotypes (kept homozygous derived with probability 1, kept heterozygous with probability 0.5, and kept homozygous ancestral with probability 1). We calculate divergence in windows of 5kb or 10kb, noting that the signal for each was not overly different, and took *cobraa-path*’s mean probability of admixture from *A* or *B* in windows of the same size.

We downloaded Sankararaman et al.’s inferred probability of Neanderthal admixture [50] from https://reich.hms.harvard.edu/datasets/landscape-neandertal-ancestry-present-day-humans, and lifted over to GRCh38. We note that Sankararaman et al. give inferred probabilities per 1000GP population, as opposed to individual-specific probabilities which is what we have from *cobraa-path*.

## 8 Supplementary Figures and Tables

### 8.1 Figures

**Supplementary Figure 1:**
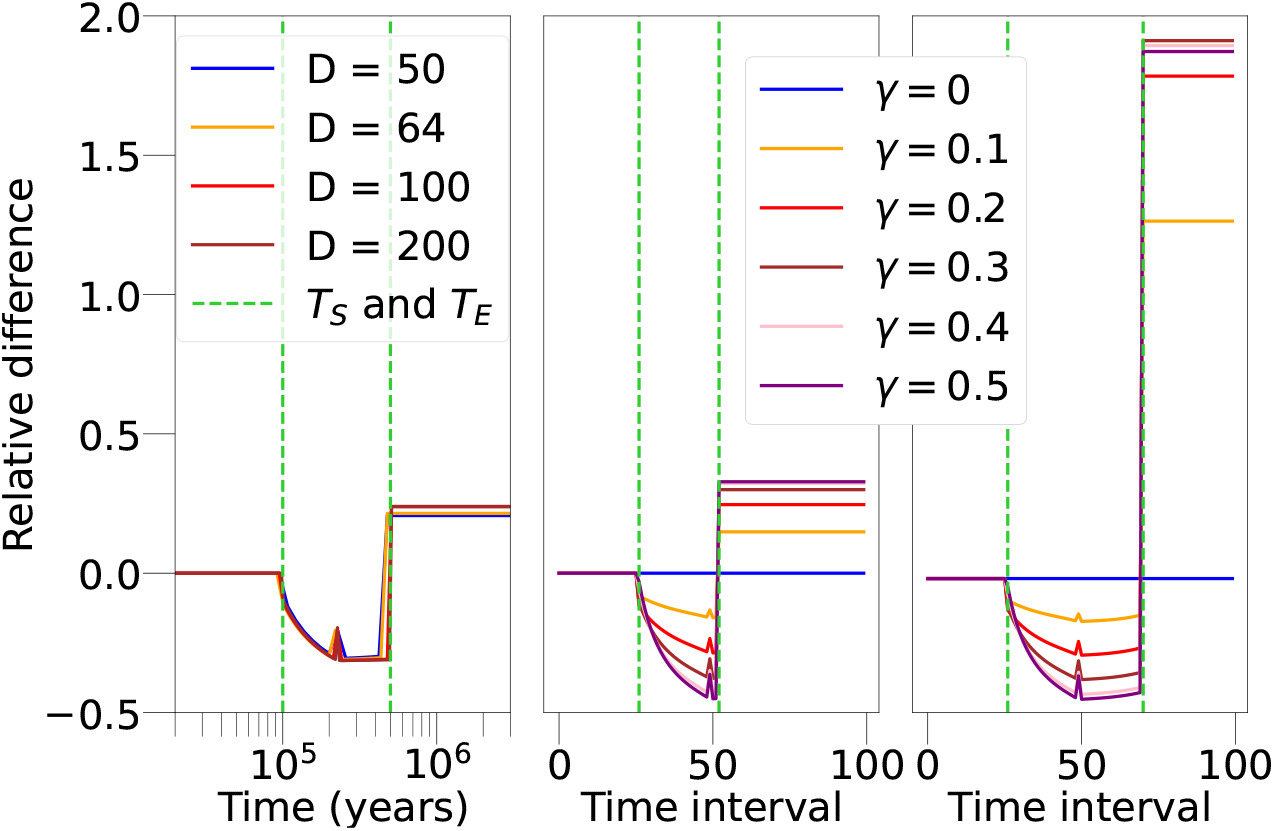
Relative difference in transition matrices for different model hyperparameters and parameters. For the matrix *ξ* in Figure 2b, plot a row which corresponds to a given coalescence time in the structured period. The x-axis corresponds to the time (left panel, time in years; middle and right panels, state indices of the HMM), with the vertical, green, dashed lines indicating the split and rejoin time of the structured model. Left panel has the same *pulse* parameters but an increasing number of discrete time intervals, *D*. The relative difference does not increase or decreases as function of *D*, indicating that this result is not a consequence of time discretisation. Middle and right panels show how the relative difference increases as does the admixture fraction, *γ*. The right panel has a longer period of separation between populations *A* and *B* than does the left or middle panel, which also increases the relative difference.

**Supplementary Figure 2:**
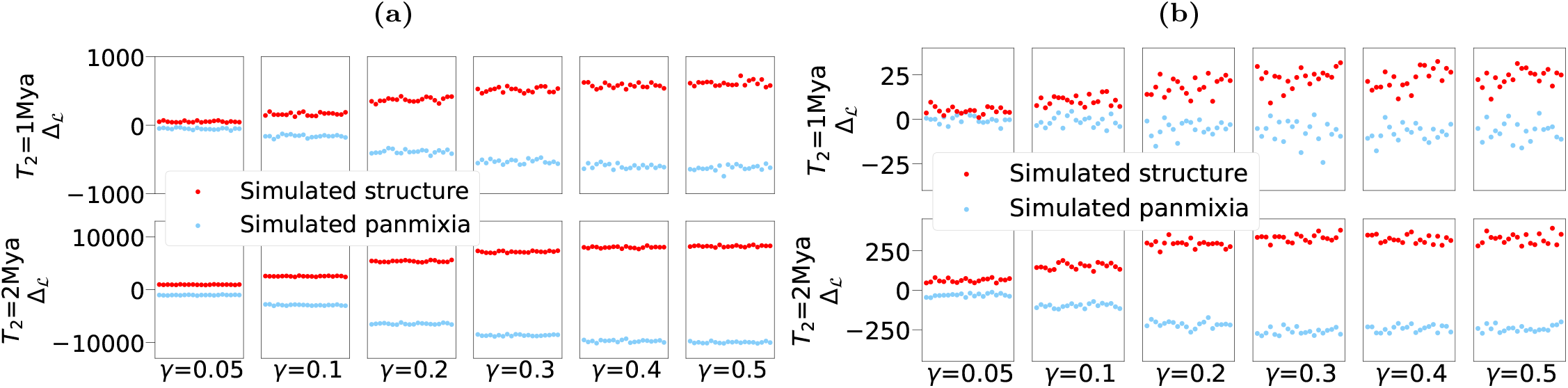
Differences in log-likelihood (ℒ) between a simulated structured or unstructured model, each with the same coalescence rate. We simulated 20 replicates of each model, with a sequence length of 100Mb, population size *N* = 2e+4, mutation rate per base pair per generation *µ* = 1.25e-8, and recombination rate per neighbouring base pairs per generation *r* = 1e-8. When simulating from the structured model we let the admixture fraction vary between 5% and 50%, set *T*_1_ as 200kya and *T*_2_ as 1Mya or 2Mya; when simulating from the unstructured model we create a series of size changes such that the coalescence rate is exactly the same as the corresponding structured model. For each simulation, we calculate the ℒ of the data under a structured and unstructured model. Thus there are 4 possibilities: structured simulation with structured inference, structured simulation with unstructured inference, unstructured simulation with structured inference, and unstructured simulation with unstructured inference. We plot the LL difference between the structured and unstructured inference, Δ_ℒ_ = *S*ℒ_*S*_ − ℒ_*U*_, and colour the points red when the simulation is structured and blue when the simulated is unstructured. **(a)** Data are the sequence of coalescence times across the genome (i.e. the probability of the Markov chain). We see clearly that the loglikelihood is able to distinguish the two histories, because Δ_ℒ_ is positive when the simulated evolutionary history is structured and negative when the simulated evolutionary history is panmictic. This is true even for a small admixture fraction, and the ℒ differences increase as does the admixture fraction or the separation time **(b)** Data are the sequence of heterozygotes or homozygotes across the genome (i.e. the probability of the HMM). If both the admixture fraction and separation time are sufficiently large then the two evolutionary histories are distinguishable using their ℒ, though it seems small admixture fractions are not identifiable with a 100Mb sequence. The magnitude of difference in ℒs between (a) and (b) reflect the uncertainty in inferring the coalescence time. The increase in Δ_ℒ_ as the admixture fraction or separation time increase is consistent with the relative error plots as seen in Figure S1.

**Supplementary Figure 3:**
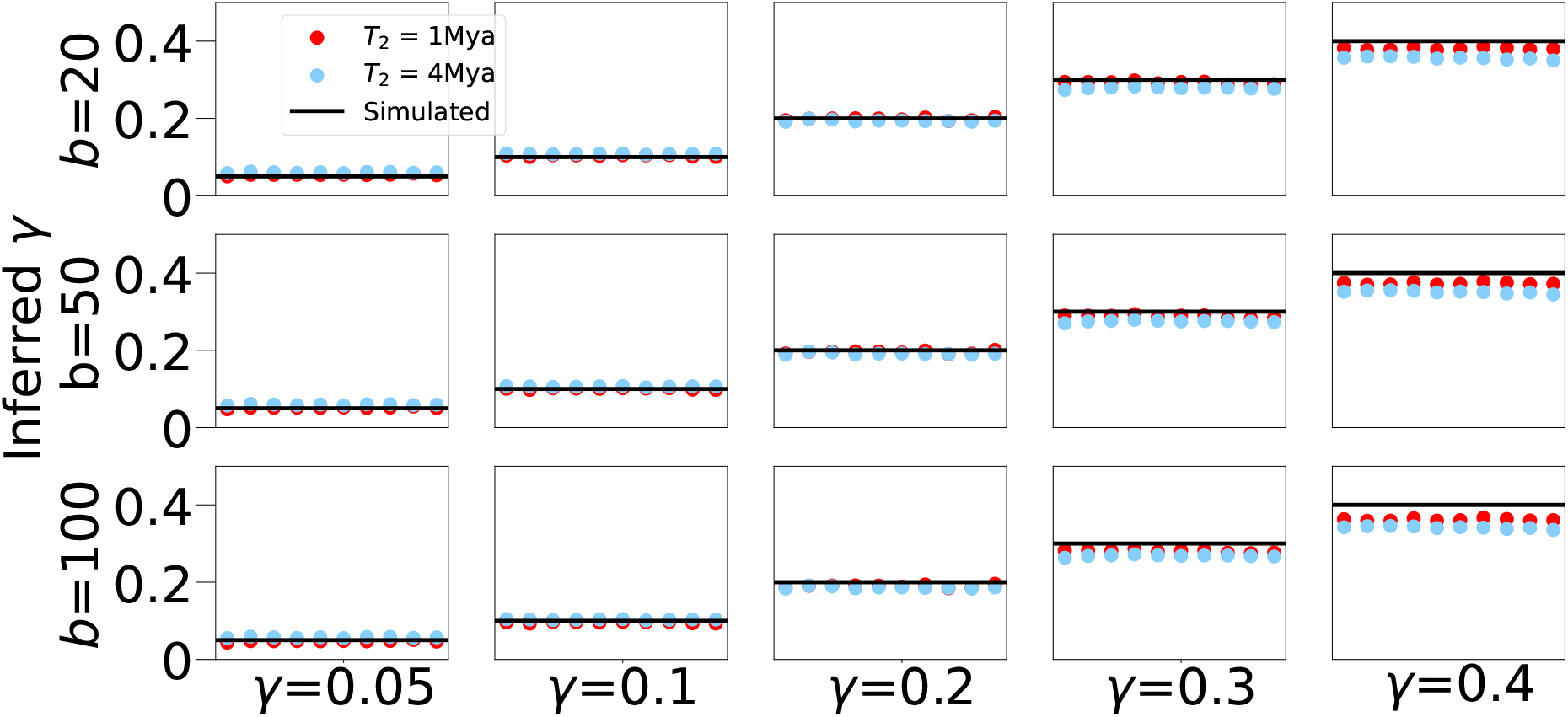
Ability of *cobraa* to infer the admixture fraction *γ*, when the population sizes and split/rejoin times are known, for various bin sizes *b*.

**Supplementary Figure 4:**
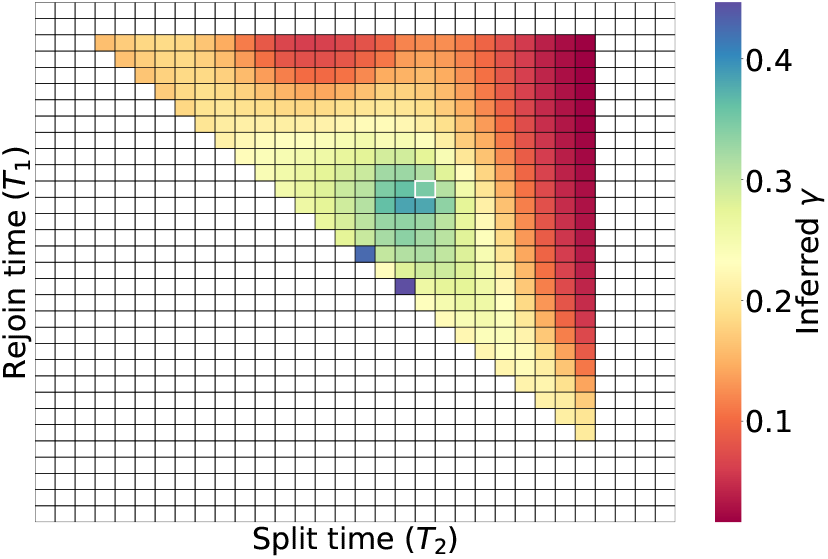
The *cobraa* inferred admixture fraction *γ*, for each time pairing (*T*_1_, *T*_2_). The corresponding log-likelihood for each pairing is shown in Figure 3d.

**Supplementary Figure 5:**
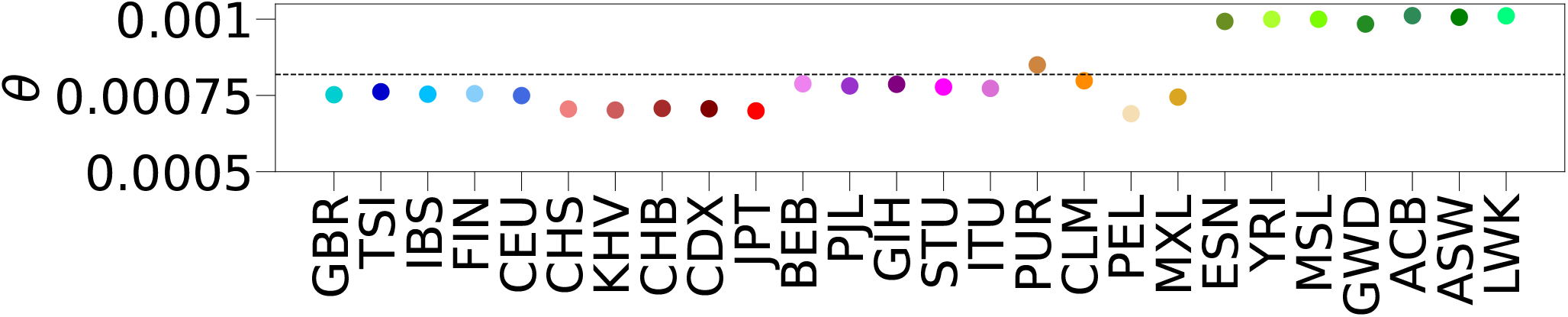
The scaled mutation rate *θ* = 4*Nµ* per population in the 1000 Genomes Project, using Waterson’s estimator. The dashed, horizontal line indicates the mean. See Table S1 for full information.

**Supplementary Figure 6:**
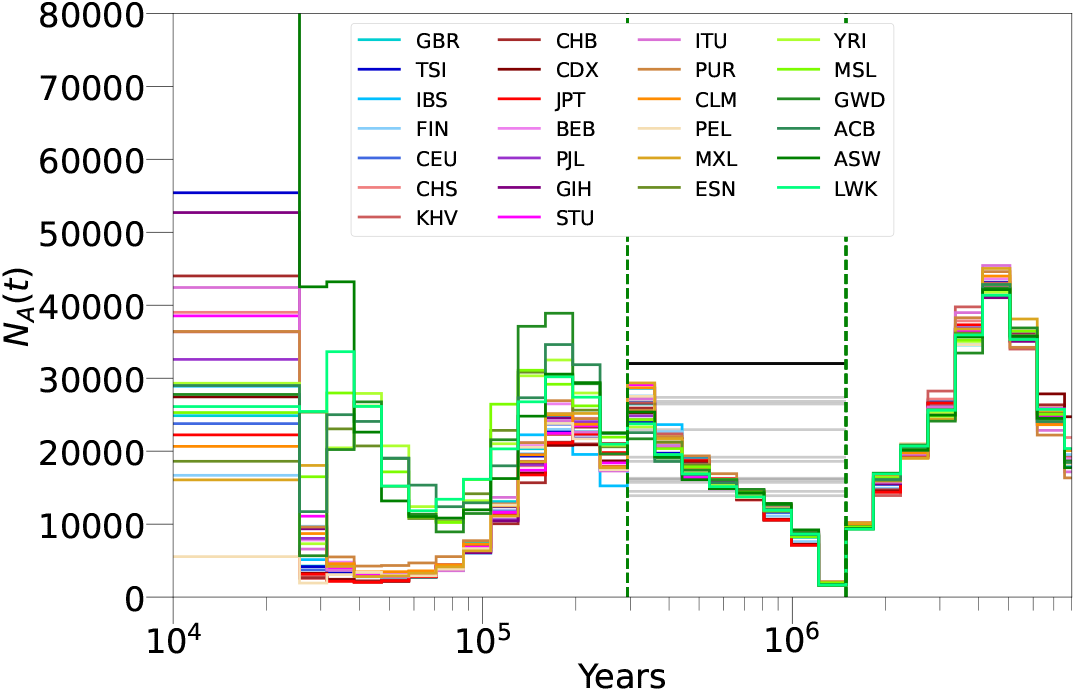
Inference of *N*_*A*_(*t*) from *cobraa* on 26 populations from the 1000 Genomes project, with the inferred size of the ghost population *N*_*B*_(*t*) shown in grey. When optimising we enforced that *N*_*B*_(*t*) must be constant due to identifiabiltiy problems.

**Supplementary Figure 7:**
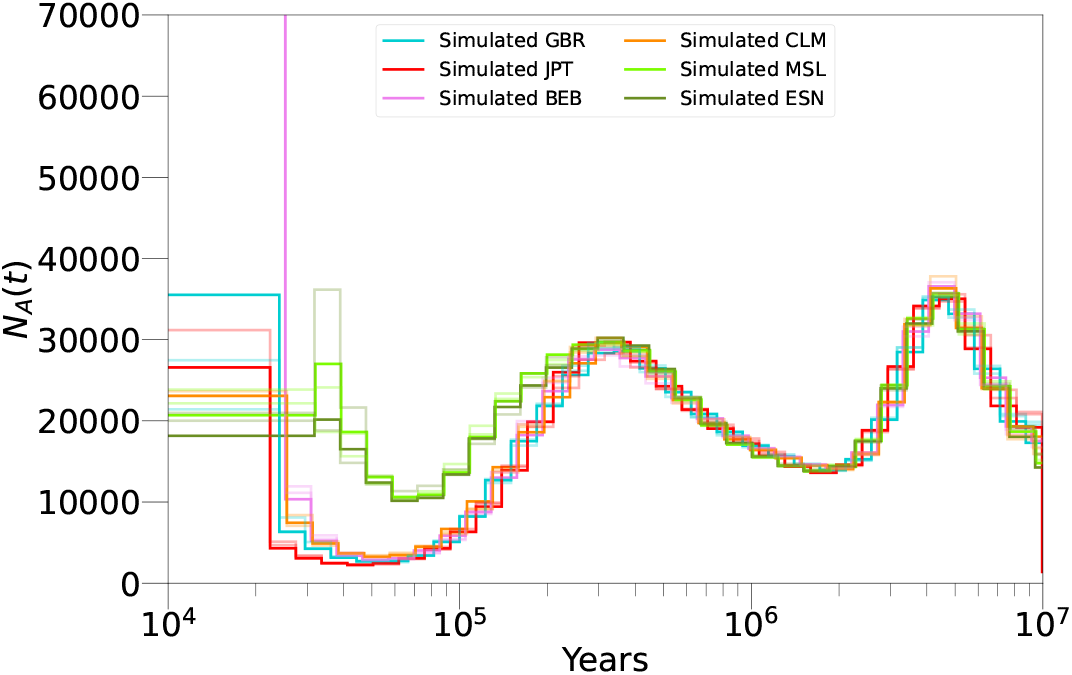
We simulate from the structured model as inferred by *cobraa* (Figures 4b and 4c), then run PSMC on these simulations. The resulting inference, plotted here, looks extremely similar to the PSMC inference on real data (Figure 4a), suggesting the structured model is not necessarily incompatible with previous estimates.

**Supplementary Figure 8:**
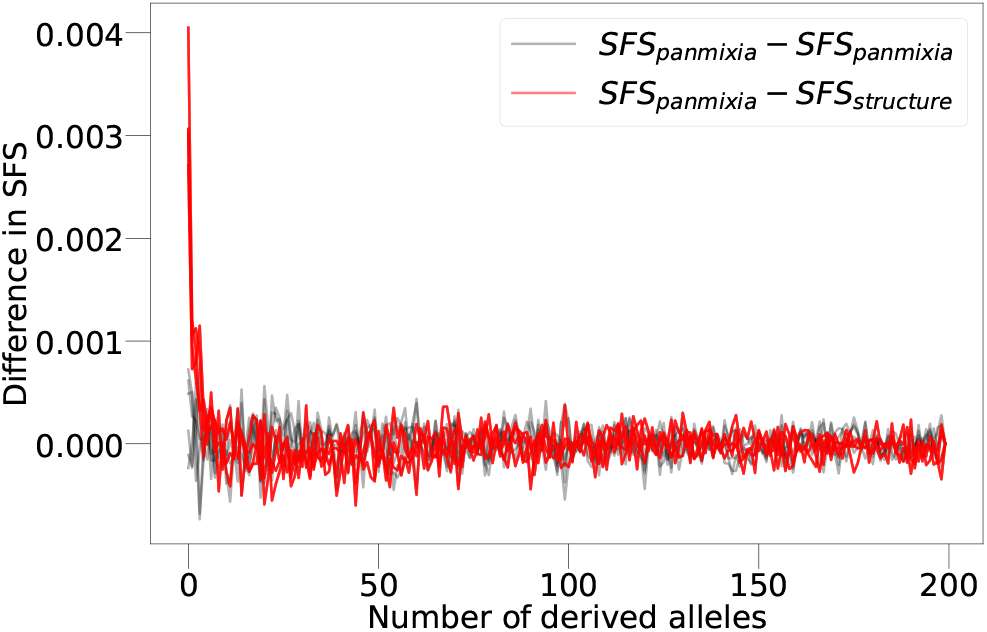
With a panmictic and structured evolutionary history with the same coalescence rate, as shown in Figure 2, we simulated 200 individuals from each model and record the SFS. In red, we plot *SFS*_*panmixia*_ − *SFS*_*structure*_, and see that rare alleles are more common under the panmictic model. In grey we show different replicates of *SFS*_*panmixia*_ − *SFS*_*panmixia*_ to get an estimate of sampling noise. The differences in *SFS*_*panmixia*_ − *SFS*_*structure*_ indicates that the SFS could in principle be used to overcome the pairwise coalescence rate identifiability problem as discused by Mazet et al. [20, 21].

**Supplementary Table 1:** Sequence information for each sample we used in the 1000 Genomes Project. The third column is the number of SNPs that passed all filters. The fourth column is the number of bases that were marked as “missing data”. The fifth column is number of bases that were either homozygous or heterozygous that passed all filters. The sixth column is the average heterozygosity, calculated by dividing the number of heterozygous positions (second column) divided by the number of called bases (fourth column).

| Population | Sample | # SNPs | # Masked bases | # Called bases | Heterozygosity |
| --- | --- | --- | --- | --- | --- |
| GBR | HG00118 | 1617906 | 721916763 | 2150511886 | 0.00075 |
| TSI | NA20752 | 1637719 | 722969288 | 2149265625 | 0.00076 |
| IBS | HG01783 | 1614003 | 731281619 | 2140269189 | 0.00075 |
| FIN | HG00266 | 1619493 | 729407481 | 2142893467 | 0.00075 |
| CEU | NA12718 | 1611666 | 721749959 | 2150584901 | 0.00074 |
| CHS | HG00443 | 1518102 | 720272359 | 2152069281 | 0.0007 |
| KHV | HG02113 | 1508267 | 723553886 | 2148789845 | 0.0007 |
| CHB | NA18530 | 1521984 | 720811264 | 2151278579 | 0.0007 |
| CDX | HG02373 | 1522518 | 716656317 | 2155682074 | 0.0007 |
| JPT | NA18939 | 1503662 | 721812512 | 2150488705 | 0.00069 |
| BEB | HG03006 | 1698194 | 717857271 | 2154406877 | 0.00078 |
| PJL | HG03234 | 1681759 | 719513151 | 2152802272 | 0.00078 |
| GIH | NA20845 | 1686133 | 722230248 | 2143732963 | 0.00078 |
| STU | HG03753 | 1667039 | 728686017 | 2143519959 | 0.00077 |
| ITU | HG03977 | 1657672 | 728197863 | 2144214469 | 0.00077 |
| PUR | HG01171 | 1829263 | 720624146 | 2151746582 | 0.00085 |
| CLM | HG01250 | 1719514 | 717744633 | 2154605688 | 0.00079 |
| PEL | HG02285 | 1482859 | 723765263 | 2148505378 | 0.00069 |
| MXL | NA19648 | 1602318 | 719980307 | 2152287788 | 0.00074 |
| ESN | HG03515 | 2133476 | 723310219 | 2149247137 | 0.00099 |
| YRI | NA18488 | 2144349 | 727775665 | 2144733225 | 0.00099 |
| MSL | HG03212 | 2147856 | 724459558 | 2148121619 | 0.00099 |
| GWD | HG02568 | 2104590 | 733799722 | 2138584844 | 0.00098 |
| ACB | HG01882 | 2171991 | 725229153 | 2147260898 | 0.00101 |
| ASW | NA19625 | 2156788 | 729430494 | 2143090371 | 0.001 |
| LWK | NA19017 | 2175804 | 720231319 | 2152220886 | 0.00101 |

**Supplementary Figure 9:**
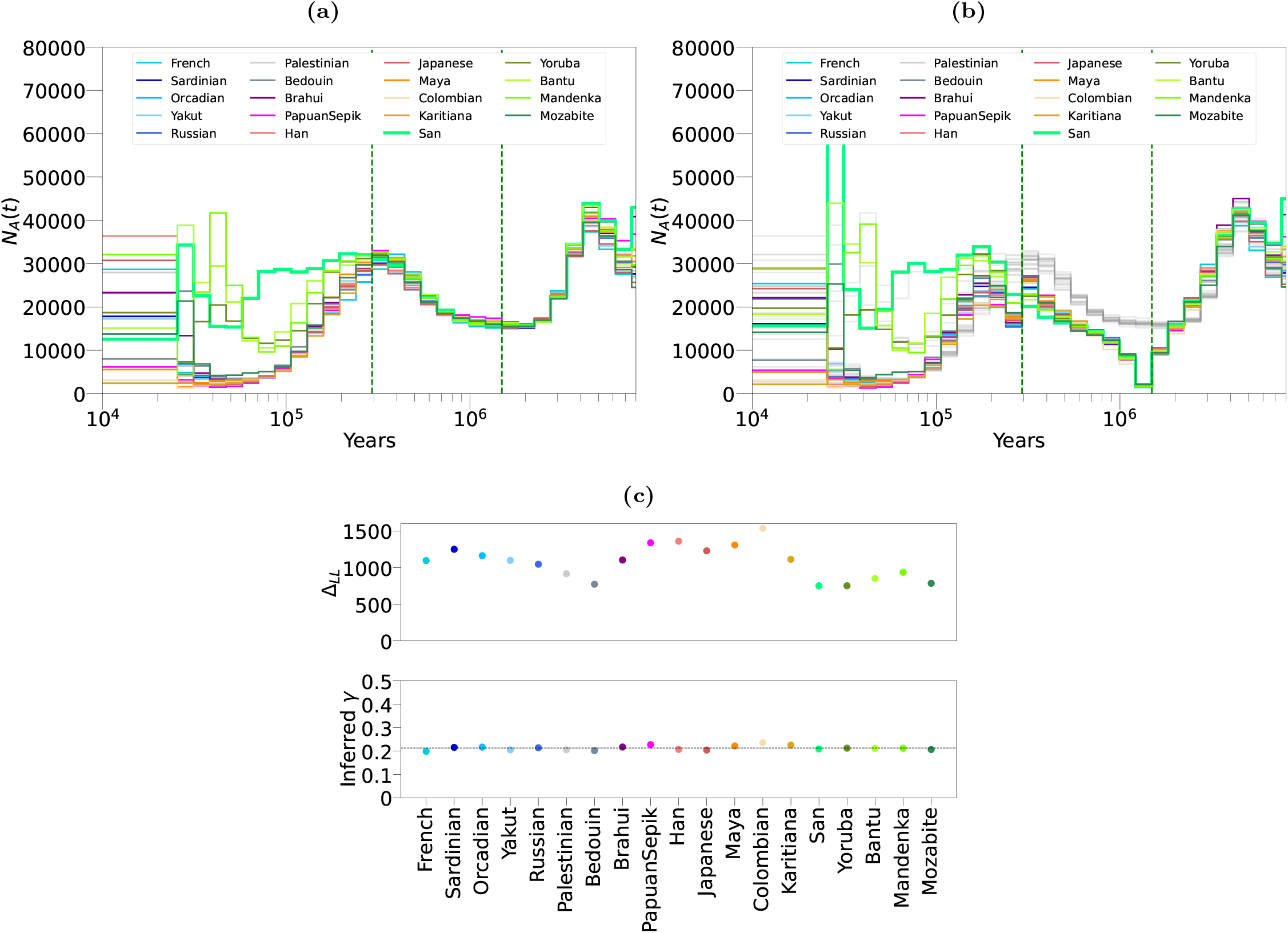
Inference from PSMC and *cobraa* on 19 populations from the Human Genome Diversity Project, using one individual per population, iterating until convergence. **(a)** PSMC’s estimate of *N*_*A*_(*t*). **(b)** *cobraa*’s estimate of *N*_*A*_(*t*), with the estimated split/admixture time shown in vertical, dashed, green lines. For direct comparison, the PSMC inference from (a) is also plotted in grey. **(c)** The top panel shows the difference between the log-likelihood from *cobraa*’s inference and PSMC’s inference, Δ_ℒ_ = ℒ_*S*_ − ℒ_*U*_ ; the bottom panel shows *cobraa*’s inferred admixture fraction *γ*. The results on the HGDP dataset are extremely similar to those as inferred on the 1000GP data (Figure 4).

**Supplementary Figure 10:**
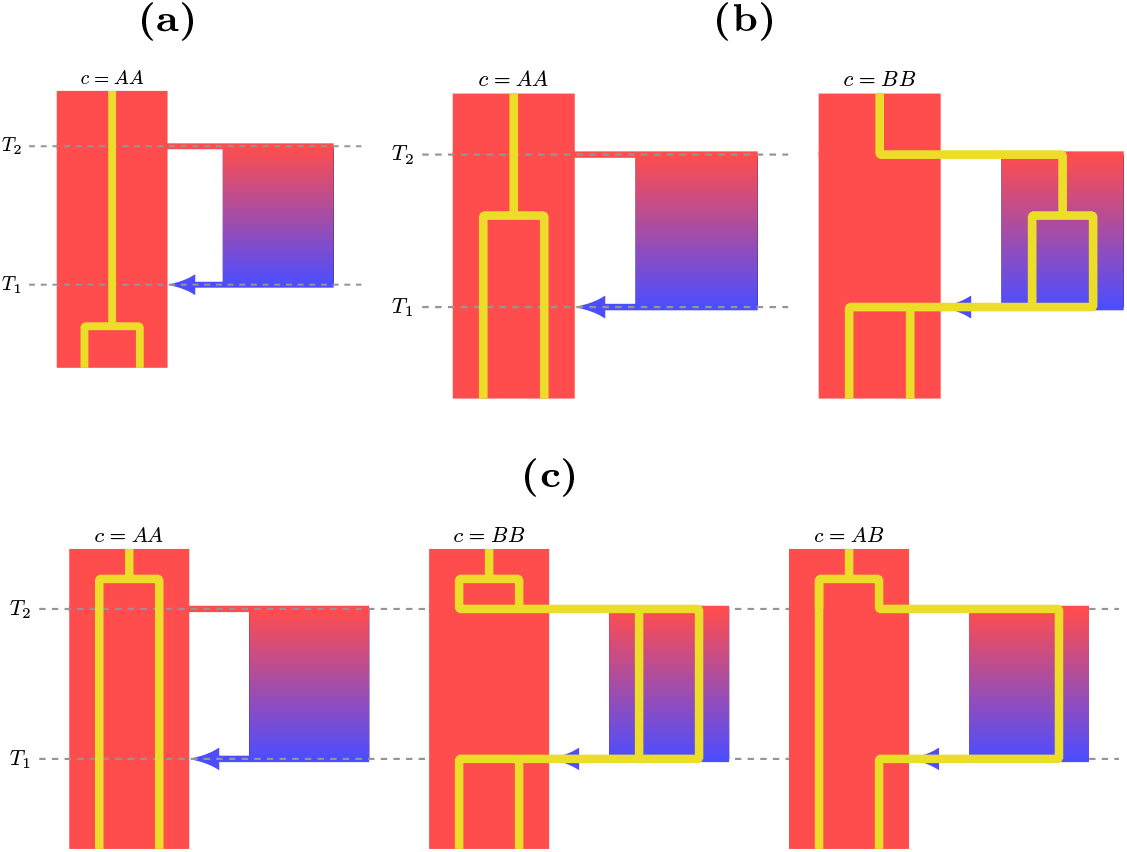
A diagram of the possible ancestral lineage paths, *c*, as explicitly modelled in *cobraa-path*. **(a)** If the coalescence time was more recent than the admixture event (*T*_1_), the only possibility is that the lineages coalesce in population *A*, which we denote as *c* = *AA*. **(b)** If the coalescence time was in the structured period, between *T*_1_ and *T*_2_, then *c* = *AA* or *c* = *BB*. **(c)** If the coalescence time was more ancient than the split time, *T*_1_, then *c* = *AA, c* = *BB* or *c* = *AB*.

**Supplementary Figure 11:**
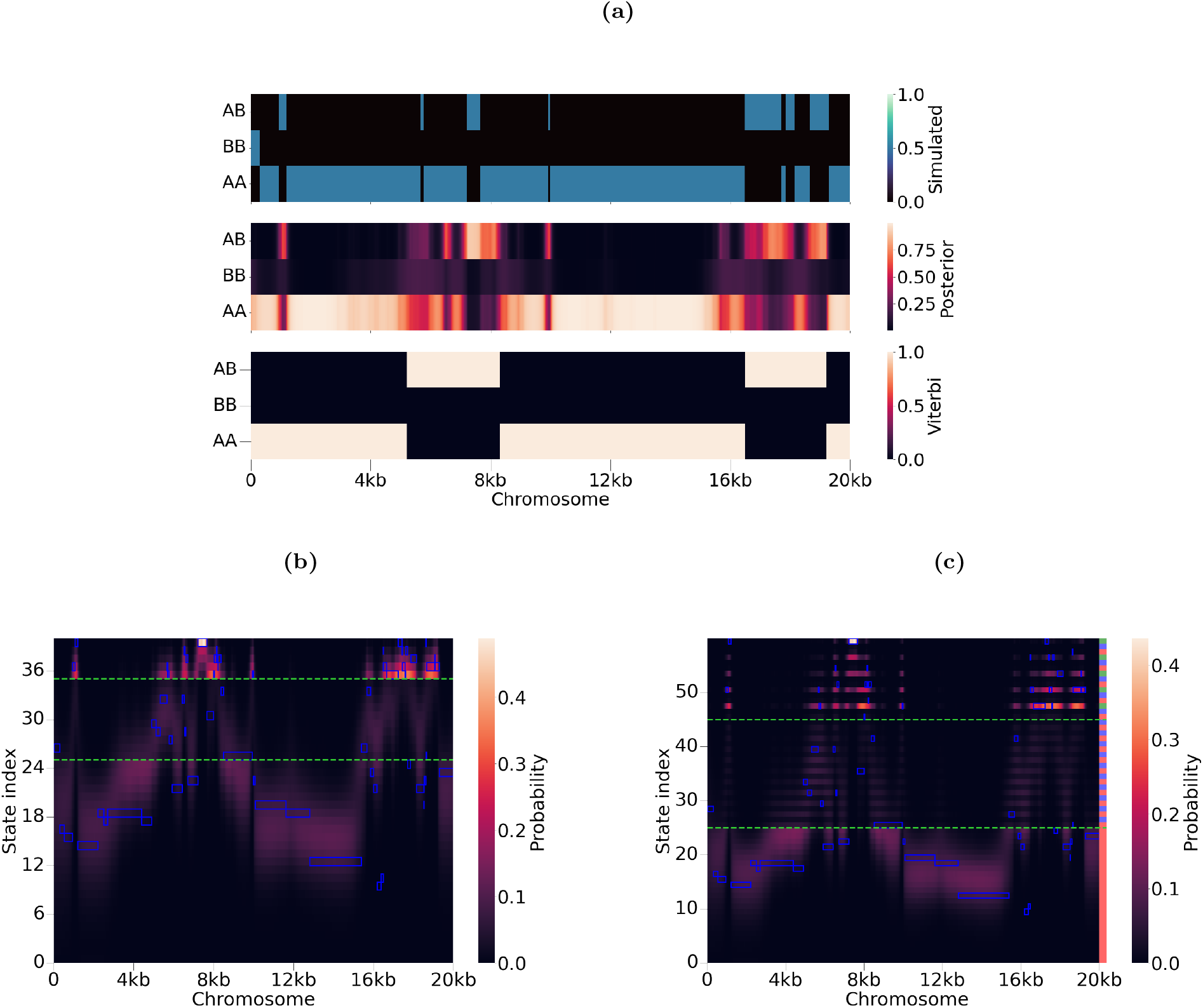
We can decode the HMM of *cobraa-path* to infer the admixed regions of the genome (i.e. parts of the genome where the ancestral lineage pair went through *AB* or *BB*). The top panel of **(a)** shows the simulated lineage path across the genome, where the x-axis indicates the chromosomal position and the y-axis the ancestral lineage path. A structured model was simulated with *µ/r* = 1.25, 40% admixture, and constant population sizes. Using the simulated structured parameters, the middle panel shows the marginal posterior probability of each lineage path (from the forward/backward algorithm), and the bottom shows the most likely lineage path (from the Viterbi algorithm). **(b)** The full *cobraa* decoding of the simulation (hidden states are discretised coalescence times), where the y-axis indicates the coalescence time with 0 being the present. The green, dashed, horizontal lines indicates the simulated split and rejoin times. **(c)** The full *cobraa-path* decoding of the simulation (hidden states are discretised coalescence times and ancestral lineage path). The y-axis indicates not only the coalescence time but also the ancestral lineage path, which is indicated by the shading in the right-most column. Red indicates *AA*, blue *BB*, and green *AB*. In the structured period, a red-blue pair indicates the same coalesence time, and more anciently than the split time a red-blue-green triple indicate the same coalescence time. The simulated states through the model are shown by the highlighed blue cells, and the posterior probabilities are indicated by the shading of the heatmap, with cream representing total confidence and black indicating no confidence. The y-axis in (a) indicates the ancestral lineage path, and in (b) and (c) the coalescence time, with more ancient states at the top and more recent at the bottom. The horizontal, green, dashed lines in (b) and (c) indicate the split and rejoin time.

**Supplementary Figure 12:**
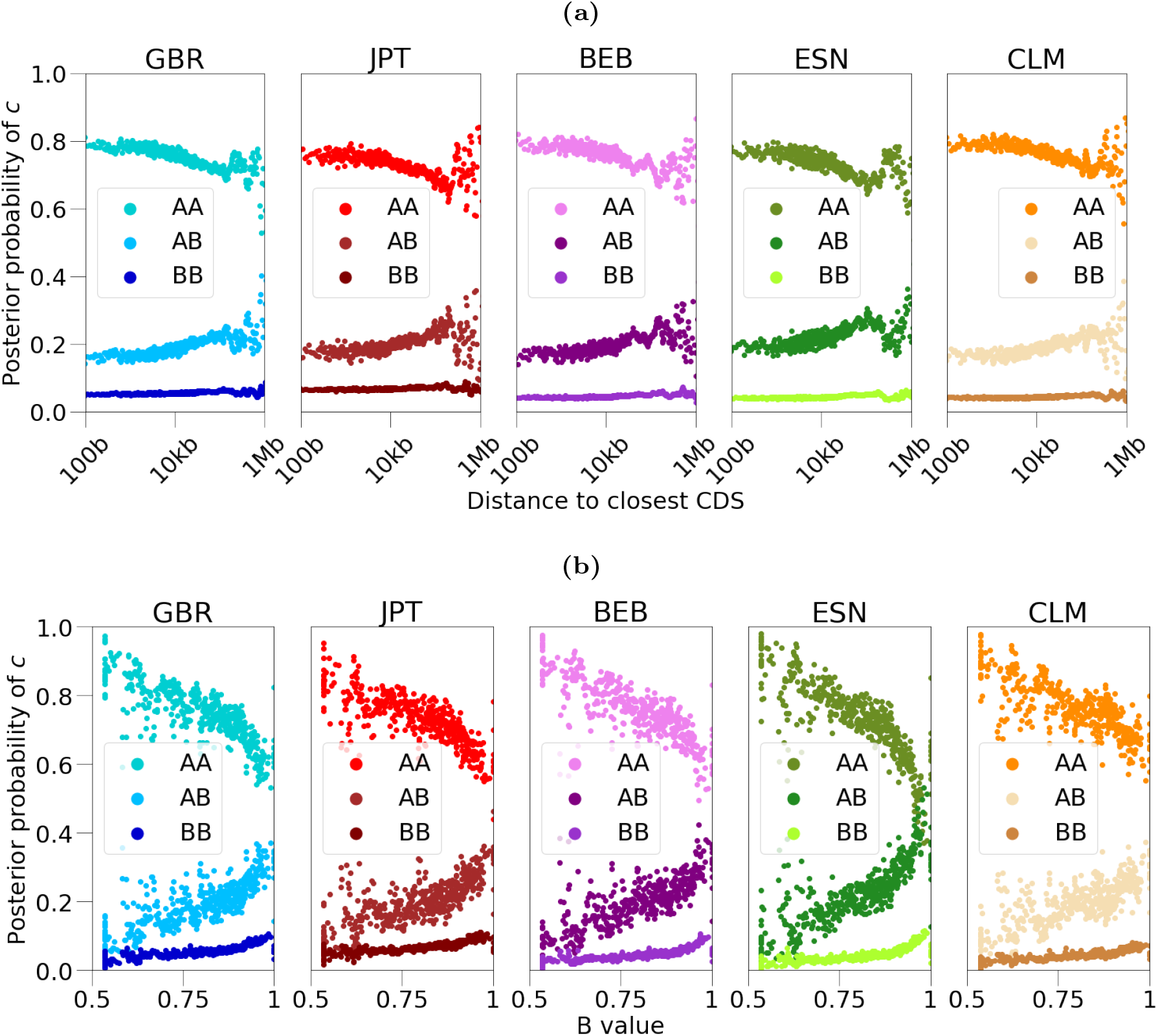
Correlation between the inferred probability of an admixed lineage path (conditional on the coalescence time being older than the admixture event) and the distance to coding sequence (CDS) **(a)** or strength of B-value **(b)**. To get the inferred probability of an admixed lineage path, we decode the HMM of *cobraa-path* with the parameters as inferred from *cobraa* (Figures 4b and 4c), and marginalise out the coalescence times. The Spearman correlation coefficients for each population are shown in Table S2.

**Supplementary Figure 13:**
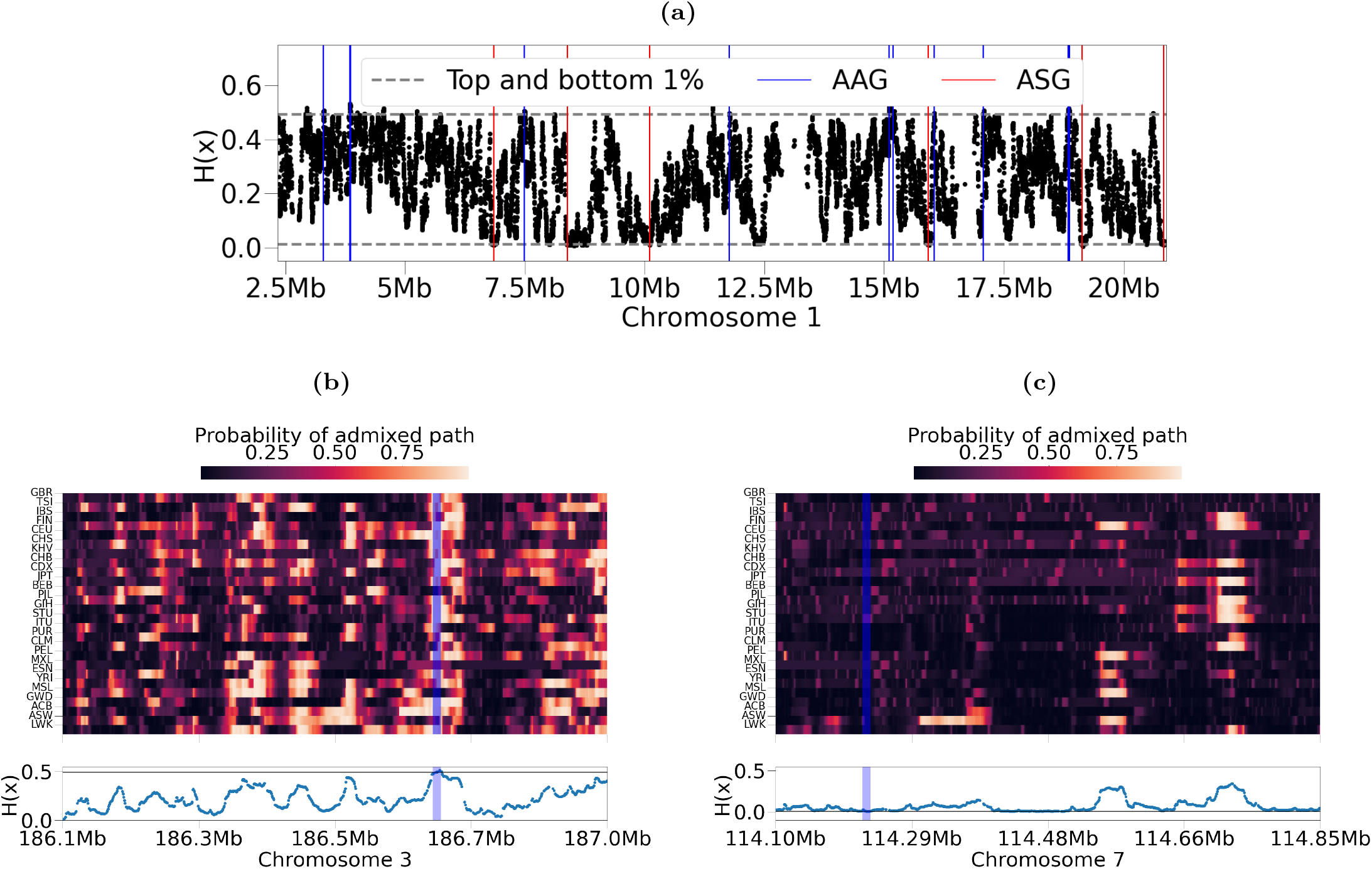
*cobraa-path* can be used to infer the probability that one or both of the lineages went through the ghost population. Using these probabilities, we defined *H*(*x*) to calculate the expected amount of admixture across all populations at each position. **(a)** An illustration of *H*(*x*) across chromosome 1 (black dots), with the top and bottom 1% threshold indicated in the grey dashed line. Protein coding genes overlapping with the top or bottom 1% of *H*(*x*) (AAGs and ASGs) are shown in blue and red, respectively, which illustrates that many regions exceeding the threshold values do not overlap with protein coding genes. Plotted values of *H*(*x*) are shown with points, rather than a continuous line due to missing data causing discontinuity. **(b)** The top panel shows the probability of admixture for each population, on chromosome 3 around the gene KNG1 (highlighted in blue), which is one of the 666 protein coding genes overlapping the regions in the top 1% of *H*(*x*). The probability of admixture for population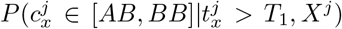, where *x* is the genomic position, *t* is the coalescence time at *x, c* is the ancestral lineage path at *x*, and *X* is all the observed data. The bottom panel shows the corresponding value of *H*(*x*). **(c)** The top panel shows the probability of admixture for each population, on chromosome 7 around the gene FOXP2 (highlighted in blue), which is one of the 1270 protein coding genes overlapping the regions in the bottom 1% of *H*(*x*). The bottom panel shows the corresponding value of *H*(*x*).

**Supplementary Figure 14:**
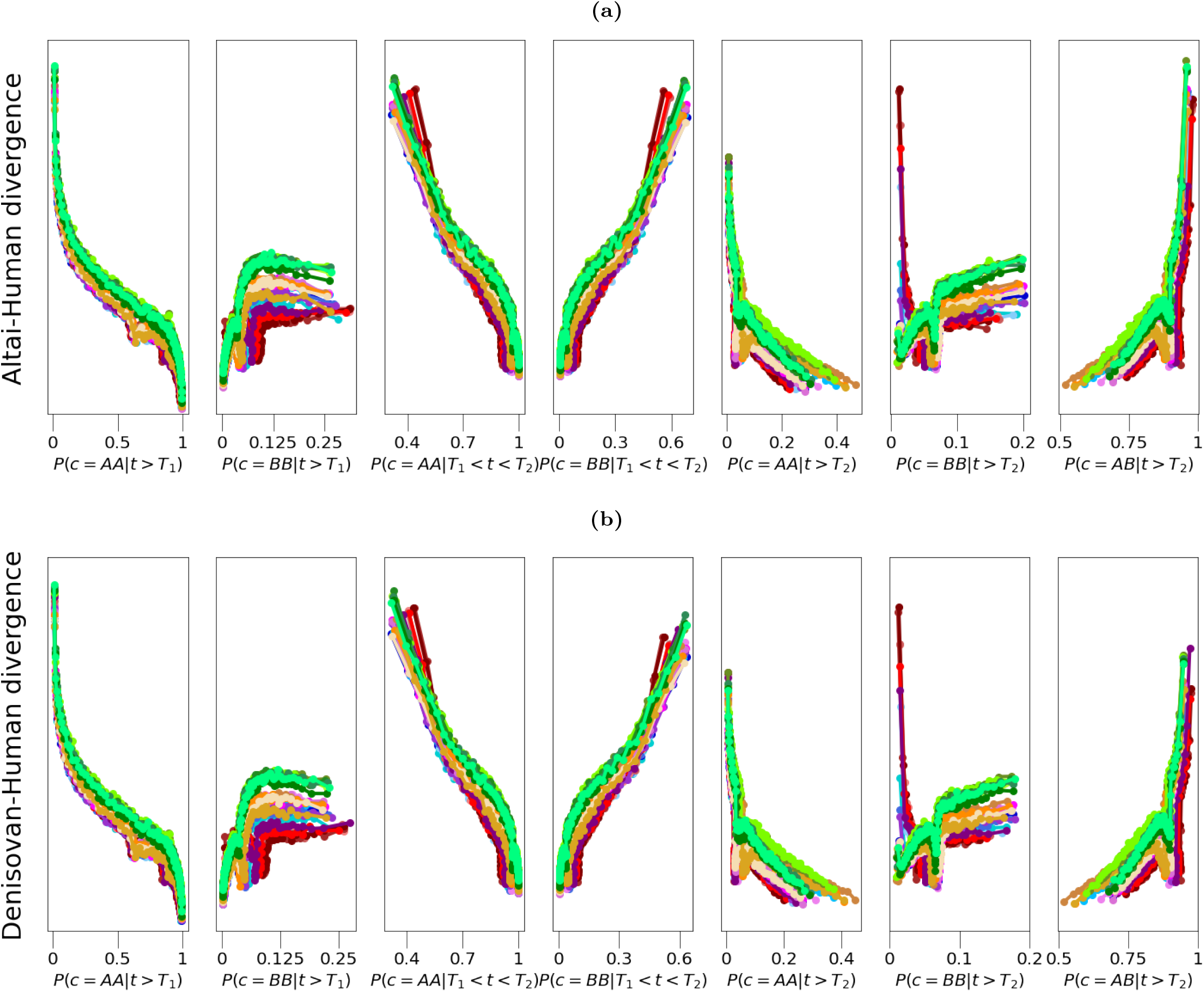
Relationship between the probability of ancestral lineage path, and **(a)** human-Neanderthal or **(b)** human-Denisovan divergence, for all 1000GP populations. The different columns represent ancestral lineage path probabilities with different conditioning on coalescence time. Because *P* (*c* = *AA*|*T*_1_ *< t < T*_2_) = 1 − *P* (*c* = *BB*|*T*_1_ *< t < T*_2_), the third and fourth plots on each row are a vertical mirrors of each other. All Spearman correlations are reported in Table S6.

**Supplementary Figure 15:**
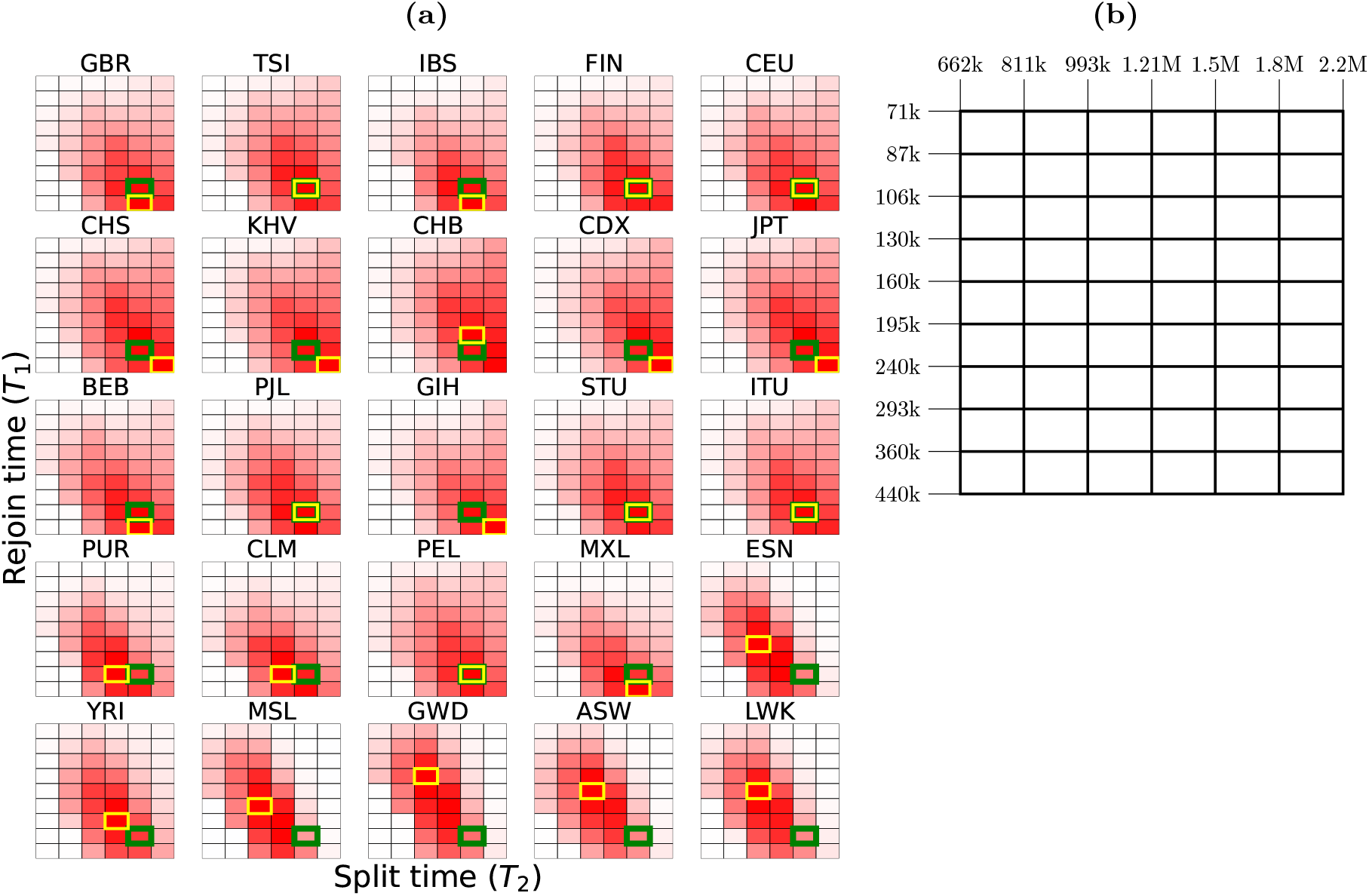
**(a)** The log-likelihood (L) difference between *cobraa* and PSMC, for all given pairings of *T*_1_ and *T*_2_ for each population (ACB excluded). Each algorithm was run until convergence, defined as the change in ℒ between subsequent iterations of the EM algorithm being less than one. Red indicates a positive difference, and white zero. Each population is shown on its own scale. The green highlighted cell indicates the composite maximum likelihood (CML) estimate and the yellow highlighted cell indicates the maximum likelihood estimate per population. Higher on the y-axis indicates more recent *T*_1_, and leftmost on the x-axis indicates more recent *T*_2_. **(b)** A diagram indicating the corresponding time interval boundaries associated with each cell in **a)**.

**Supplementary Figure 16:**
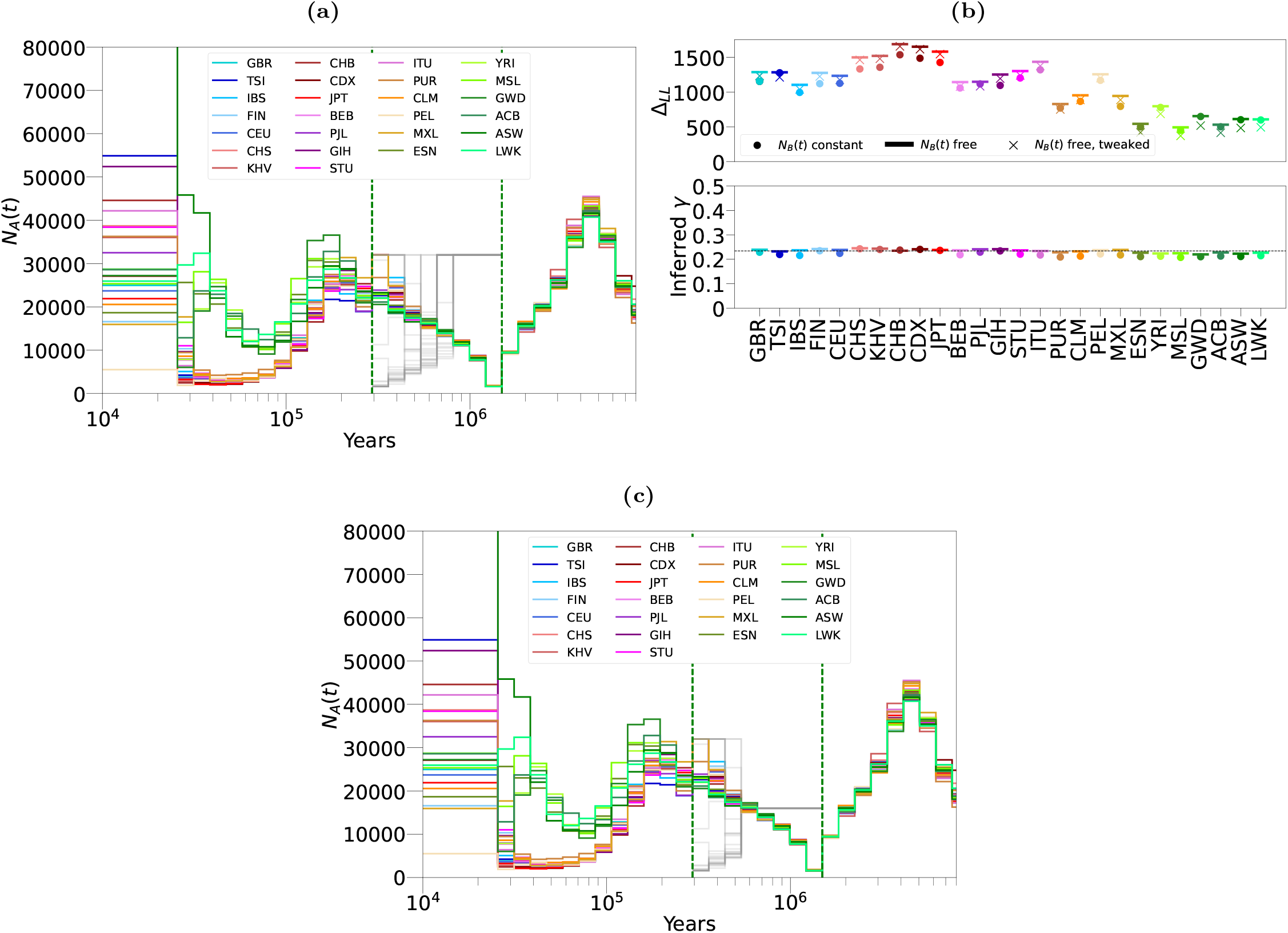
Inference from *cobraa* on 26 populations from the 1000 Genomes Project, with *N*_*B*_(*t*) allowed to vary (contrary to Figure 4 in which *N*_*B*_(*t*) was forced to be constant). **(a)** Estimates of *N*_*A*_(*t*) (coloured lines) and *N*_*B*_(*t*) (grey lines). **(b)** The difference in log-likelihood from *cobraa*’s and PSMC’s inference, Δ_ℒ_ = ℒ_*S*_ − ℒ_*U*_ (top panel), and the inferred admixture fraction (bottom panel). The circles indicate Δ_*L*_ and the inferred *γ* from the the model where *N*_*B*_(*t*) was enforced to be constant (i.e. are exactly the same as Figure 4c), the horizontal lines indicate the inference with *N*_*B*_(*t*) allowed to vary, and the cross indicates the log-likelihood of the free *N*_*B*_(*t*) model after the size of *N*_*B*_(*t*) is reduced post divergence, as depicted in **(c)**.

**Supplementary Figure 17:**
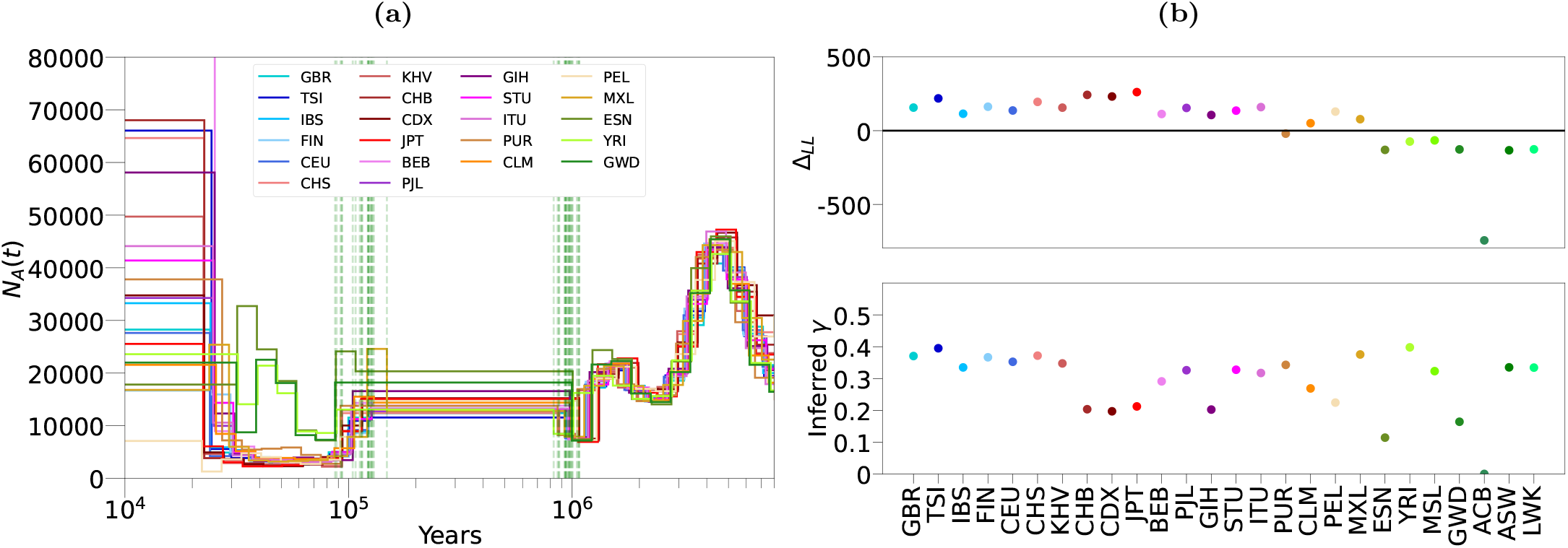
Inference from *cobraa* on the 1KGP populations, when we enforce that *N*_*A*_(*t*) must be constant in the structured period. **(a)** Inference of *N*_*A*_(*t*) and the split/rejoin times, **(b)** Estimates of the admixture fraction (top) and the difference in log-likelihood between *cobraa* and PSMC (bottom). We conclude a structured model with this constraint is not well supported by the data, as Δ_ℒ_ is often negative and the estimates of the admixture fraction have high variance.

**Supplementary Figure 18:**
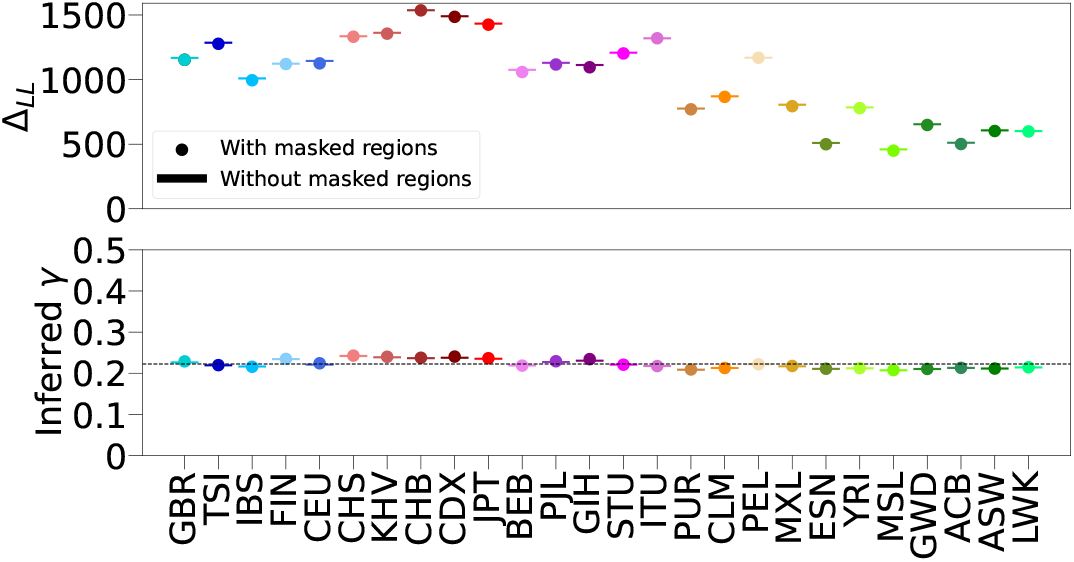
Missing data in the HMM does not make a difference for inference. We reran PSMC and *cobraa* analysis after removing missing data from the observations (see Methods), and found the results to be indistinguishable. The circles indicate Δ_*L*_ and the inferred *γ* from the full data as (i.e. are exactly the same as Figure 4c), and the horizontal lines indicate the analysis after the missing data was removed.

**Supplementary Figure 19:**
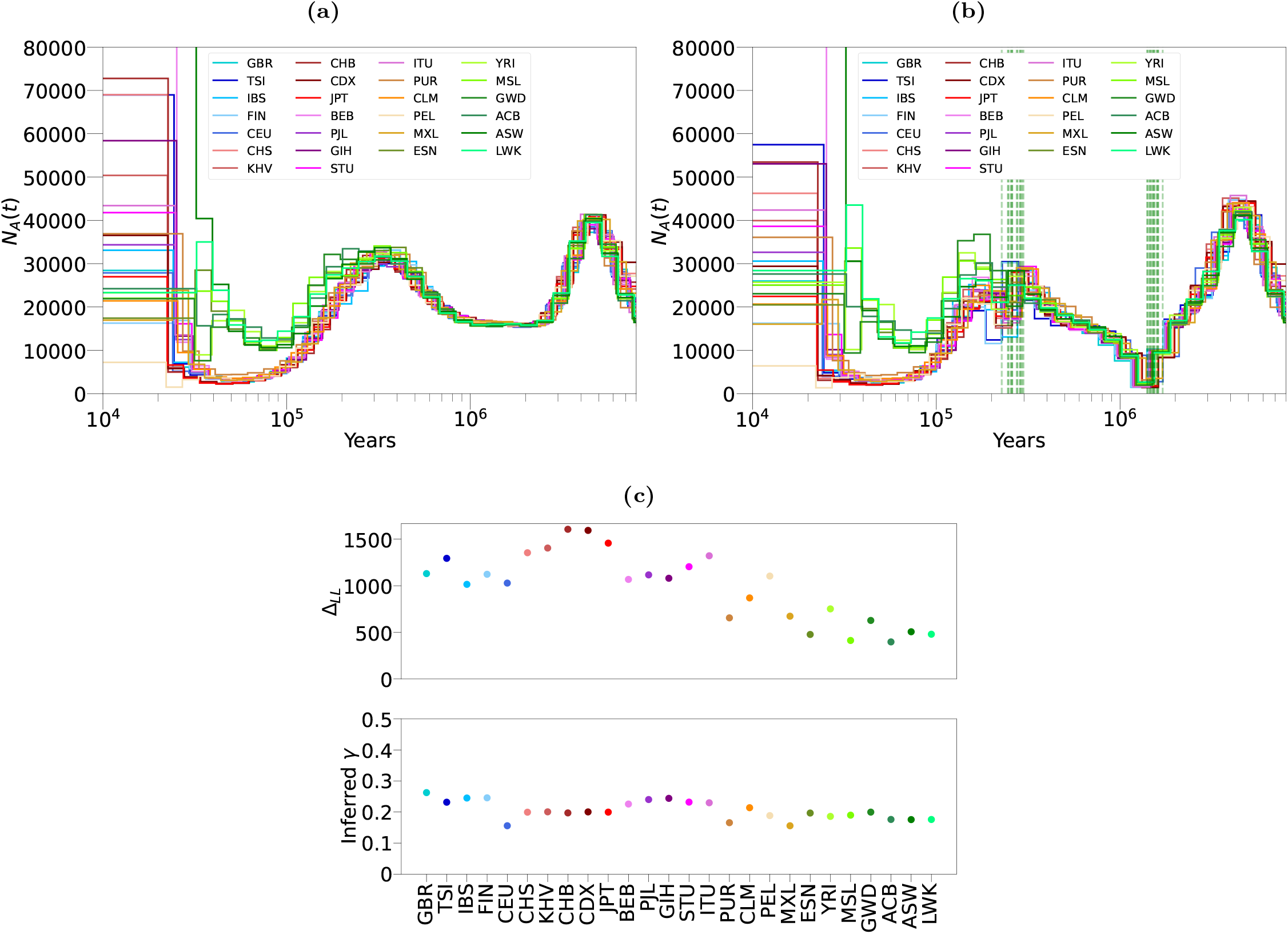
Inference from PSMC and *cobraa* on 26 populations from the 1000 Genomes project, using one individual per population, with *θ* inferred from the data. **(a)** PSMC’s estimate of *N*_*A*_(*t*). *cobraa*’s estimate of *N*_*A*_(*t*), with the estimated split/admixture time shown in vertical, dashed, green lines. **(c)** The top panel shows the difference between the log-likelihood from *cobraa*’s inference and PSMC’s inference, Δ_ℒ_ = ℒ_*S*_ − ℒ_*U*_ ; the bottom panel shows *cobraa*’s inferred admixture fraction *γ*.

**Supplementary Figure 20:**
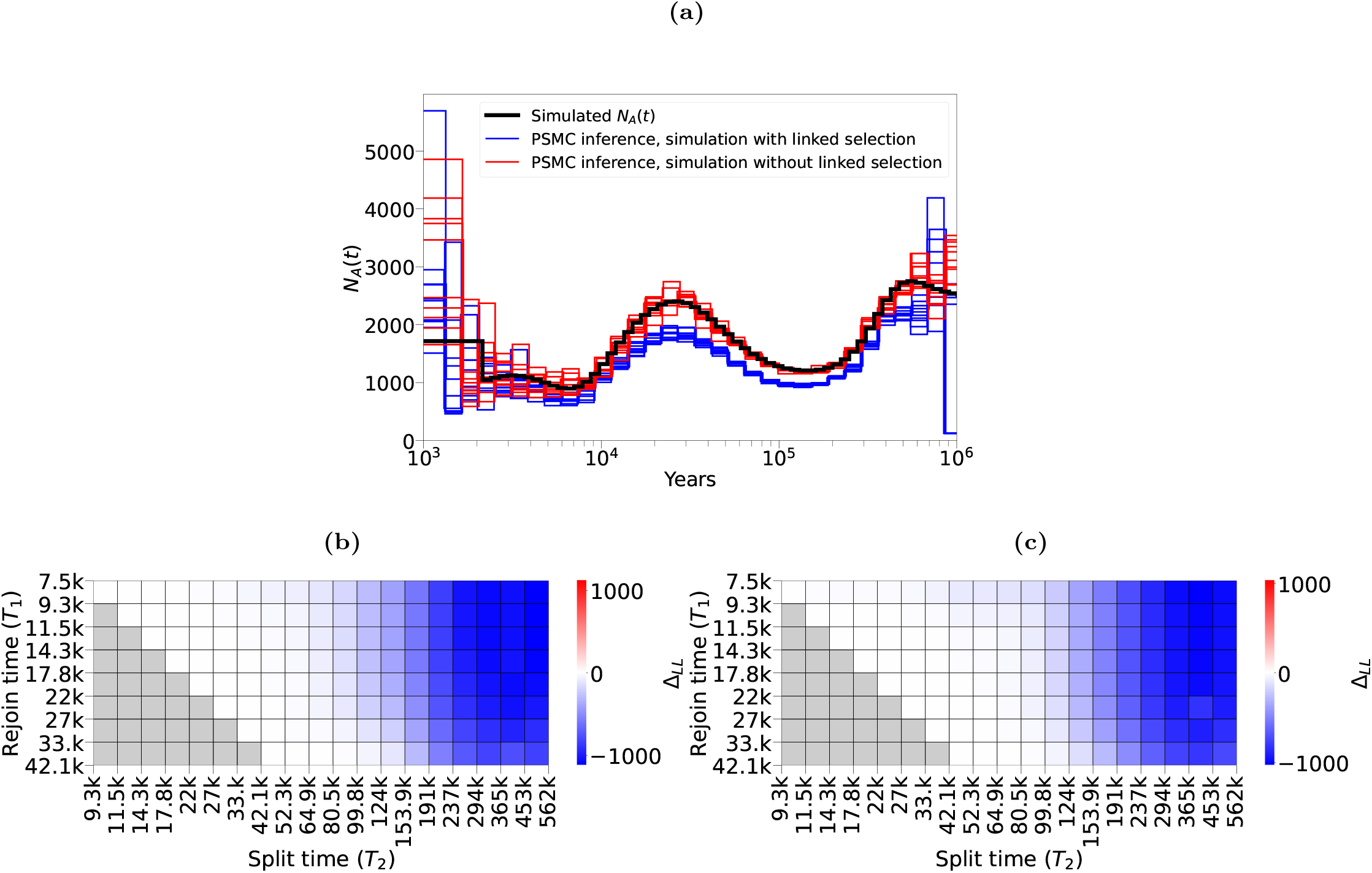
Inference from PSMC and *cobraa* on simulations with and without widespread linked selection, taken from [70] (see Methods). **(a)** PSMC inference on a simulation with and without widespread linked selection. We ran *cobraa* over various pairings of *T*_1_ and *T*_2_, and calculated the loglikelihood difference between this and the inference from PSMC, 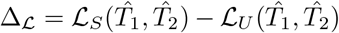,In **(b)** and, we show Δ_ℒ_ for a neutral simulation and simulation with widespread linked selection, respectively. The log-likelihood differences indicate that *cobraa* does not falsely infer linked selection as structure.

**Supplementary Table 2:** Spearman correlation between the inferred probability of an admixed lineage path, *Pr*(*c* = *k*|*X, t > T*_1_), and the distance to CDS (dcCDS) or B-value, for each population on chromosome 1. The associated p-value (testing for uncorrelated datasets generating a Spearman coefficient equal to or more extreme than the one reported) is less than 1e-100 for every correlation in each population.

| Population | dcCDS vs $Pr(c = k\vert X, t > T_1)$ | | | B-value vs $Pr(c = k\vert X, t > T_1)$ | | |
| --- | --- | --- | --- | --- | --- | --- |
| | $k = AA$ | $k = AB$ | $k = BB$ | $k = AA$ | $k = AB$ | $k = BB$ |
| GBR | -0.09 | 0.07 | 0.07 | -0.27 | 0.18 | 0.3 |
| TSI | -0.09 | 0.07 | 0.09 | -0.28 | 0.2 | 0.35 |
| IBS | -0.09 | 0.07 | 0.08 | -0.24 | 0.18 | 0.29 |
| FIN | -0.09 | 0.07 | 0.05 | -0.25 | 0.16 | 0.25 |
| CEU | -0.09 | 0.07 | 0.08 | -0.26 | 0.18 | 0.32 |
| CHS | -0.08 | 0.06 | 0.04 | -0.28 | 0.19 | 0.18 |
| KHV | -0.05 | 0.04 | 0.01 | -0.25 | 0.16 | 0.19 |
| CHB | -0.06 | 0.04 | 0.01 | -0.25 | 0.16 | 0.16 |
| CDX | -0.1 | 0.08 | 0.03 | -0.27 | 0.17 | 0.2 |
| JPT | -0.09 | 0.07 | 0.05 | -0.25 | 0.16 | 0.22 |
| BEB | -0.09 | 0.07 | 0.09 | -0.25 | 0.19 | 0.31 |
| PJL | -0.1 | 0.09 | 0.08 | -0.29 | 0.21 | 0.35 |
| GIH | -0.09 | 0.08 | 0.04 | -0.27 | 0.18 | 0.25 |
| STU | -0.1 | 0.08 | 0.11 | -0.27 | 0.2 | 0.33 |
| ITU | -0.09 | 0.07 | 0.09 | -0.24 | 0.17 | 0.31 |
| PUR | -0.1 | 0.08 | 0.1 | -0.27 | 0.2 | 0.32 |
| CLM | -0.07 | 0.06 | 0.07 | -0.24 | 0.17 | 0.29 |
| PEL | -0.07 | 0.06 | 0.07 | -0.24 | 0.17 | 0.31 |
| MXL | -0.07 | 0.07 | 0.05 | -0.25 | 0.18 | 0.31 |
| ESN | -0.1 | 0.08 | 0.08 | -0.33 | 0.28 | 0.38 |
| YRI | -0.12 | 0.11 | 0.11 | -0.31 | 0.25 | 0.37 |
| MSL | -0.11 | 0.1 | 0.11 | -0.33 | 0.27 | 0.37 |
| GWD | -0.1 | 0.09 | 0.08 | -0.32 | 0.26 | 0.38 |
| ACB | -0.11 | 0.09 | 0.1 | -0.34 | 0.28 | 0.4 |
| ASW | -0.12 | 0.1 | 0.12 | -0.32 | 0.26 | 0.38 |
| LWK | -0.11 | 0.1 | 0.1 | -0.31 | 0.26 | 0.36 |
| mean | -0.097 | 0.081 | 0.078 | -0.28 | 0.21 | 0.31 |

**Supplementary Table 3:** Biological processes associated with the 666 admixture abundant genes (AAGs). First column is the reported biological process reported from Panther’s [49] analysis, second column is number of human genes associated with this process (out of 20,592), third column is number of AAGs associated with this process, fourth column is expected number of genes assuming a random sampling, fifth column is the enrichment with respect to the number of expected genes, sixth column is the p-value from Fisher’s Exact Test, and the seventh column is the Benjamini-Hochberg False Discovery Rate. Full information including parent processes are available for download online.

| Biological function | # Human genes | # AAGs | Expected | Enrichment | p-value | FDR |
| --- | --- | --- | --- | --- | --- | --- |
| Neuron cell-cell adhesion | 16 | 6 | 0.52 | 11.63 | 4.42e-5 | 4.21e-2 |
| Startle response | 30 | 8 | 0.97 | 8.27 | 1.85e-5 | 3.53e-2 |
| Neuron recognition | 45 | 9 | 1.45 | 6.20 | 4.06e-5 | 4.12e-2 |
| Neurotransmitter transport | 142 | 17 | 4.58 | 3.71 | 1.04e-5 | 2.27e-2 |
| Extracellular matrix organization | 278 | 24 | 8.96 | 2.68 | 3.00e-5 | 4.16e-2 |
| Chemical synaptic transmission | 420 | 34 | 13.54 | 2.51 | 3.12e-6 | 1.19e-2 |
| Circulatory system process | 508 | 36 | 16.38 | 2.20 | 2.58e-5 | 3.93e-2 |
| Regulation of biological quality | 2849 | 132 | 91.87 | 1.44 | 2.48e-5 | 4.20e-2 |
| Gene expression | 2491 | 44 | 80.32 | 0.55 | 6.04e-6 | 1.53e-2 |

**Supplementary Table 4:** Biological processes associated with the 1270 admixture scarce genes (ASGs). First column is the reported biological process reported from Panther’s [49] analysis, second column is number of human genes associated with this process (out of 20,592), third column is number of ASGs associated with this process, fourth column is expected number of genes assuming a random sampling, fifth column is the enrichment with respect to the number of expected genes, sixth column is the p-value from Fisher’s Exact Test, and the seventh column is the Benjamini-Hochberg False Discovery Rate. Full information including parent processes are available for download online.

| Biological function | # Human genes | # ASGs | Expected | Enrichment | p-value | FDR |
| --- | --- | --- | --- | --- | --- | --- |
| Pre-miRNA processing | 14 | 7 | 0.85 | 8.28 | 1.10e-4 | 1.11e-2 |
| Cortical actin cytoskeleton organization | 41 | 10 | 2.48 | 4.04 | 5.22e-4 | 4.12e-2 |
| Golgi to plasma membrane transport | 50 | 11 | 3.02 | 3.64 | 5.91e-4 | 4.62e-2 |
| Regulation of intracellular steroid hormone receptor signaling pathway | 74 | 15 | 4.47 | 3.36 | 1.42e-4 | 1.37e-2 |
| Homophilic cell adhesion via plasma membrane adhesion molecules | 167 | 33 | 10.09 | 3.27 | 3.37e-8 | 9.02e-6 |
| Regulation of embryonic development | 92 | 16 | 5.56 | 2.88 | 4.00e-4 | 3.33e-2 |
| Protein deubiquitination | 109 | 18 | 6.58 | 2.73 | 3.13e-4 | 2.73e-2 |
| Peptidyl-serine phosphorylation | 163 | 26 | 9.85 | 2.64 | 4.23e-5 | 5.16e-3 |
| Cytosolic transport | 133 | 21 | 8.03 | 2.61 | 2.95e-4 | 2.58e-2 |
| Protein dephosphorylation | 137 | 21 | 8.28 | 2.54 | 3.48e-4 | 2.97e-2 |
| Mesenchymal cell differentiation | 172 | 24 | 10.39 | 2.31 | 4.66e-4 | 3.80e-2 |
| Vesicle localization | 181 | 25 | 10.93 | 2.29 | 3.70e-4 | 3.10e-2 |
| Proteasome-mediated ubiquitin-dependent protein catabolic process | 297 | 41 | 17.94 | 2.29 | 4.72e-6 | 8.00e-4 |
| Endosomal transport | 265 | 34 | 16.01 | 2.12 | 1.26e-4 | 1.23e-2 |
| Chromatin remodeling | 621 | 79 | 37.52 | 2.11 | 6.13e-9 | 1.95e-6 |
| Regulation of GTPase activity | 298 | 37 | 18.00 | 2.06 | 1.15e-4 | 1.15e-2 |
| Cell division | 525 | 60 | 31.72 | 1.89 | 1.12e-5 | 1.65e-3 |
| Regulation of neuron projection development | 459 | 52 | 27.73 | 1.88 | 6.14e-5 | 6.89e-3 |
| Regulation of neuron projection development | 459 | 52 | 27.73 | 1.88 | 6.14e-5 | 6.89e-3 |
| Positive regulation of cell projection organization | 358 | 40 | 21.63 | 1.85 | 6.18e-4 | 4.73e-2 |
| DNA repair | 519 | 57 | 31.35 | 1.82 | 5.16e-5 | 6.05e-3 |
| Protein ubiquitination | 625 | 67 | 37.76 | 1.77 | 2.59e-5 | 3.46e-3 |
| Positive regulation of catabolic process | 523 | 56 | 31.60 | 1.77 | 1.17e-4 | 1.16e-2 |
| Regulation of cell cycle phase transition | 434 | 46 | 26.22 | 1.75 | 6.31e-4 | 4.81e-2 |
| Neuron projection morphogenesis | 491 | 51 | 29.66 | 1.72 | 4.64e-4 | 3.80e-2 |
| MRNA metabolic process | 622 | 64 | 37.58 | 1.70 | 1.05e-4 | 1.07e-2 |
| DNA-templated transcription | 539 | 55 | 32.56 | 1.69 | 4.20e-4 | 3.46e-2 |
| Cell cycle process | 880 | 89 | 53.16 | 1.67 | 9.52e-6 | 1.44e-3 |
| Brain development | 742 | 75 | 44.83 | 1.67 | 4.17e-5 | 5.21e-3 |
| Heart development | 566 | 57 | 34.19 | 1.67 | 4.17e-4 | 3.45e-2 |
| Mitotic cell cycle | 594 | 59 | 35.88 | 1.64 | 5.54e-4 | 4.35e-2 |
| Microtubule-based process | 824 | 81 | 49.78 | 1.63 | 5.15e-5 | 6.08e-3 |
| Regulation of anatomical structure morphogenesis | 845 | 82 | 51.05 | 1.61 | 8.33e-5 | 8.58e-3 |
| Protein localization to organelle | 739 | 71 | 44.64 | 1.59 | 3.50e-4 | 2.97e-2 |
| Protein transport | 1090 | 103 | 65.85 | 1.56 | 2.09e-5 | 2.84e-3 |
| Positive regulation of cell differentiation | 871 | 82 | 52.62 | 1.56 | 1.89e-4 | 1.76e-2 |
| Regulation of cellular component biogenesis | 974 | 89 | 58.84 | 1.51 | 2.39e-4 | 2.16e-2 |
| Intracellular signal transduction | 1514 | 136 | 91.46 | 1.49 | 1.17e-5 | 1.71e-3 |
| Negative regulation of DNA-templated transcription | 1279 | 112 | 77.27 | 1.45 | 2.00e-4 | 1.84e-2 |
| Regulation of protein modification process | 1230 | 106 | 74.31 | 1.43 | 4.98e-4 | 4.00e-2 |
| Positive regulation of DNA-templated transcription | 1711 | 147 | 103.36 | 1.42 | 4.22e-5 | 5.19e-3 |
| Regulation of intracellular signal transduction | 1781 | 151 | 107.59 | 1.40 | 5.78e-5 | 6.63e-3 |
| Negative regulation of response to stimulus | 1664 | 140 | 100.53 | 1.39 | 1.60e-4 | 1.52e-2 |
| Positive regulation of signal transduction | 1568 | 130 | 94.73 | 1.37 | 4.73e-4 | 3.84e-2 |
| Cellular component assembly | 2459 | 196 | 148.55 | 1.32 | 1.18e-4 | 1.16e-2 |
| G protein-coupled receptor signaling pathway | 1248 | 39 | 75.39 | 0.52 | 5.44e-6 | 8.92e-4 |
| Adaptive immune response | 681 | 11 | 41.14 | 0.27 | 6.50e-8 | 1.60e-5 |
| Lymphocyte mediated immunity | 257 | 2 | 15.53 | 0.13 | 6.48e-5 | 7.00e-3 |
| Detection of chemical stimulus involved in sensory perception of smell | 443 | 1 | 26.76 | 0.04 | 1.63e-10 | 9.54e-8 |
| Antimicrobial humoral response | 150 | 0 | 9.06 | < 0.01 | 2.69e-4 | 2.39e-2 |

**Supplementary Table 5:** Correlation between the probability of a Neanderthal region, calculated in [50], and the probability that the same region descends from population *A* or *B*, for various populations. All correlations have a p-value less than 1e-25.

| Population | $c = AA$ | $c = BB$ |
| --- | --- | --- |
| CEU | -0.083 | 0.118 |
| GBR | -0.115 | 0.147 |
| FIN | -0.097 | 0.161 |
| IBS | -0.07 | 0.092 |
| TSI | -0.084 | 0.12 |
| JPT | -0.06 | 0.086 |
| CHS | -0.074 | 0.093 |
| CHB | -0.09 | 0.1 |

**Supplementary Table 6:** Correlations between various ancestral lineage paths and human-archaic divergence.

| Population | Archaic | $P(c = AA\vert t > T_1)$ | $P(c = BB\vert t > T_1)$ | $P(c = AA\vert T_1 \leq t < T_2)$ | $P(c = BB\vert T_1 \leq t < T_2)$ | $P(c = AA\vert t > T_2)$ | $P(c = BB\vert t > T_2)$ | $P(c = AB\vert t > T_2)$ |
| --- | --- | --- | --- | --- | --- | --- | --- | --- |
| GBR | Altai | -0.336 | 0.124 | -0.225 | 0.225 | -0.156 | 0.037 | 0.102 |
|  | Denisovan | -0.337 | 0.124 | -0.227 | 0.227 | -0.156 | 0.037 | 0.101 |
| TSI | Altai | -0.341 | 0.207 | -0.237 | 0.237 | -0.165 | 0.084 | 0.091 |
|  | Denisovan | -0.343 | 0.207 | -0.238 | 0.238 | -0.165 | 0.083 | 0.09 |
| IBS | Altai | -0.337 | 0.195 | -0.225 | 0.225 | -0.175 | 0.081 | 0.123 |
|  | Denisovan | -0.337 | 0.196 | -0.227 | 0.227 | -0.176 | 0.082 | 0.122 |
| FIN | Altai | -0.336 | 0.052 | -0.219 | 0.219 | -0.157 | 0.004 | 0.112 |
|  | Denisovan | -0.337 | 0.052 | -0.22 | 0.22 | -0.157 | 0.006 | 0.112 |
| CEU | Altai | -0.338 | 0.171 | -0.232 | 0.232 | -0.161 | 0.06 | 0.096 |
|  | Denisovan | -0.338 | 0.172 | -0.234 | 0.234 | -0.161 | 0.061 | 0.093 |
| CHS | Altai | -0.317 | -0.018 | -0.205 | 0.205 | -0.144 | -0.027 | 0.113 |
|  | Denisovan | -0.319 | -0.019 | -0.206 | 0.206 | -0.145 | -0.028 | 0.115 |
| KHV | Altai | -0.313 | -0.01 | -0.202 | 0.202 | -0.143 | -0.027 | 0.113 |
|  | Denisovan | -0.315 | -0.01 | -0.203 | 0.203 | -0.146 | -0.028 | 0.116 |
| CHB | Altai | -0.315 | -0.058 | -0.194 | 0.194 | -0.141 | -0.044 | 0.118 |
|  | Denisovan | -0.316 | -0.057 | -0.196 | 0.196 | -0.142 | -0.044 | 0.119 |
| CDX | Altai | -0.317 | -0.04 | -0.2 | 0.2 | -0.152 | -0.044 | 0.127 |
|  | Denisovan | -0.319 | -0.041 | -0.202 | 0.202 | -0.153 | -0.045 | 0.128 |
| JPT | Altai | -0.309 | -0.004 | -0.202 | 0.202 | -0.143 | -0.023 | 0.113 |
|  | Denisovan | -0.31 | -0.004 | -0.202 | 0.202 | -0.144 | -0.025 | 0.115 |
| BEB | Altai | -0.352 | 0.215 | -0.246 | 0.246 | -0.18 | 0.094 | 0.114 |
|  | Denisovan | -0.354 | 0.216 | -0.248 | 0.248 | -0.181 | 0.094 | 0.114 |
| PJL | Altai | -0.348 | 0.182 | -0.245 | 0.245 | -0.169 | 0.066 | 0.101 |
|  | Denisovan | -0.35 | 0.184 | -0.248 | 0.248 | -0.17 | 0.067 | 0.102 |
| GIH | Altai | -0.348 | 0.054 | -0.227 | 0.227 | -0.166 | 0.002 | 0.122 |
|  | Denisovan | -0.348 | 0.054 | -0.228 | 0.228 | -0.166 | 0.003 | 0.122 |
| STU | Altai | -0.344 | 0.208 | -0.241 | 0.241 | -0.171 | 0.088 | 0.094 |
|  | Denisovan | -0.347 | 0.21 | -0.245 | 0.245 | -0.174 | 0.09 | 0.097 |
| ITU | Altai | -0.344 | 0.214 | -0.245 | 0.245 | -0.174 | 0.091 | 0.101 |
|  | Denisovan | -0.345 | 0.214 | -0.245 | 0.245 | -0.175 | 0.09 | 0.102 |
| PUR | Altai | -0.369 | 0.22 | -0.252 | 0.252 | -0.208 | 0.113 | 0.168 |
|  | Denisovan | -0.368 | 0.22 | -0.253 | 0.253 | -0.208 | 0.113 | 0.167 |
| CLM | Altai | -0.352 | 0.209 | -0.24 | 0.24 | -0.189 | 0.097 | 0.143 |
|  | Denisovan | -0.354 | 0.211 | -0.242 | 0.242 | -0.192 | 0.098 | 0.145 |
| PEL | Altai | -0.31 | 0.192 | -0.219 | 0.219 | -0.15 | 0.079 | 0.079 |
|  | Denisovan | -0.311 | 0.191 | -0.219 | 0.219 | -0.15 | 0.076 | 0.078 |
| MXL | Altai | -0.331 | 0.16 | -0.221 | 0.221 | -0.179 | 0.068 | 0.139 |
|  | Denisovan | -0.331 | 0.16 | -0.223 | 0.223 | -0.179 | 0.068 | 0.139 |
| ESN | Altai | -0.408 | 0.268 | -0.307 | 0.307 | -0.267 | 0.163 | 0.219 |
|  | Denisovan | -0.409 | 0.267 | -0.308 | 0.308 | -0.268 | 0.161 | 0.22 |
| YRI | Altai | -0.41 | 0.271 | -0.311 | 0.311 | -0.259 | 0.158 | 0.182 |
|  | Denisovan | -0.41 | 0.271 | -0.311 | 0.311 | -0.259 | 0.156 | 0.183 |
| MSL | Altai | -0.409 | 0.273 | -0.31 | 0.31 | -0.272 | 0.171 | 0.229 |
|  | Denisovan | -0.409 | 0.271 | -0.31 | 0.31 | -0.271 | 0.169 | 0.229 |
| GWD | Altai | -0.408 | 0.276 | -0.309 | 0.309 | -0.256 | 0.163 | 0.179 |
|  | Denisovan | -0.406 | 0.275 | -0.309 | 0.309 | -0.255 | 0.161 | 0.179 |
| ACB | Altai | -0.42 | 0.28 | -0.32 | 0.32 | -0.274 | 0.17 | 0.207 |
|  | Denisovan | -0.42 | 0.281 | -0.322 | 0.322 | -0.275 | 0.169 | 0.209 |
| ASW | Altai | -0.418 | 0.268 | -0.307 | 0.307 | -0.262 | 0.156 | 0.196 |
|  | Denisovan | -0.42 | 0.269 | -0.31 | 0.31 | -0.262 | 0.158 | 0.195 |
| LWK | Altai | -0.414 | 0.273 | -0.314 | 0.314 | -0.261 | 0.162 | 0.186 |
|  | Denisovan | -0.414 | 0.271 | -0.314 | 0.314 | -0.262 | 0.159 | 0.189 |

**Supplementary Table 7:** Triplet codes used for each of the 26 populations in the 1000 Genomes Project.

| ID | Population |
| --- | --- |
| ASW | African Ancestry in Southwestern USA |
| ACB | African Caribbeanin Barbados |
| BEB | Bengali in Bangladesh |
| GBR | British from England and Scotland |
| CDX | Chinese Dai in Xishuangbanna, China |
| CLM | Colombian in Medellin, Colombia |
| ESN | Esan in Nigeria |
| FIN | Finnish in Finland |
| GWD | Gambian in Western Division - Mandinka |
| GIH | Gujarati Indians in Houston, Texas, United States |
| CHB | Han Chinese in Beijing, China |
| CHS | Han Chinese South, China |
| IBS | Iberian populations in Spain |
| ITU | Indian Telugu in the UK |
| JPT | Japanese in Tokyo, Japan |
| KHV | Kinh in Ho Chi Minh City, Vietnam |
| LWK | Kenya Luhya in Webuye, Kenya |
| MSL | Sierra Leone Mende in Sierra Leone |
| MXL | Mexican Ancestry in Los Angeles, California, United States |
| PEL | Peruvian in Lima, Peru |
| PUR | Puerto Rico Puerto Rican in Puerto Rico |
| PJL | Punjabi in Lahore, Pakistan |
| STU | Sri Lankan Tamil in the UK |
| TSI | Toscani in Italy |
| YRI | Yoruba in Ibadan, Nigeria |
| CEU | Utah residents with Northern and Western European ancestry from the CEPH collection |

**Supplementary Table 8:** Using the inferred parameters on the seen data (training), we calculate the loglikelihood of unseen data (testing) for a structured and unstructured model. The strongly positive Δ_ℒ_ = ℒ_*S*_ − ℒ_*U*_ in the test data indicates the better fitting structured model is not due to overfitting.

| Population | Training |  | Testing |  |
| --- | --- | --- | --- | --- |
| | Sample | $\Delta_{\mathcal{L}}$ | Sample | $\Delta_{\mathcal{L}}$ |
| GBR | HG00118 | 1153.3 | HG00119 | 1140.1 |
| TSI | NA20752 | 1277.0 | NA20753 | 1101.0 |
| IBS | HG01783 | 994.9 | HG02221 | 882.7 |
| FIN | HG00266 | 1120.5 | HG00267 | 963.9 |
| CEU | NA12718 | 1125.6 | NA12748 | 1133.9 |
| CHS | HG00443 | 1331.8 | HG00445 | 1412.0 |
| KHV | HG02113 | 1355.6 | HG02141 | 1368.4 |
| CHB | NA18530 | 1535.8 | NA18534 | 1320.4 |
| CDX | HG02373 | 1485.8 | HG02394 | 1494.4 |
| JPT | NA18939 | 1425.2 | NA18941 | 1295.6 |
| BEB | HG03006 | 1058.4 | HG03007 | 1276.4 |
| PJL | HG03234 | 1116.8 | HG03235 | 1083.2 |
| GIH | NA20845 | 1095.4 | NA20847 | 1124.5 |
| STU | HG03753 | 1202.3 | HG03760 | 1132.7 |
| ITU | HG03977 | 1319.8 | HG03780 | 1263.1 |
| PUR | HG01171 | 769.8 | HG01173 | 872.5 |
| CLM | HG01250 | 866.2 | HG01251 | 994.4 |
| PEL | HG02285 | 1168.4 | HG02286 | 1315.7 |
| MXL | NA19648 | 794.1 | NA19649 | 908.3 |
| ESN | HG03515 | 499.5 | HG03514 | 647.0 |
| YRI | NA18488 | 778.9 | NA19222 | 757.4 |
| MSL | HG03212 | 449.7 | HG03055 | 694.6 |
| GWD | HG02568 | 648.5 | HG02759 | 391.9 |
| ACB | HG01882 | 501.0 | HG01883 | 526.8 |
| ASW | NA19625 | 601.9 | NA19700 | 600.7 |
| LWK | NA19017 | 598.4 | NA19019 | 609.6 |

## 9 Supplementary Text

### 9.1 Previous work

The sequentially Markovian coalescent (SMC) [30] was introduced to describe analytically the probability of local changes in ancestry. Moving left to right across a genome sequence, it describes the probability of the next tree height (i.e. coalescence time) as conditionally dependent only on the current tree height, as opposed to the whole genealogy of the sequence thus far [97], enabling tractable inference. Marjoram and Wall proposed a modification to the SMC model [31], which more accurately models the coalescent with recombination [98] (previously, authors have used SMC’ to denote the modification proposed by Marjoram and Wall, though here and in the main text we use SMC to mean the modified version). By assuming panmixia and using a discretised version of the conditional distribution from SMC in the transition matrix of its hidden Markov model (HMM), the pairwise sequentially Markovian coalescent (PSMC) [19] infers the changes in the effective population size over time and the coalescence times across the genome.

Wang et al. [39] introduced MSMC-IM to estimate a time-dependent rate of gene flow between two sampled populations. Given sequences from a pair of populations, it uses MSMC2 to infer coalescence rates within and across these populations, then fits a continuous Isolation-Migration model to these. It assumes that the migration rate was symmetric, and does not attempt to interpret coalescence rates in terms of population size. Simulations from Shchur et al. [32] indicated that PSMC’s transition matrix has information that may be able to distinguish structure from panmixia. They developed MiSTI, which uses the joint SFS from two diploid sequences to estimate time-dependent size changes and continuous, non-symmetric migration rates.

### 9.2 PSMC’s HMM and notation

In this section we review the model underlying PSMC, and introduce the notation that we use throughout. The original PSMC [19] uses Cardin and McVean’s SMC model [30], which is sufficient for accurate parameter inference, but MSMC, MSMC2, and PSMC+ [99, 39, 70] use Marjoram and Wall’s SMC model [31], which is a more accurate model of the coalescent with recombination [98]. For simplicity, henceforth we use “PSMC” to refer to a non-implementation specific version of the PSMC using the Marjoram and Wall’s SMC framework.

We measure time, *t*, in coalescent units going backwards in time (i.e. *t* = 0 is the present). Taking two lineages from population *A* at *t* = 0, PSMC infers the coalescence rate *λ*_*A*_(*t*) and uses

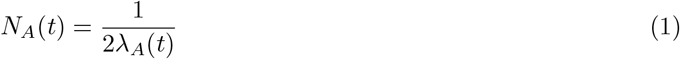

to estimate the changes through time in the effective size of population *A*.

#### 9.2.1 Emissions and transitions

PSMC is a HMM where the observations are the sequence of homozygotes or heterozygotes across the genome, which we label *X* = (*x*_1_, *x*_2_, …, *x*_*L*_) where *L* is the length of the sequence and *x* = 0 for a homozygote, *x* = 1 for a heterozygote and *x* = −1 for missing data. The latent variable at each position is the coalescence time, or time to most recent common ancestor (TMRCA), which we denote **Z** = (*z*_1_, *z*_2_, …, *z*_*L*_), where theoretically *z*_*i*_ takes values [0, ∞). We make the infinite sites assumption that at most one mutation can occur at any site, therefore at a single base pair level the emission probabilities are:

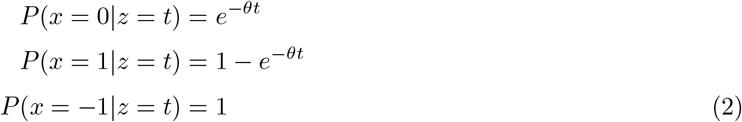

where *θ* = 4*N*_*A*_*µ* is the scaled mutation rate, *N*_*A*_ is the diploid, long-term effective population size, *µ* is the per base per generation mutation rate, and *t* is measured in coalescent units. By chopping the genome sequence into bins of *b* base pairs, we can achieve a linear computational speed-up by a factor of *b*. In each bin there are then *h* heterozygotes, *m* missing bases and *b* − *m* − *h* homozygotes, with 0 ≤ *m, h* ≤ *b*. The number of heterozygotes in a bin can then be modelled as a Poisson, and so the emission probabilities are thus:

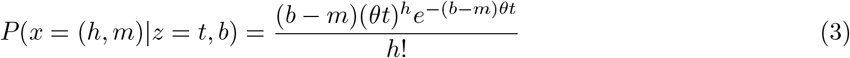

Denote the *i*’th locus as having TMRCA *s* and the (*i* + 1)’th locus having tMRCA *t*. Let *R* and *R*^*′*^ denote the instance or lack of a recombination, respectively, between loci *i* and *i* + 1. At the base pair level, the probability of recombining or not is *P* (*R*|*z*_*i*_ = *t*) = *ρ* and *P* (*R*^*′*^|*z*_*i*_ = *t*) = 1 − *ρ*, where 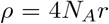 is the scaled recombination rate, *r* is recombination rate per neighbouring base pairs per generation, and under SMC’ recombination can occur at most once between neighbouring loci. With the sequence in bins of size *b*, the probability of not recombining between neighbouring bins is *P* (*R*^*′*^|*z*_*i*_ = *t, b*) = (1 − *ρ*)^*b*^ so the probability of at least one recombination occurring is *P* (*R*|*z*_*i*_ = *t, b*) = 1 − (1 − *ρ*)^*b*^ ≈ 1 − *e*^*−bρ*^. In what follows, for simplicity we typically do not write the *b* term explicitly.

The probability of transitioning between *z*_*i*_ = *s* and *z*_*i*+1_ = *t* is governed by the SMC model [31]. For brevity let *P* (*t*|*s*) = *P* (*z*_*i*+1_ = *t*|*z*_*i*_ = *s*) denote the probability of transitioning under the SMC. Considering the presence or absence of recombination, we can write this as

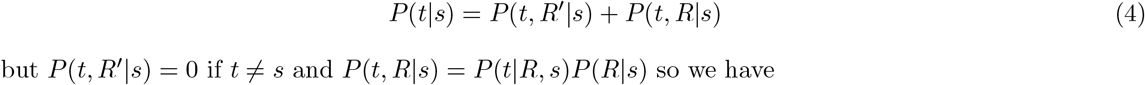

but *P* (*t, R*^*′*^|*s*) = 0 if *t*≠*s* and *P* (*t, R*|*s*) = *P* (*t*|*R, s*)*P* (*R*|*s*) so we have

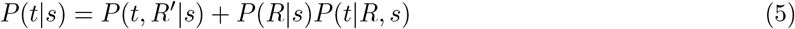

Using the SMC model for two samples, given that a recombination has occurred there are three distinct types of coalescent event:

**Supplementary Figure 21:**
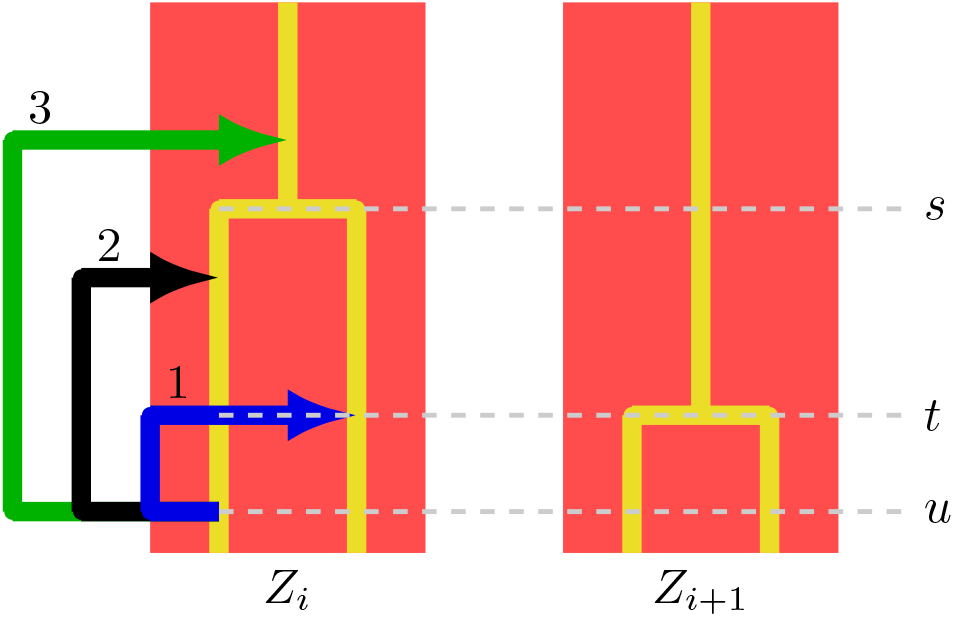
A depiction of the SMC framework for locus *i* and *i* + 1. The tree height (i.e. TMRCA) at locus *i* is *s* and at *i* + 1 is *t*. Under the SMC model, given that a recombination has occurred at time *u* there are three possible types of coalescence event. The coloured arrows depict the “floating” lineage, which can either coalesce to the branch on which recombination did not occur (1), the branch on which recombination did occur (2), or the ancestral branch (3). In the diagram shown, a type 1 event is shown.

1. The floating^1^ lineage coalesces to the other solid lineage at time *t < s*
2. The floating lineage coalesces back to the same branch on which the recombination occurred, resulting in *t* = *s*
3. The floating lineage coalesces to the solid ancestral lineage at time *t > s*

A depiction of this process is given in Figure S21. We note that Cardin and McVean’s SMC model only allows events 2 or 3.

Let *u* denote the time at which a recombination occurred (then strictly *u* ≤ *t* and *u < s*). If a type 1 event occurs, we must have no coalescence with either solid lineages between *u* and *t*, then a coalescence at time *t* to the branch that did not recombine. For a type 2 event, we must have the floating lineage coalescence with the lineage on which recombinaton occurred, at any time between *u* and *s*. For a type 3 event, we must have no coalescence between *u* and *t* on either solid branches, and subsequently no coalescence between *s* and *t* on the single ancestral solid lineage. Define

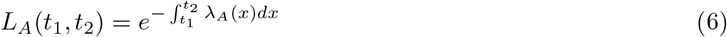

then we can write the probability of transitioning under the SMC’ as:

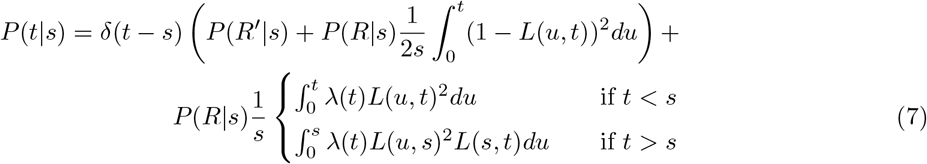

Where we integrate over *u* because it is a variable with values in [0, *min*(*s, t*)], the 1*/s* terms are a normalisation constant for the integration over *u*, and *δ* is the Kronecker delta which handles presence/absence of recombination. Finally, the stationary distribution is defined as

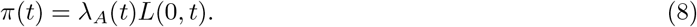

See [39] for the full details.

#### 9.2.2 Discretisation

Primarily the goal of PSMC is to infer changes in the effective population size over time from some population *A*. To do this, PSMC discretises the space of TMRCAs into *D* time intervals by partitioning [0, ∞) into a state space of *D* time intervals with boundaries ***τ*** = (*τ*_0_, …, *τ*_*D*_) which for *i* ∈ (1, …, *D* − 1) are defined by

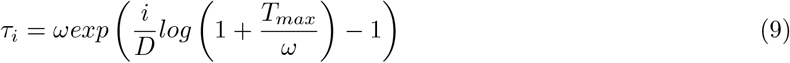

where *ω* and *T*_*max*_ control their spread, and *τ*_0_ = 0 and *τ*_*D*_ = ∞. The set of hidden states in the HMM is **T** = {*T*_1_, …, *T*_*D*_} and if the TMRCA at locus *i* is *t* with *τ*_*α*_ ≤ *t < τ*_*α*+1_ then *z*_*i*_ = *T*_*α*_. Denote the set of *D* inverse effective population size parameters with ***λ***_*A*_ = (*λ*_*A*_1, …, *λ*_*A*_*D*). Suppose that *t* is contained within interval *α*, and *s* in interval *β*, i.e. *T*_*α*_ ≤ *t < T*_*α*+1_ and *T*_*β*_ ≤ *s < T*_*β*+1_. To get the discrete emission probabilities, in MSMC2 [99] Schiffels et al. integrate between the neighbouring boundaries combined with the stationary distribution *π*(*t*), though we simply take the midpoint:

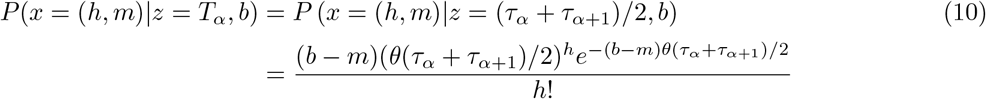

Similarly, to create the transition matrix *Q*(*α*|*β*) = *P* (*T*_*α*_|*T*_*β*_), Schiffels et al. take the expected coalescence time in interval *β*:

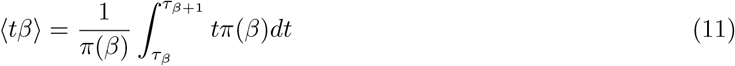

and integrate out *α*:

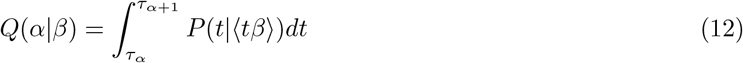

#### 9.2.3 Parameter inference

To infer the demographic parameters, PSMC uses an expectation-maximisation (EM) algorithm. Defining *f*_*β*_(*i*) = *P* (*x*_1:*i*_, *z*_*i*_ = *T*_*β*_) and *b*_*α*_(*i*) = *P* (*x*_*i*+1:*L*_|*z*_*i*_ = *T*_*α*_) as the forward and backward quantities respectively, we can calculate the expected transition matrix

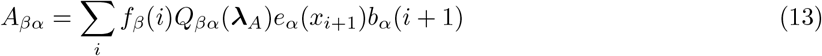

where *Q*^*βα*^(***λ***_*A*_) denotes *Q*(*α*|*β*, ***λ***_*A*_), and update the demographic parameters from iteration *v* by maximising

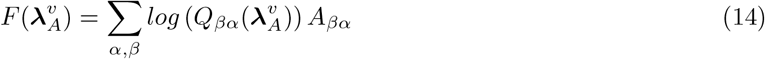

and setting

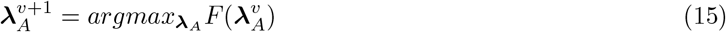

### 9.3 *cobraa* ‘s HMM

The hidden states and emissions of *cobraa* are exactly the same as in PSMC. The emissions of *cobraa* are therefore given in equation (10). The transition probabilities however are different, because *cobraa* models changes in local ancestry according to a structured model, where as PSMC models according to an unstructured model. In the next section we thus derive the transition probabilities as a function of the structured model, by allowing migration in the SMC framework.

#### 9.3.1 Transition probabilities under the structured model

In this section, we use the same notation for the SMC model as in 9.2.1: *s* is the given coalescence time for locus *i*, with next lowest time interval boundary *τ*_*β*_; *t* is the new coalescence time for locus *i* + 1 which is only dependent on locus *i*, with next lowest time interval boundary *τ*_*α*_; *u* is the time of recombination which is strictly less than the minimum of (*s, t*) and greater than zero.

Under the SMC model, lineages can only either recombine or coalesce, resulting in a singular constraint that coalescence time must be greater than recombination time. If we introduce population structure, then lineages are also able to migrate. This induces some mutual exclusiveness, because certain types of coalescence event are dissalowed. For example, in the *pulse* structure model, if we have a recombination at time *u < T*_1_ where both solid lineages stay in population *A* but the floating lineage migrates to population *B*, then we cannot have any coalescence event at *t* ∈ (*T*_1_, *T*_2_] because there are no solid lineages for the floating lineage to coalesce with. We therefore partition the structured transition matrix into 10 distinct Cases:

1. *t* < *T*_1_ and *s>t* with s*∈* (*t,∞*)
2. *T*_1_ ≤ *t* < *T*_2_ and *T*_1_ ≤ *s* < *T*_2_ with *s > t*.
3. *T*_1_ ≤ *t* < *T*_2_ and *T*_2_ ≤ *s*
4. *T*_2_ ≤ *t* and *T*_2_ ≤ *s* with *s > t*.
5. *t* < *T*_1_ and *s< T*_1_ with t*∈s*
6. *s* < *T*_1_ and *T*_1_ ≤ *t* < *T*_2_.
7. *T*_1_ ≤ t < *T*_2_ and *T*_1_ ≤ *s* < *T*_2_with *t* > *s*.
8. *s* < *T*_1_, *t* > *T*_2_
9. *T*_1_ ≤ *s* < *T*_1_ and *t* > *T*_1_
10. *T*_2_ ≤ *t* and *T*_2_ ≤ *s* with *t > s*.

**Supplementary Figure 22:**
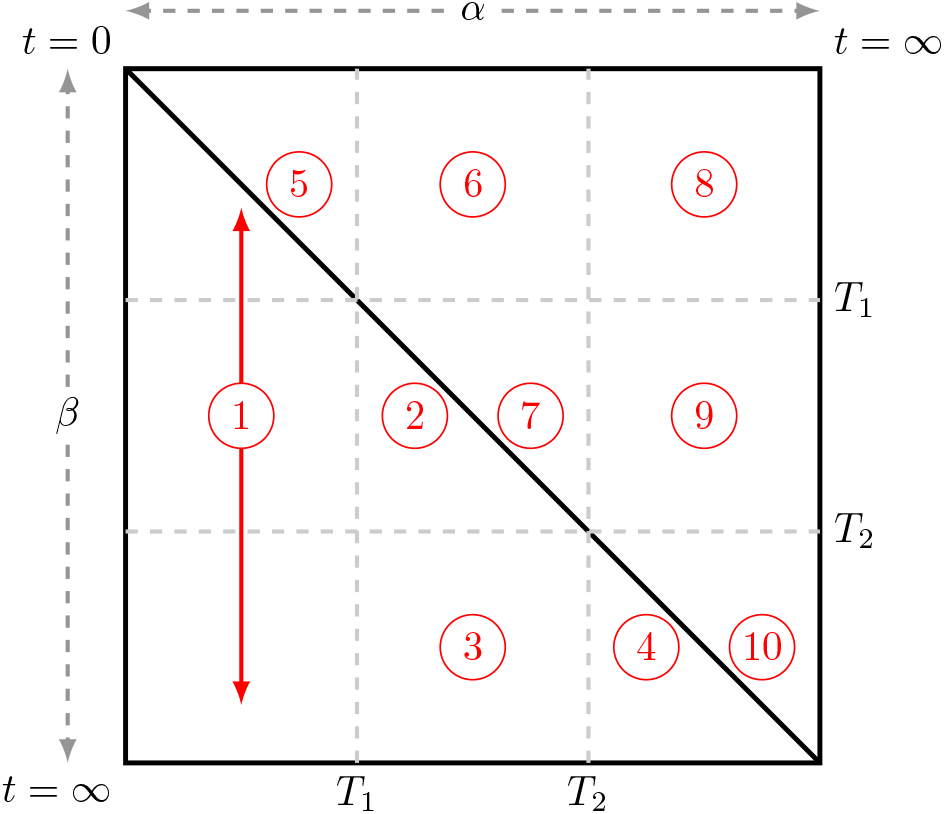
Under the SMC model with a *pulse* model of population structure, given that a recombination has occurred, we partition the transition matrix in to 10 different possible coalescence events.

An illustration of these cases is shown in Figure S22. Note that Case 1 and Case 5 are the same as the panmictic case with *s > t* and *t < s*, respectively. A possible configuration from Case 3 is shown in Figure S23, where at locus *i* the two solid lineages went through separate populations; a lineage recombines at time *u < T*_1_ and migrates to population *B*, where it coalesces with the other lineage at time *t*. This results in the new genealogy at locus *i* + 1. In what follows let *P*_*X*_ (*t*|*s*) = *P* (*z*_*i*+1_ = *t*|*z*_*i*_ = *s*, Case X) denote the probability of transitioning from TMRCA *s* to TMRCA *t* under the *pulse* migration model where *s* and *t* are bounded by the description given for Case *X*. After discretisation we write the transition matrix as 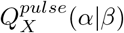 to denote 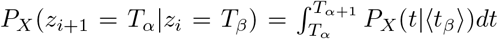 where ⟨*t*_β_ ⟩ is given by equation (11). The transition matrix under the *pulse* model *Q*^*pulse*^ is a function of ***λ***_*A*_, ***λ***_*B*_, *γ, T*_2_, and *T*_1_, which are, respectively, the discretised form of 1*/N*_*A*_(*t*) and 1*/N*_*B*_(*t*), admixture fraction, and the split and rejoin time, though we frequently omit these for brevity.

We now walk through the derivation of Case 3. The other cases are given in the following sections. Two lineages combine at *s > T*_2_ (the given current genealogy), so we must consider whether they both went through *A*, both through *B*, or one through *A* and one through *B*. Let 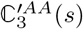 denote two uncoalesced lineages at time *s* with both lineages having passed through population *A*, under the *pulse* migration model where *s* is bounded by the description for Case 3. Then

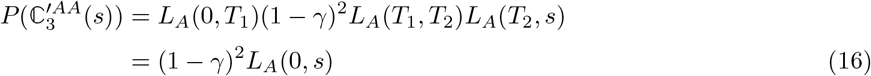

Where the *L*_*A*_ terms are the probabilities of not coalescing (equation (6)) and the (1 − *γ*)^2^ term is the

**Supplementary Figure 23:**
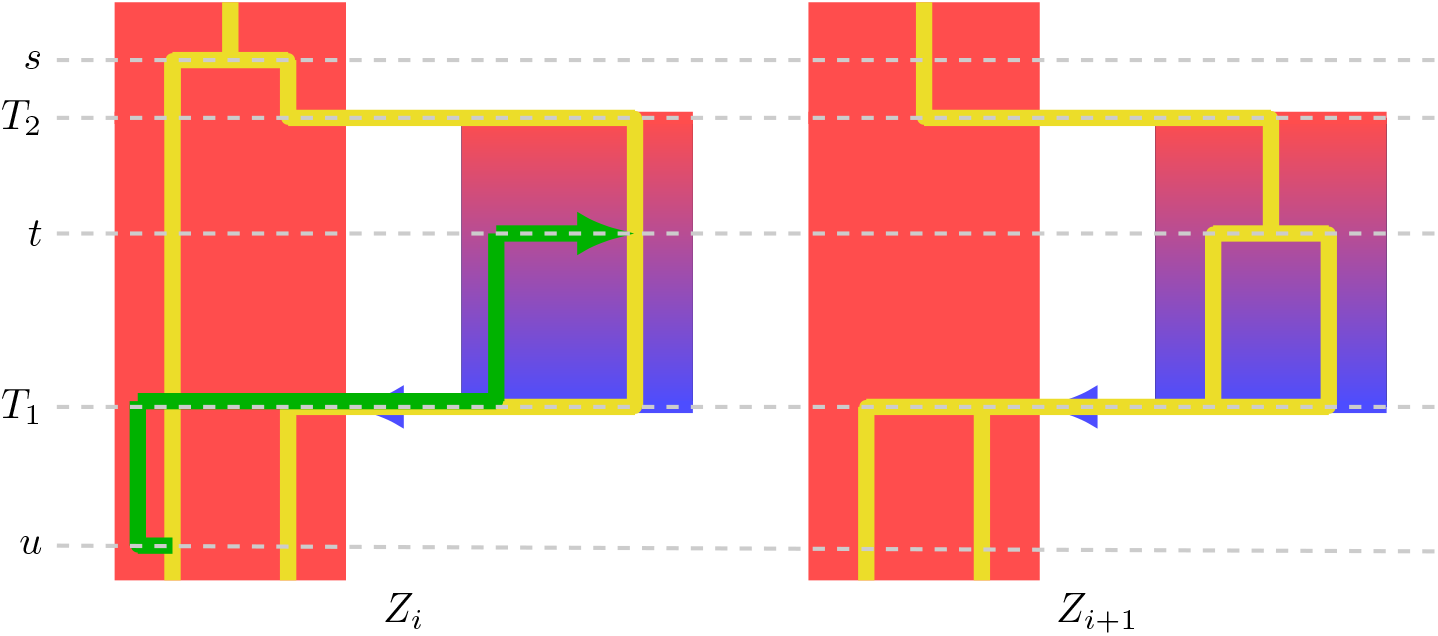
Diagram of the SMC model under *pulse* population structure. Given the genealogy at locus *i*, a recombination occurs at time *u* and the floating lineage (green) migrates to population *B* at time *T*_1_. The floating lineage then coalesces to the other solid lineage from which it recombined at time *t*, resulting in the new genealogy at locus *i* + 1.

probability that both lineages stay in population *A* at time *T*_1_. Similarly, we can write

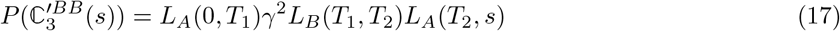

and

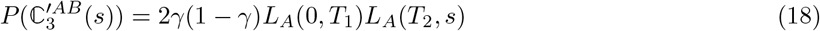

where 2*γ*(1 − *γ*) is the probability that one lineage goes through *A* and the other through *B* at time *T*_1_, and the *L*_*A*_(*T*_1_, *T*_2_) or *L*_*B*_(*T*_1_, *T*_2_) term vanishes because two lineages can not coalesce if they are in different populations. Using 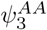 to denote the probability of both lineages passing through population *A*, then by marginalising over 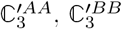 and 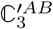 we obtain

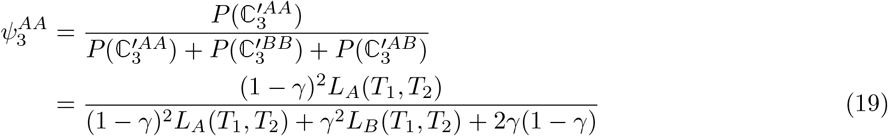

and similiarly for both lineages passing through *B*

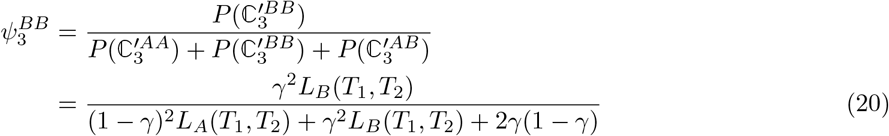

and for one lineage through *A* and one through *B*

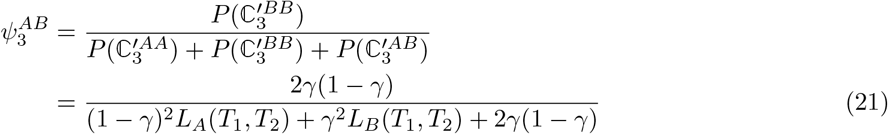

Now we consider the position of recombination, *u*, where strictly 0 ≤ *u* ≤ *min*(*s, t*). If *u* ≤ *T*_1_ then we must account for the probability of the floating lineage migrating to *B* or staying in *A*, and if there is one lineage in *A* and one in *B* then recombination can not occur after *T*_1_ because we are conditioning on a coalescent event in (*T*_1_, *T*_2_] and *t*≠*s*. Therefore, we have

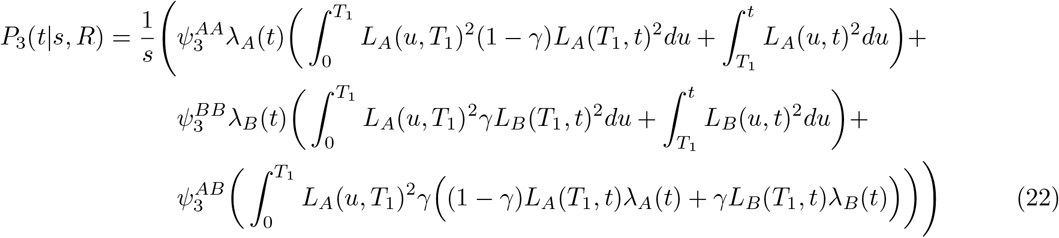

Recalling the piecewise constant assumption, we integrate out *u* by noticing that

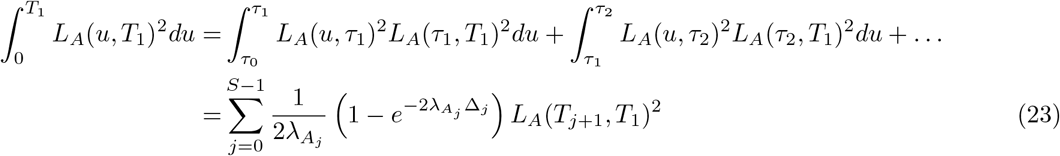

where *S* ∈ {1, *D* − 1} is the time index for *T*_1_ and Δ_*j*_ = *τ*_*j*+1_ − *τ*_*j*_ is the difference between neighbouring time interval boundaries. We generalise this expression and define

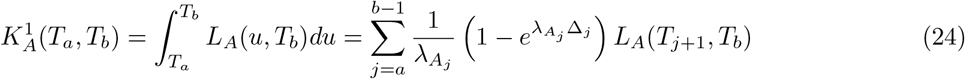

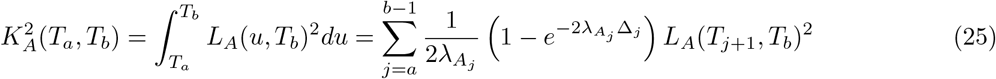

so for example

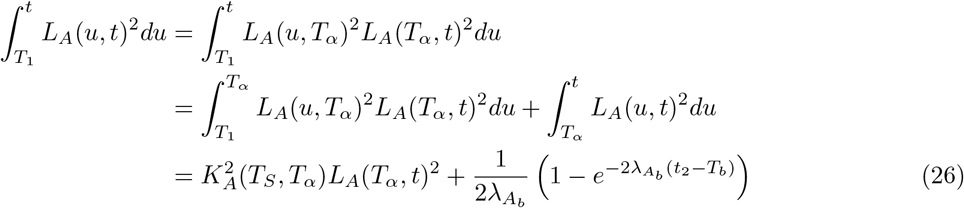

Then, we can write

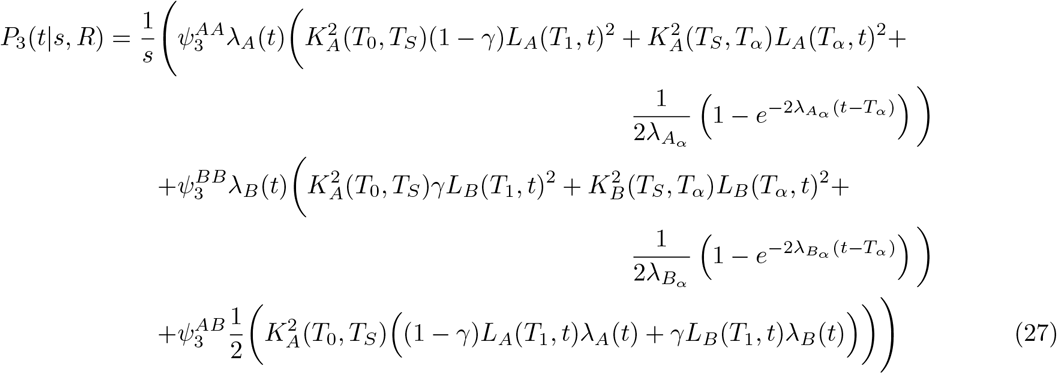

To get the transition probabilities, we must now integrate out *s* and *t* from *P*_3_(*t*|*s, R*), and take the discretised inverse population size parameters ***λ***_*A*_ and ***λ***_*B*_ . Similarly to Schiffels et al. in [99, 38], we use the expected coalescence time ⟨*t*_*β*_⟩ in interval *β* to integrate out *s* (see equation (11).

Define

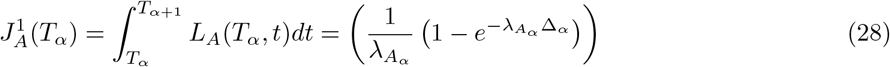

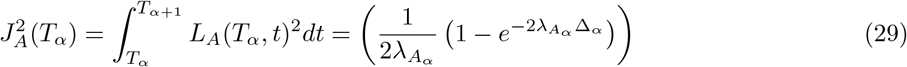

and

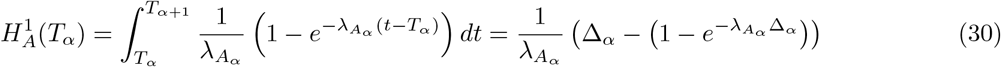

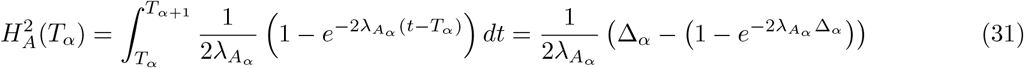

Then we have

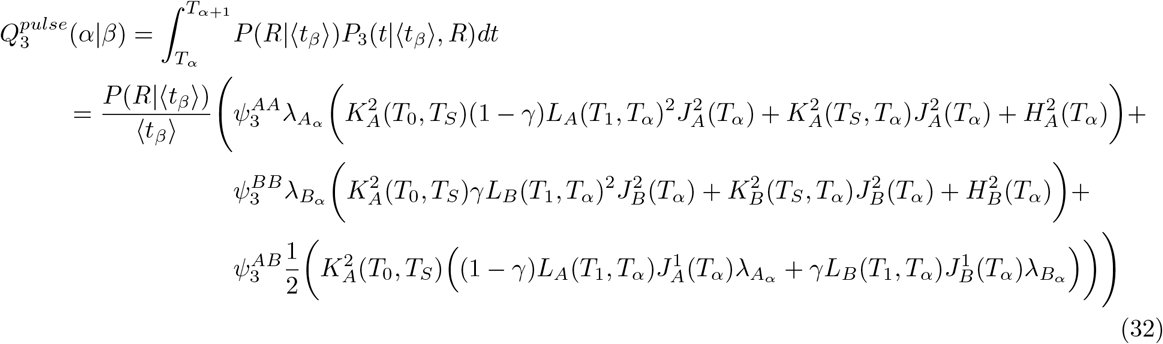

##### 9.3.1.1 Case 1

*t < T*_1_ and *s > t* with *s* ∈ (*t*, ∞)

Case 1 is the same as panmixia with *s > t*, because *t < T*_1_ so *u < T*_1_ hence we do not need to consider the structured parameters. So

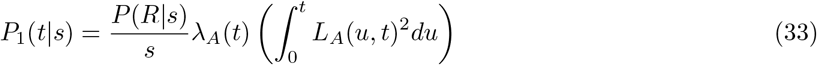

then

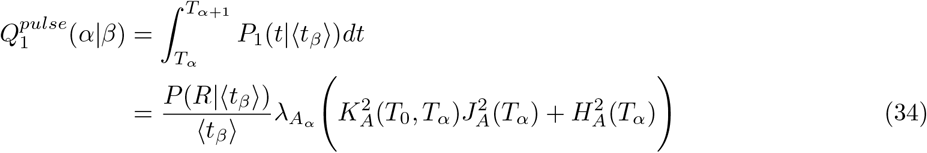

##### 9.3.1.2 Case 2

*T*_1_ ≤ *t* < *T*_2_ and *T*_1_ ≤ *s* < *T*_2_ with *s > t*.

Case 2 is very similar to Case 3, though because we are conditioning on *s, t* ∈ (*T*_1_, *T*_2_] we cannot have the solid lineages in separate populations therefore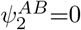 So

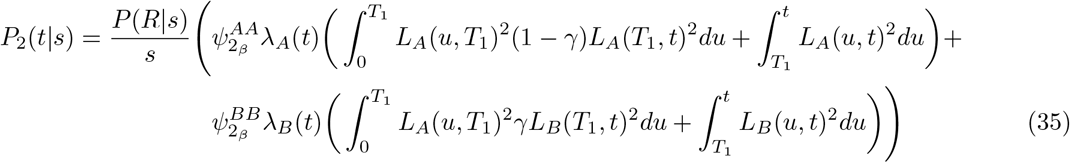

where

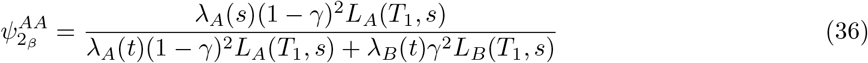

and

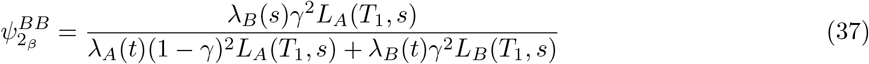

then integrating out *u, t*, and *s* yields

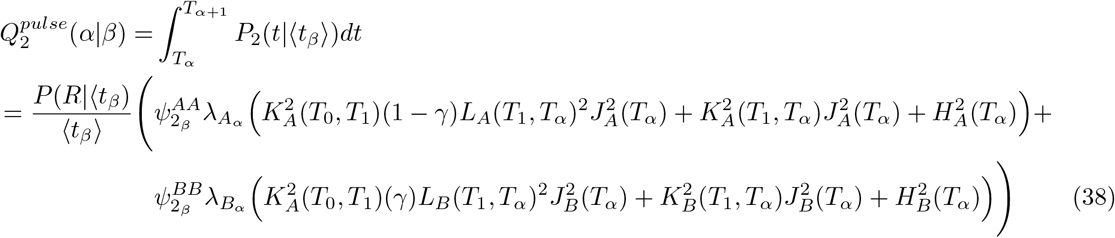

##### 9.3.1.3 Case 3

*T*_1_ ≤ *t < T*_2_ and *T*_2_ ≤ *s*

The derivation for Case 3 is given above, but for continuity we restate here

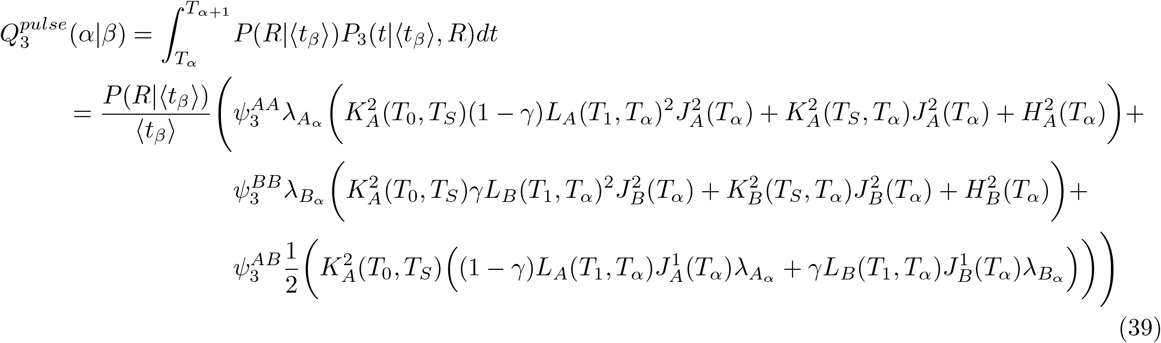

##### 9.3.1.4 Case 4

*T*_2_ ≤ *t* and *T*_2_ ≤ *s* with *s > t*

We have

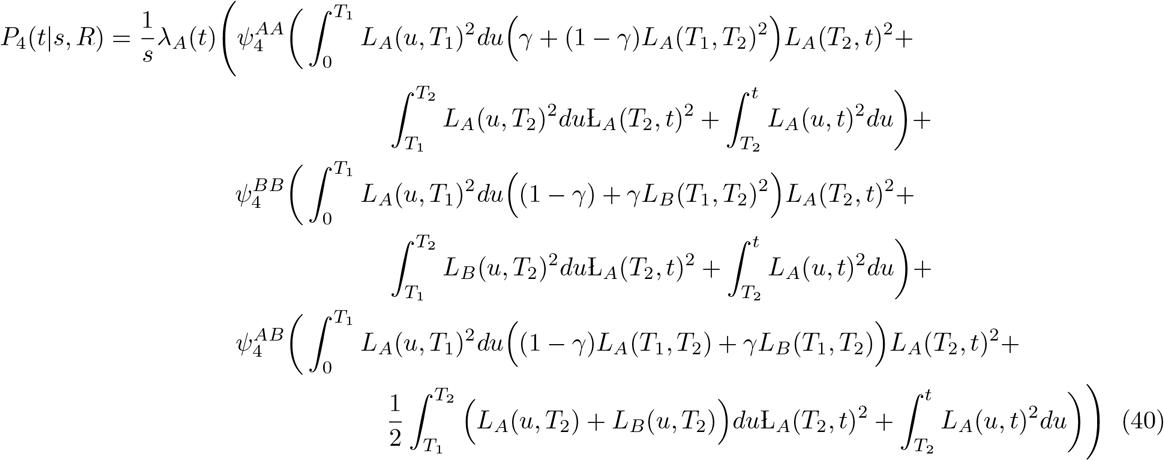

where

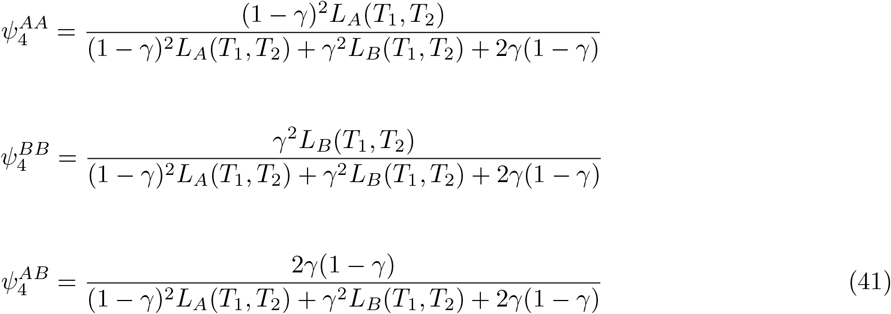

From which we obtain

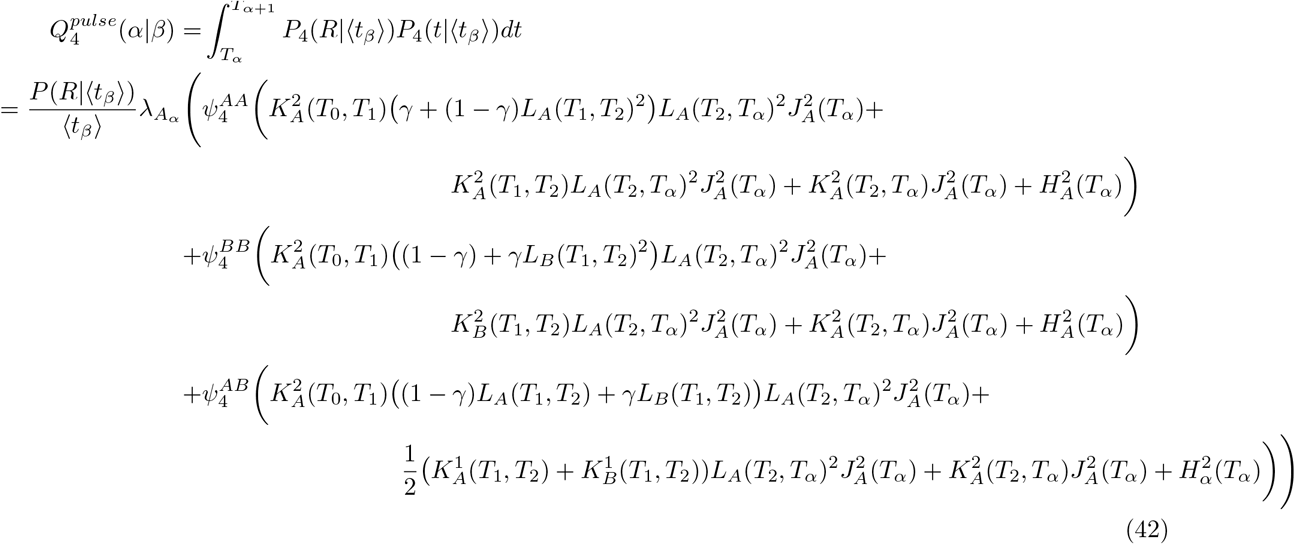

##### 9.3.1.5 Case 5

*t < T*_1_ and *s < T*_1_ with *t > s*

Case 5 is the same as panmixia with *t > s*, because *t < T*_1_ so *u < T*_1_ hence we do not need to consider the structured parameters. So

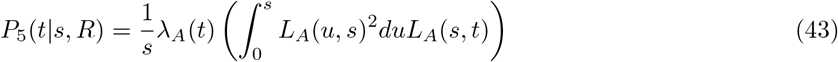

then

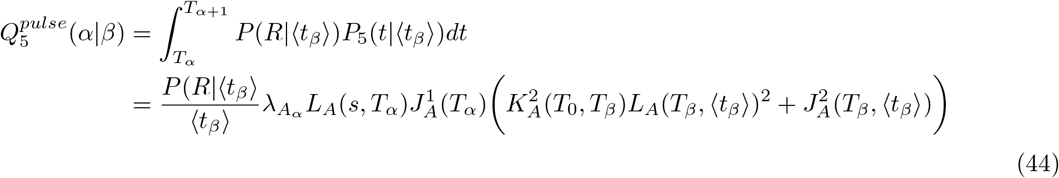

##### 9.3.1.6 Case 6

*s < T*_1_ and *T*_1_ ≤ *t < T*_2_

For Case 6, two lineages have coalesced at *s < T*_1_ so we only consider the possibility of one lineage migrating at time *T*_1_, then 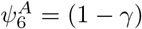 and 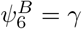 So

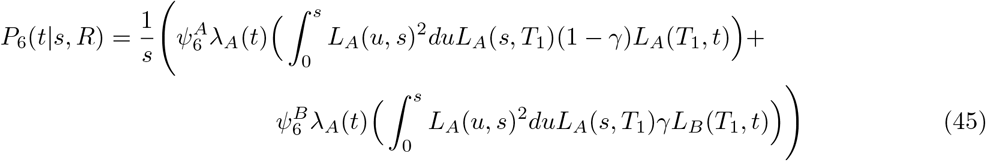

then

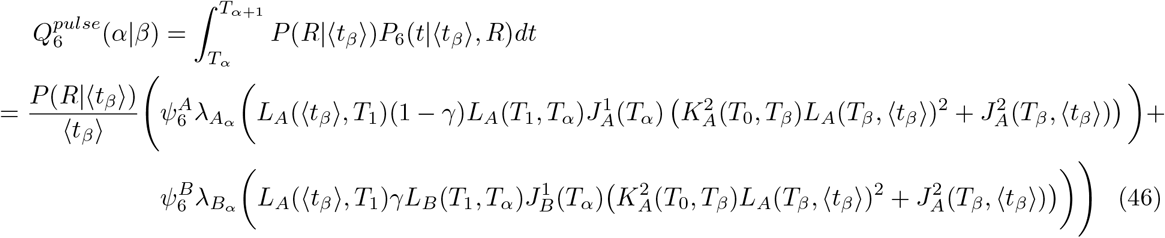

##### 9.3.1.7 Case 7

*T*_1_ ≤ *t < T*_2_ and *T*_1_ ≤ *s < T*_2_ with *t > s*

For Case 7 we have

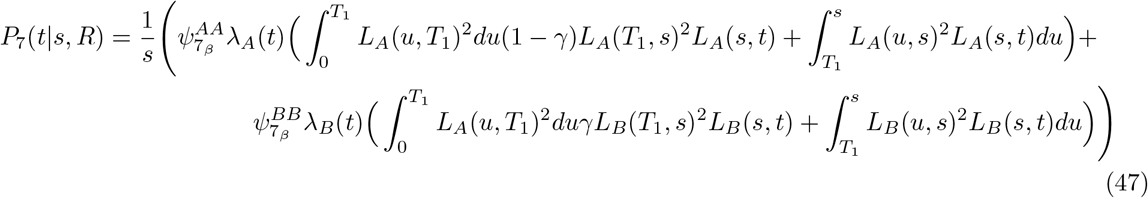

where

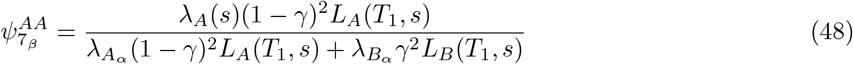

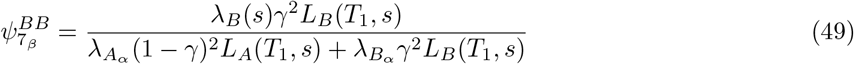

So

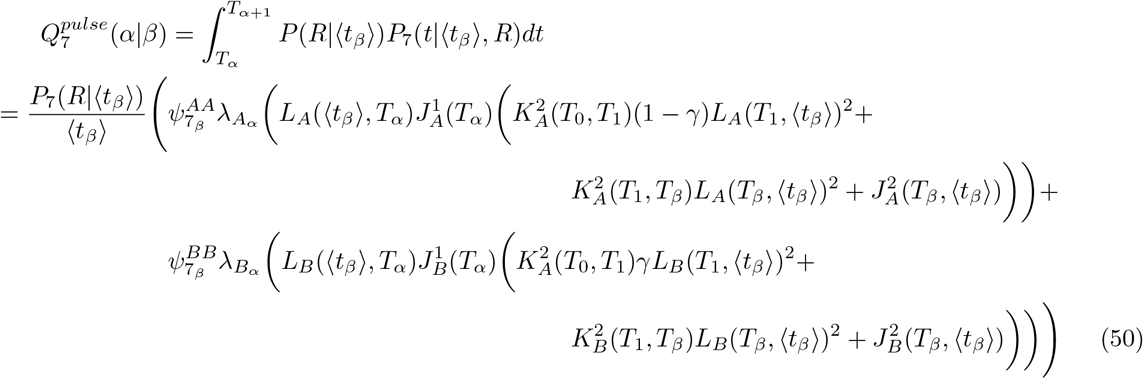

##### 9.3.1.8 Case 8

*s < T*_1_ and *t > T*_2_

For Case 8 we have where 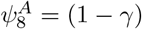 and 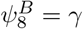

Then

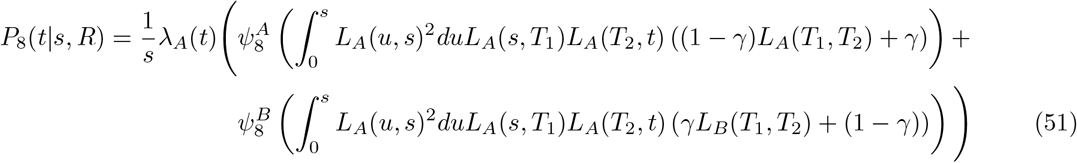

##### 9.3.1.9 Case 9

*T*_1_ ≤ *s < T*_2_ and *t > T*_2_

For Case 9 we have

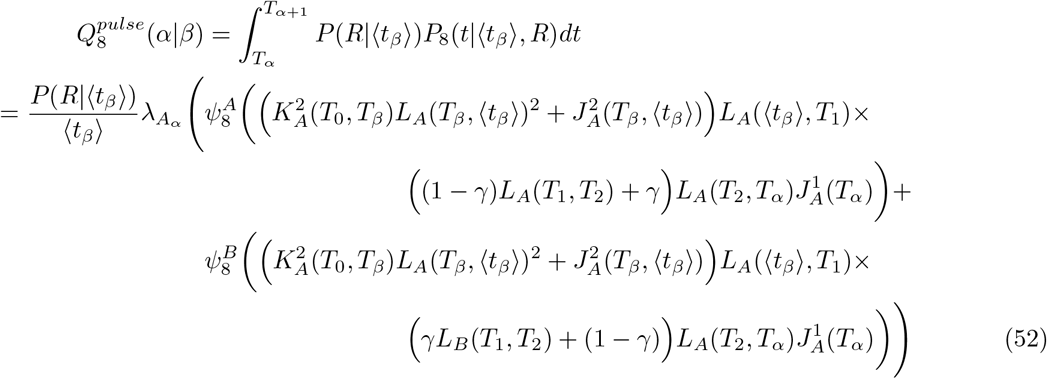

where

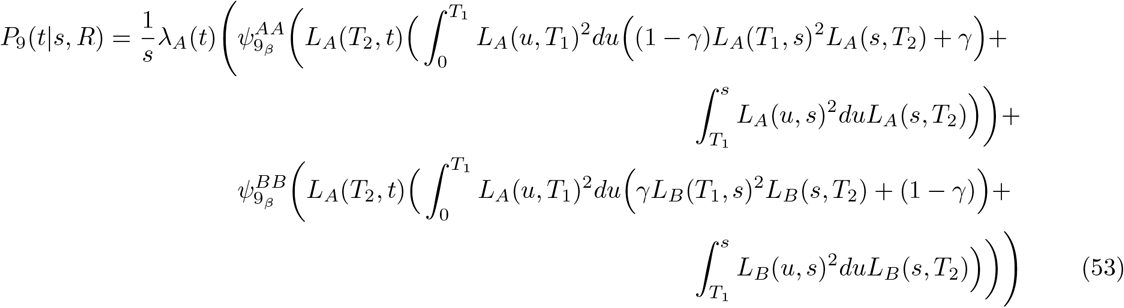

and

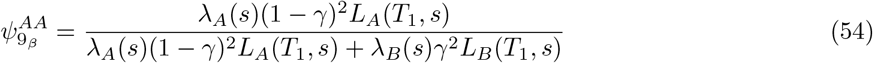

So

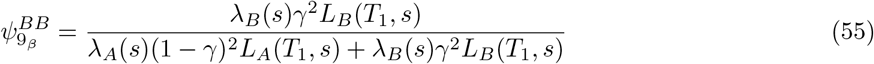

##### 9.3.1.10 Case 10

*T*_2_ ≤ *t* and *T*_2_ ≤ *s* with *t > s*

For Case 10 we have

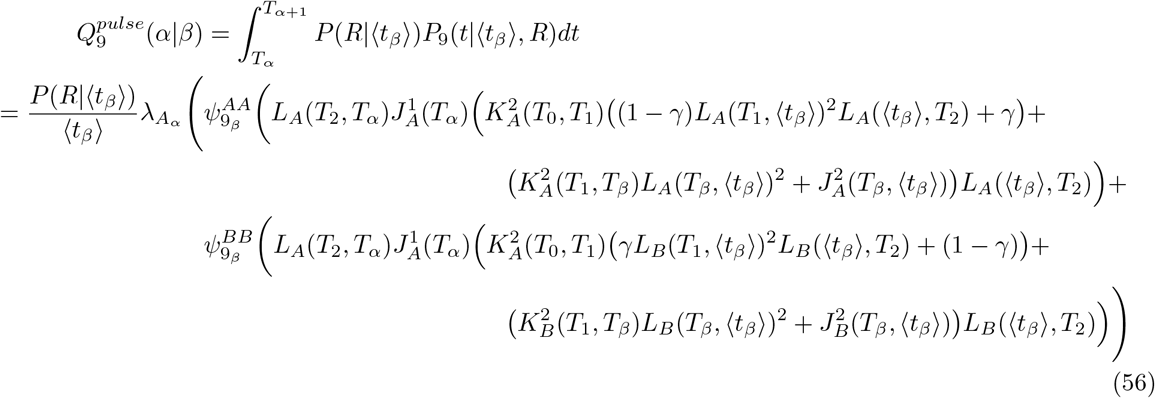

where

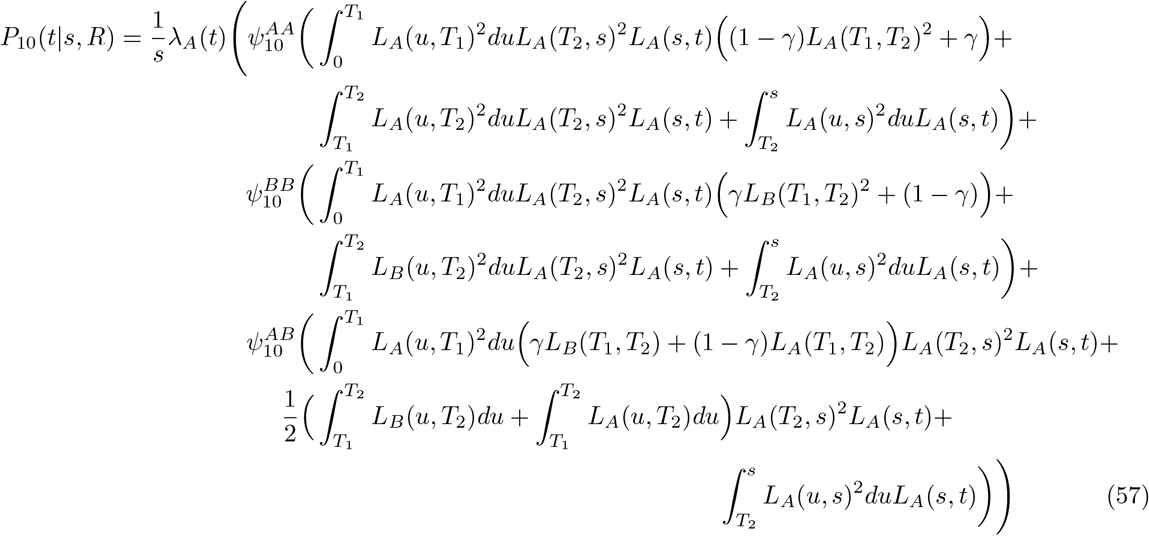

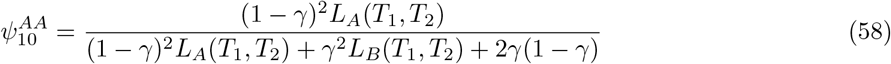

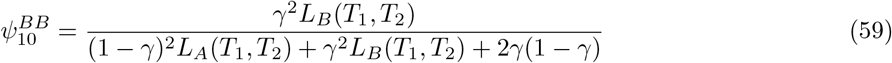

so

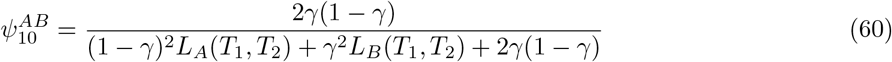

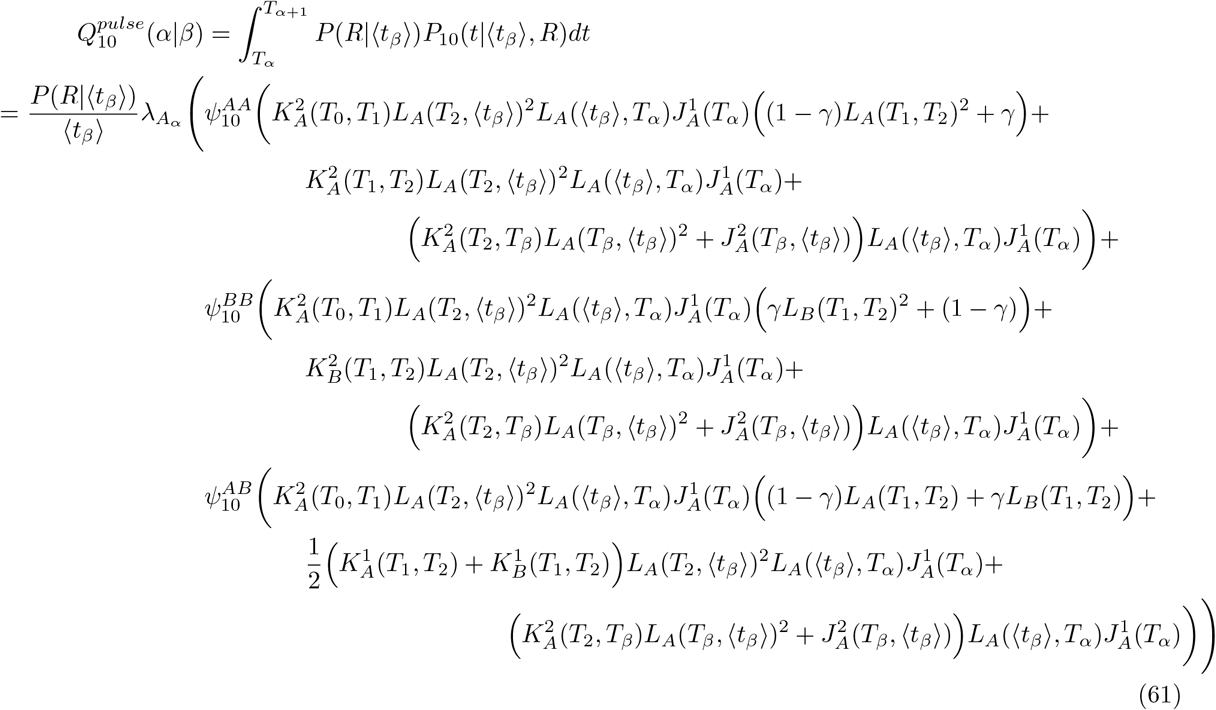

##### 9.3.1.11 Transitions to self

Combining all the different Cases, we can write

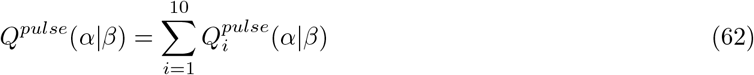

then use the law of total probability to describe the probability that a hidden state transitions to itself (i.e. the probability that no recombination occurred, or after a recombination the lineage coalesced with itself):

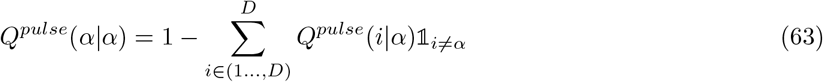

#### 9.3.2 Demonstration of correctness

To validate the transition probabilities for a *pulse* structured model under the SMC model, we compare our theory against simulations using *msprime* [100]. In this section, we denote empirical summary statistics from the simulated data with a circumflex and we omit the *pulse* superscript and the function’s demographic parameters.

With a specific demographic model (defined by parameters ***λ***_*A*_, ***λ***_*B*_, *γ, T*_1_, *T*_2_, *D, ω, T*_*max*_), and simulating with *msprime* we can:

1. Record the sequence of marginal coalescence events **Z** from the ARG. Denote its length by *N*
2. Discretise the marginal coalescence times into the HMM’s corresponding time states **T**, which is defined by *D, ω, T*_*max*_: 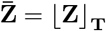
3. Treating 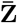 as a Markov chain, we can calculate a matrix which counts the number of transitions from state *β* to *α*

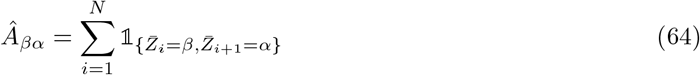

where 1_*E*_ is 1 if event *E* is true and 0 otherwise.
4. From the matrix of counts we can calculate the MLE of the transition matrix for sequence 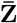

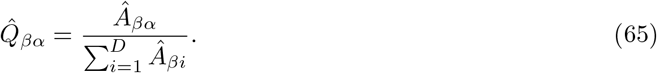

From *Â* we can calculate the proportion of time spent in each time state with

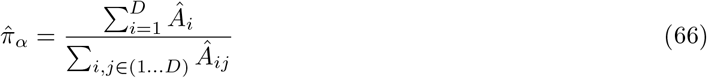

We can also compare 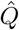 with the probabilities given by the theoretical matrix *Q*.

With an arbitrary demographic model

**Model 9.1** *pulse structured model*

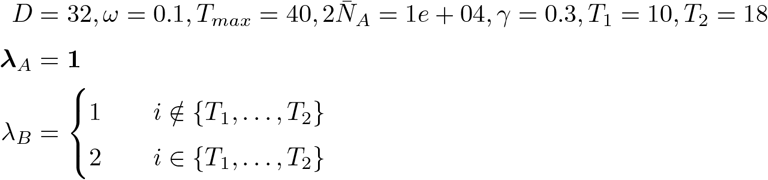

We simulate ∼2e+7 transitions under the Hudson [29, 101] and SMC [30, 31] model with *msprime* [100] and compare this to *cobraa*’s theoretical transition matrix *Q*^*pulse*^ (we use Marjoram and Wall’s SMC model for simulation, which *msprime* labels “SMCprime”) . When discretising time, we can either take the expected coalescence time in each interval, ⟨*t*_*β*_⟩, or the midpoint between boundaries. We calculate the difference between the median simulated coalescence time in each time window and either the expected coalescence time or midpoint in Figure S24a, observing that each gets increasingly inaccurate as the time windows get larger in ancient time.

In Figure S24b and S24c we plot the proportion of time spent in each state for a *pulse* structured model, simulated with Hudson 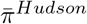 and SMC 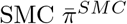, against the theoretical stationary distribution *π* as described in section 9.3.1. These indicate that the Hudson and SMC simulation model are almost indistinguishable from each other, demonstrating the accuracy of the SMC [98]. Moreover the theoretical predictions from *cobraa* are practically identical to the Hudson or SMC simulations except in the most ancient time states where there are larger residuals. These larger residuals are a consequence of the discretisation of time and not a limitation of the theoretical predictions, as the sign of the residuals in ancient time states flips between S24b and S24c.

**Supplementary Figure 24:**
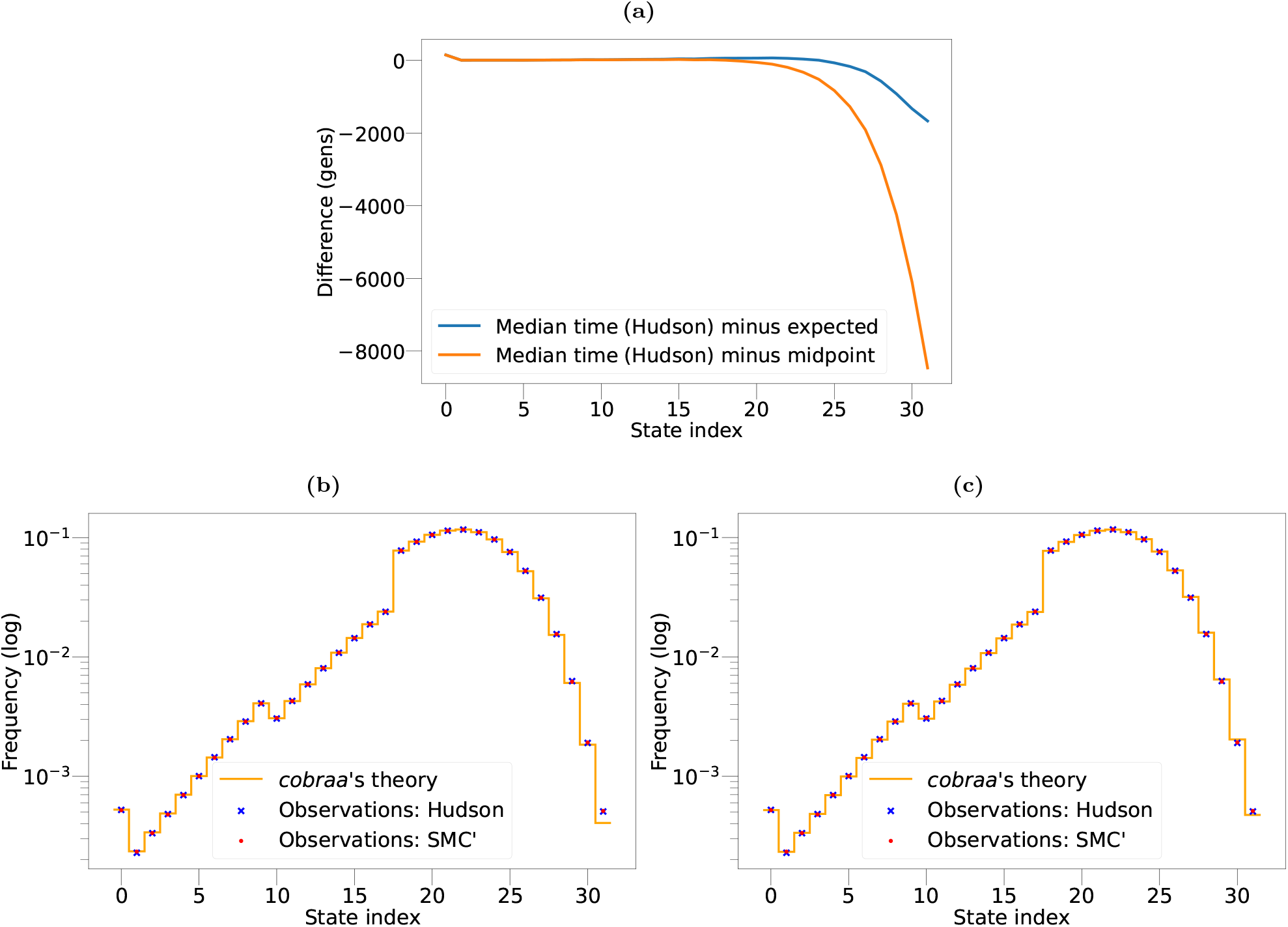
**a)** Difference between the expected coalescence time ⟨*t*_*β*_⟩ (blue) or the midpoint of each time interval (*τ*_*β*_ + *τ*_*β*+1_)*/*2 (orange) against the median of all simulated coalescence events that occurred in that interval, from Hudson’s model. Each approximation becomes more inaccurate as the size of the time interval boundary gets larger. **b)** and **c)** The (log) proportion of time spent in each state, according to a simulation with the Hudson (blue star) and SMC (red point) model, with the theoretical proportion from *cobraa* (orange line) as described by *π*(*t*). (b) Uses the expected coalescence time in each interval, where as (c) uses the midpoint of each interval. We use “SMC” to denote Marjoram and Wall’s modified SMC model, which *msprime* labels as “SMCprime”.

In Figure S25 we visualise the relative difference between *cobraa*’s theoretical transition matrix and the MLE of the transition matrix from a simulation under Hudson 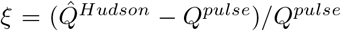 within ±5% variation. For ancient transitions we see larger residuals, though again these are a consequence of time discritisation, as seen from changing between using the expected coalescence time (Figure S25a) or the midpoint (Figure 25b). The remaining residuals are consistent with sampling noise. Furthermore we can visualise a particular row of *ξ*, to see how 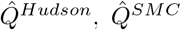 and *Q* differ. Setting *β* = 15 as a time state in the structured period, we visualise these distributions in Figure S25c. It appears that 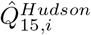 *Hudson* and 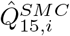 are virtually indistinguishable from each other, except from particularly recent or ancient time states. However, the number of transitions from these states are low and hence the MLE estimates may not be especially reliable for example, at state 31 there are only ∼36 transitions recorded. This possibly explains

**Supplementary Figure 25:**
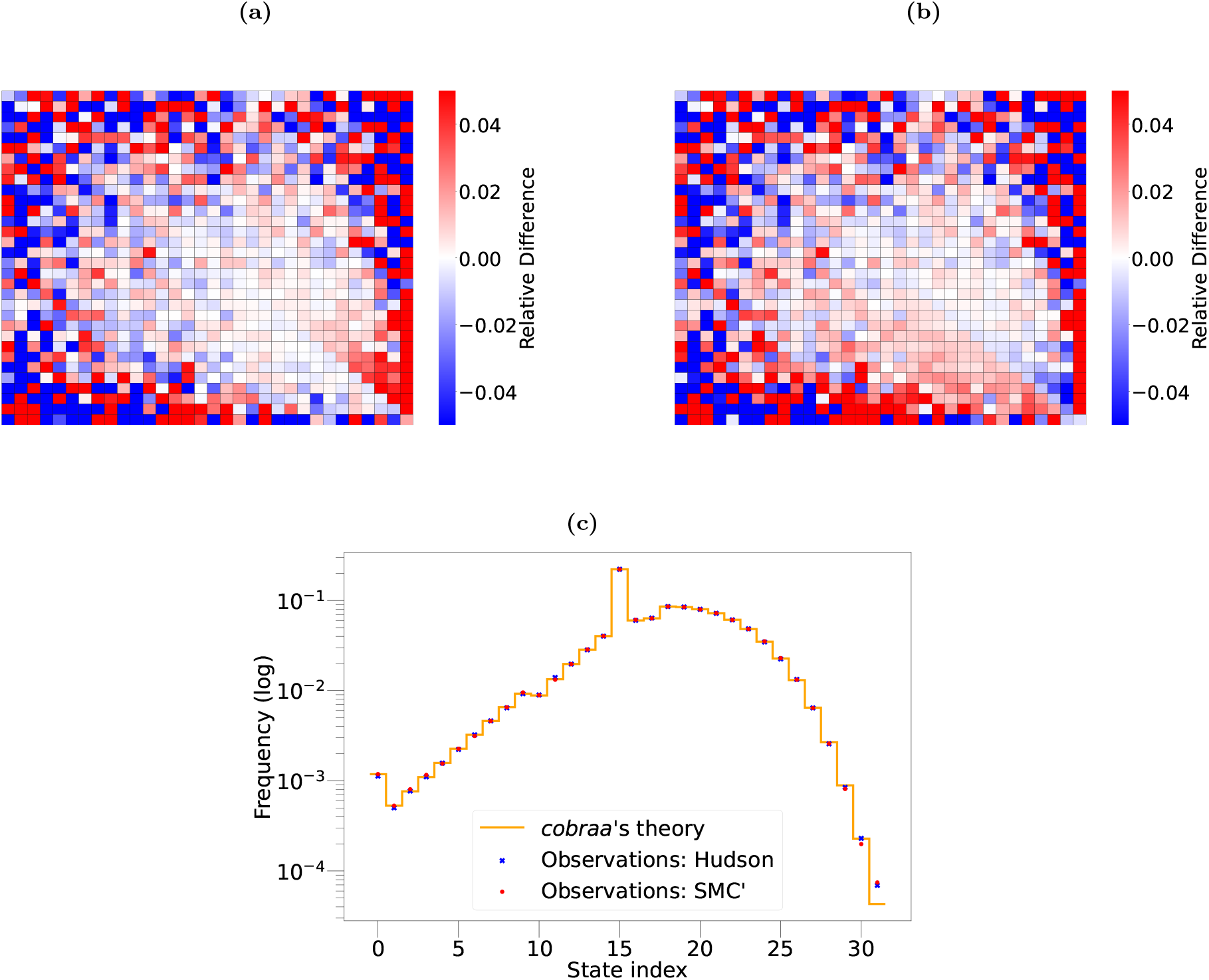
Relative difference between the MLE and theoretical transition matrix 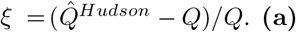 **(a)** Uses the expected coalescence time and **(b)** uses the midpoint of each discrete time window. **(c)** The (log) probability of transition probabilities given *β* = 15 (row 15 of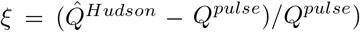), according to a simulation with the Hudson (blue star) and SMC (red point) model, with *cobraa*’s theoretical prediction (orange line) given by *Q*^*pulse*^.

the variance between the Hudson and SMC model. Qualitatively, the theoretical rate is extremely similar to both simulated distributions, except in ancient states which are again explainable by the inaccuracy of ⟨*t*_*β*_⟩ for increasingly large time inteval boundaries.

Finally, we can perform a statistical hypothesis test to see if there is a significant difference between the theory and simulations. Using a Chi-squared test for each row *α*, we compute the test statistic with

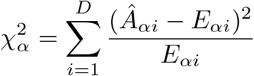

where 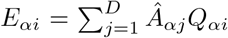 and set the degrees of freedom to *D* − 1. We do not reject the null hypothesis that *Q*_*α*_ provides a suitably good fit to the simulated data, because none of the p-values for the Hudson or SMC model exceed a 5% level of significance.

Figures S24, S25, and the statistical hypothesis testing suggests that our theory of marginal transitions of the *pulse* migration model under the SMC is a very good approximation to Hudson’s coalescent with recombination model.

#### 9.3.3 Identifiability of structured model parameters

In the main text, we stipulated that the size of population *B* must be constant, *N*_*B*_. The theory in the previous section allows time variance in the ghost’s population size, *N*_*B*_(*t*), though we found significant identifiability problems in distinguishing between *N*_*A*_(*t*) and *N*_*B*_(*t*), as discussed below.

##### 9.3.3.1 EM analysis

We test how well the population size parameters and admixture fraction can be inferred by *cobraa*, provided that the split times are known. The ability of a coalescent HMM to perform accurate inference is dependent on the ratio of mutation rate to recombination rate *µ/r*. Initially, to test *cobraa* under a relatively easy scenario, we set *µ/r* = 10 and simulated 10 replicates of 3Gb of a structured model with changes in *N*_*A*_(*t*) or *N*_*B*_(*t*), *T*_1_ = 250ky, *T*_2_=1My, and *γ* = 0.4. We run *cobraa* until convergence (defined as the change in log-likelihood being less than one, *ψ*_ℒ_ *<* 1, in subsequent iterations of the EM algorithm) on 10 replicates, and plot the inference in Figure S26. The simulated *N*_*A*_(*t*) and *N*_*B*_(*t*) are shown in black and gold, respectively, with the inferred *N*_*A*_(*t*) and *N*_*B*_(*t*) shown in red and blue, respectively. The simulated split and rejoin times are indicated by the vertical, green, dashed lines. In Figure S26a population *A* doubles in size in the structured period whilst *B* stays constant, and vice-a-versa in Figure S26b.

We see that the split fraction is consistently estimated with reasonable accuracy, but the inferred *N*_*A*_(*t*) and *N*_*B*_(*t*) parameters are significantly biased. It seems that *cobraa* can not distinguish between changes in *N*_*A*_(*t*) or *N*_*B*_(*t*), and often one is overestimated at the expense of the other being underestimated. To examine this further, we investigated the likelihood surface.

##### 9.3.3.2 Likelihood surface

We consider a structured model with constant size in *A* and *B*, an admixture fraction of 40%, and a separation time of ∼800ky (with *D* = 32 time intervals, this corresponds to a split and rejoin at the 18th and 10th discrete time interval, respectively). Using *msprime* we simulate under Hudson’s model [29, 101] and record the sequence of marginal coalescence trees **Z**. We discretise this sequence of continuous coalescence times into the time windows used by *cobraa* (equation (9)), 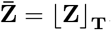 Treating 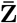 as a Markov chain, we can then calculate the log-likelihood of the sequence from *cobraa* under a particular set of parameters 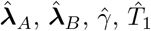 and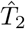, by performing

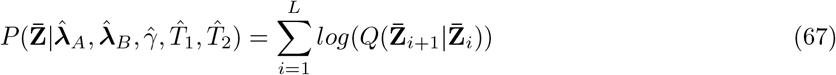

where *Q* is the *pulse* transition matrix (see section 9.3.1) defined by 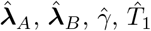 and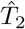 .

In an unconstrained structured model, there are *D* + (*T*_2_ − *T*_1_) + 3 parameters; *D* for ***λ***_*A*_, (*T*_1_ − *T*_2_) for ***λ***_*B*_, and one for each *γ, T*_1_ and *T*_2_. The resulting likelihood function of this unconstrained model therefore is of high dimensionality, making it computationally intractable to simulate under a wide range of values and not amenable to a simple two or three dimensional visualisation. However, as we are interested in the unidentifiability of the effective size parameters for populations *A* and *B* within the structured period (*T*_1_ ≤ *t < T*_2_), we can reduce the dimensionality to capture the essence of the problem by setting 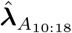 and 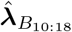 each at one value. Outside the structured period, we fix the effective population size parameters of population *A* and the split/rejoin times at their simulated value, 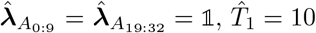 and 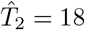. We then let 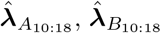 and 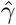 vary over over a grid of values and calculate the likelihood of the sequence under these parameters (equation (67)).

**Supplementary Figure 26:**
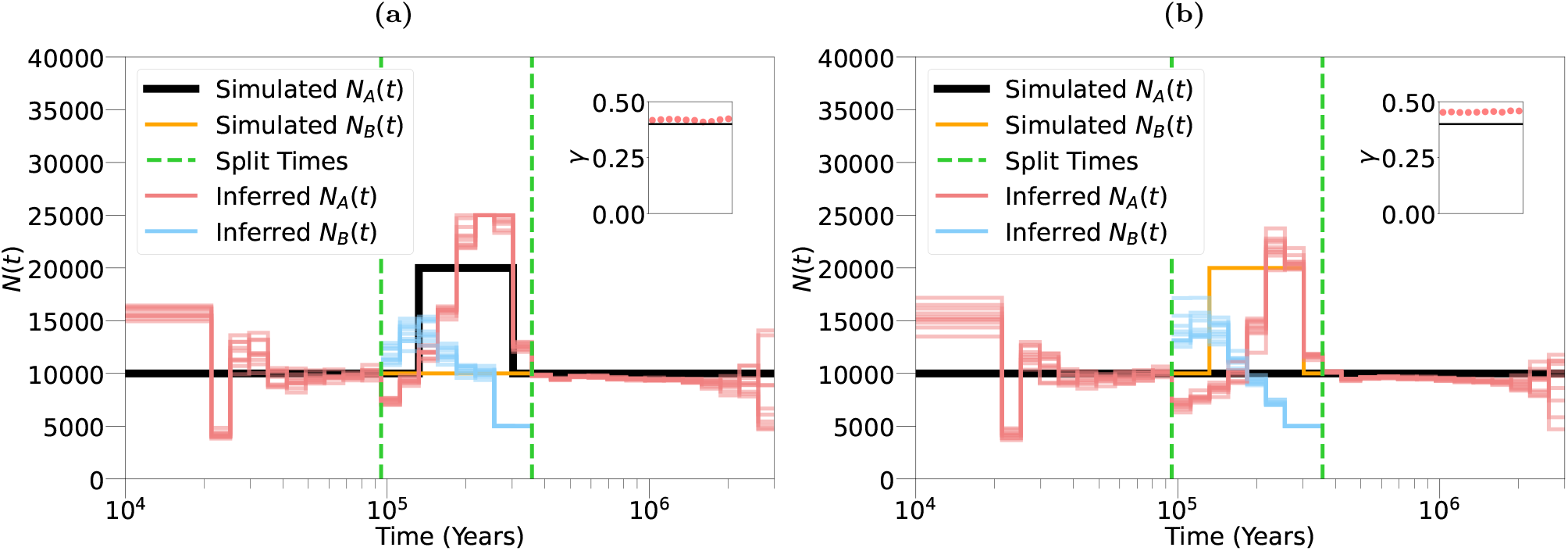
Inference of population size changes for both populations *A* and *B*, and the admixture fraction. Simulated population sizes of population *A* and *B* are shown by the solid red and blue lines respectively. The inferred population sizes of *A* and *B* are shown by the light red and light blue lines respectively. The vertical, dashed green lines indicate the simulated split and rejoin time of the model, and the inferred *γ* is detailed in the inset. Each plot shows a structured model with 40% admixture fraction and changes in (**(a)**) *N*_*A*_(*t*) or **(b)** *N*_*B*_(*t*) in the structured period.

Three visualisations of the three-dimensional likelihood surface are shown in Figure S27. In Figure S27a, a plane of the surface is shown when 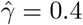 is fixed at its simulated value, to see how the likelihood changes as 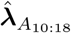 and 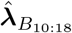 vary. In Figure S27b and Figure S27c, 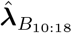 and 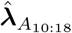 respectively are fixed at their simulated values, to get an idea of how the surface varies with 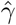 In each, the red cross indicates the simulated value of the non-fixed parameters. Figure S27a demonstrates that there are numerous points in the space of 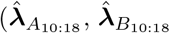) values that achieve a likelihood equally as high as the simulated values, seemingly by over-compensating one and under-compensating the other. In addition to these maxima, it is apparent that there is a whole region of high likelihood corresponding to this inverse relationship between 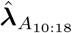 and 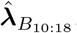. Specifically, this demonstrates that the series of marginal coalescence events 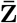 for this simple structured model can be equally well explained by model that for example has a 22% increase in the effective population size of *A* in the structured period, and 27% decrease in *B*, compared to its simulated values.

Generally speaking, if and only if ***λ***_*A*_ and ***λ***_*B*_ are equal everywhere in the structured period, then with respect to *γ* the model is symmetric about *γ* = 0.5. This is because population *A* and *B* are exchangeable in this period if we set *γ*_*B,A*_ = 1 − *γ*_*A,B*_. We see this reflected in Figures S27b and S27c, from the dual likelihood peaks at 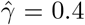 and 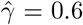 and the fact that the surface in Figure S27c is a vertical mirror of

**Supplementary Figure 27:**
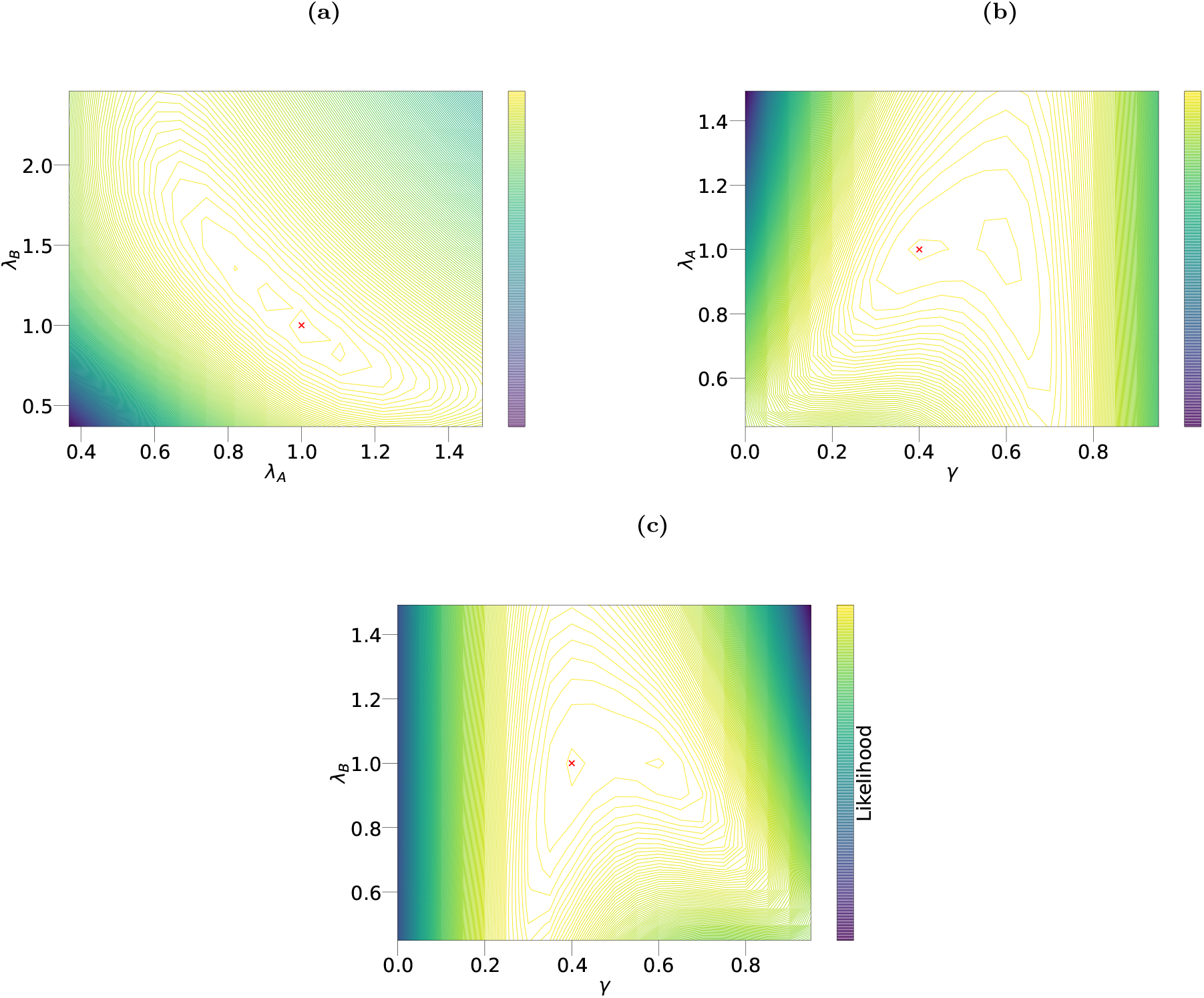
Three contour plots from the likelihood surface for a structured model. The colourbar indicates the log-likelihood, and the red cross indicates the simulated values. Shown in **(a)** is variation between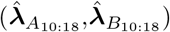 with 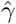 fixed at its simulated value; in **(b)** is variation between 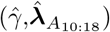 with 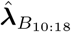 fixed at its simulated value; in **(c)** is variation between 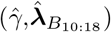 with 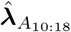 fixed at its simulated value.

S27b. From S27b, it appears that at the simulated split fraction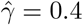 m, the likelihood function is sensitive to the value of 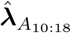, due to the sharp drops seen around its simulated value, though at the symmetric split fraction 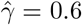 it appears to be less so, as at this axis the likelihood decreases at a slower rate. Note that the reverse is true in S27c: at the simulated split fraction 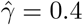, the likelihood function is not as sensitive to 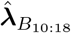 as at the symmetric split fraction 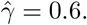

We note that this analysis was performed on the sequence of known marginal coalescence times 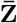 which we treat as a Markov chain, from which it is straightforward to calculate the likelihood (equation (67)). In practise, we do not know the coalescence times 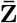 along the genome, so with a sequence of observations **X** we formulate a HMM and calculate the likelihood of the data *P* (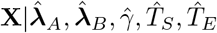) with dynamic programming. Only knowing the observed sequence **X** will necessarily flatten the peaks of the likelihood surface, due to the uncertainty of 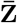 which must be probabilistically inferred, making optimising over the parameter space even harder in reality. We also emphasise that this experiment was performed with the effective population size changes for population *A* and *B* each set to one value in the structured period. In reality, we would hope for more freedom, but this stipulation allowed us to demonstrate the difficulty in navigating the parameter space even under simplified conditions.

In summary, Figure S27 demonstrates the irregularity of the likelihood surface for a *pulse* structured model. Even with a simple evolutionary history and constraining the effective population size parameters to an equal value in the structured period, there appears to be multiple local maxima and regions of high likelihood that do not correspond to the simulated parameters. This indicates that maximum likelihood inference for a model that searches for ***λ***_*A*_, ***λ***_*B*_, and *γ* may be insufficient in ascertaining the simulated parameter values, due to the sensitivity of the optimisation’s starting guess and the difficulty in navigating the parameter space. This justifies our requirement that population *B* is of constant size.

### 9.4 *cobraa-path* ‘s HMM

The hidden states of *cobraa* are coalescence time windows, meaning that when we calculate the probability of transitioning we had to include all the possible paths that a pair of lineages could have taken through the *pulse* model. This is unnatural, and condenses a lot of information into one hidden state. Instead, we can expand the HMM of *cobraa* such that the hidden state represents not just the coalescence time, but also the ancestral path taken through the two populations before the two sampled lineages coalesce. If the coalescence time occurs more recently than the admixture event, then the only possibility is that both lineages coalesced in population *A*, which we denote as *AA*. If the coalescence occurred in the structured period, then there are two possible paths, *AA* and *BB*. If the coalescence occurred more anciently than the population divergence, then there are three possible paths, *AA, BB* and *AB*. An illustration is given in Figure S10.

We call this new HMM *cobraa-path*. The hidden states are then a tuple (*t, c*) where *c* ∈ (*AA, BB, AB*) is the ancestral path taken by the two lineages which coalesce at time *t*. The emission probabilities are similar to *cobraa*’s, though given *t* are repeated across different values of *c*. For example given a coalescence of *t* that is more ancient than population divergence, the probability of observing a mutation is the same for each *c* ∈ *AA, BB, AB*. We also note that if *γ* = 0 then *cobraa-path* reduces to standard PSMC.

The transition probabilities of *cobraa-path* follow from *cobraa*. Let *Q*_*X*_ (*α, j*|*β, k*) = *P*_*X*_ (*z*_*i*+1_ = *T*_*α*_, *c*_*i*+1_ = *j*|*z*_*i*_ = *T*_*β*_, *c*_*i*_ = *k*). Using Case 3 as an illustrative example, we can decompose equation (32) into its constituent parts to rapidly obtain:

**Supplementary Figure 28:**
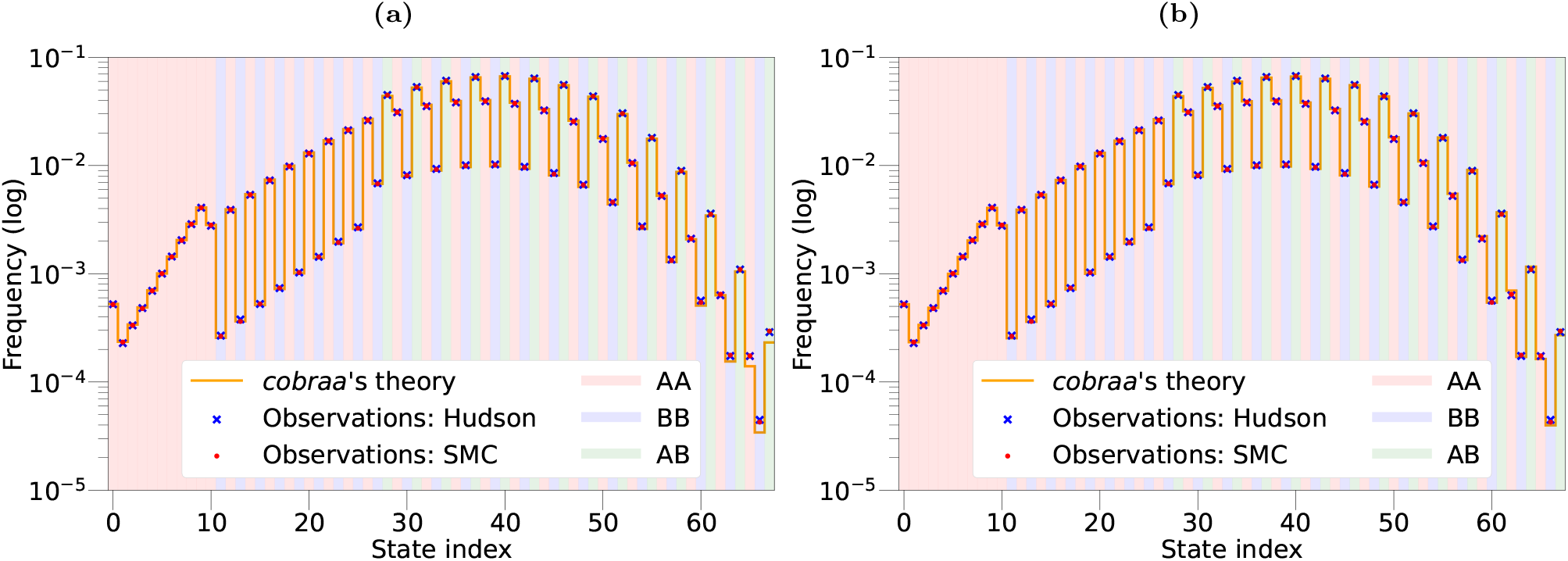
The (log) proportion of time spent in each state, according to a simulation with the Hudson (blue star) and SMC (red point) model, with the theoretical proportion from *cobraa-path* (orange line). **(a)** Using the expected coalescence time in each interval **(b)** Using the midpoint of each interval. The shading represents which ancestral path the hidden state corresponds to, with red being *AA*, blue being *BB* and green being *AB*.

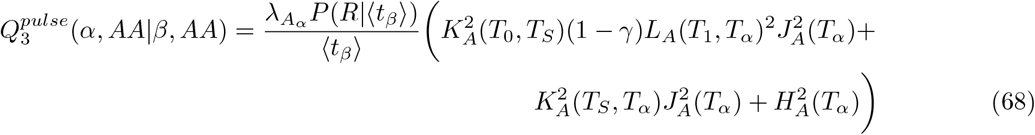

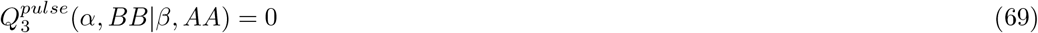

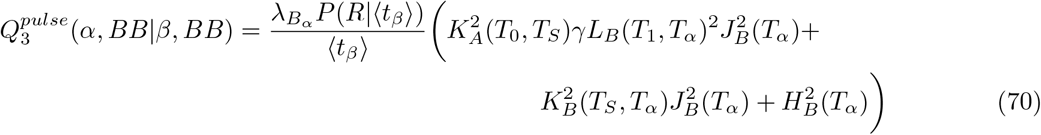

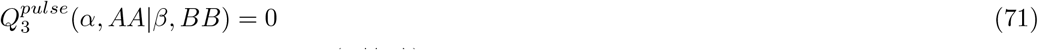

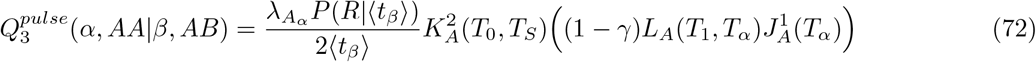

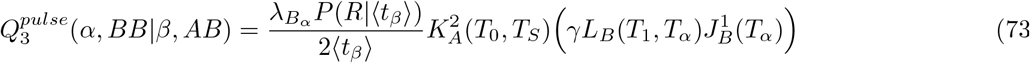

All the other cases follow similarly.

#### 9.4.1 Demonstration of correctness

We simulate marginal transition data from msprime, and record the appropriate lineage path and coalescence time information at each position, using both the Hudson and SMC model. We then compare the observed proportion of time spent in each state to *cobraa-path*’s theory in Figure S28, which indicates the theory is a very good fit to the data.

## Footnotes

1 We use the word “floating” to describe a lineage that has undergone recombination but is yet uncoalesced; we use “solid” to denote the remaining unrecombined lineages

## Notes

### Competing Interest Statement

The authors have declared no competing interest.

